# Underwater caves preserve biochemical composition and histological structure in modern and fossil mammalian bones

**DOI:** 10.64898/2026.07.14.738352

**Authors:** M. M. Walker, J.J. Miszkiewicz, J.M. Rowe, C.D. Matheson, J Vongsvivut, N.A. Sims, J Louys

## Abstract

Biochemical and microstructural bone degradation (diagenesis) occurs after death and is environment dependent. Existing macroscopic and histological research suggests bones submerged in water degrade differently to those in dry contexts. However, little is known about how submerged and dry degradation processes affect skeletal remains in caves, which have fundamentally different decomposition conditions to open environments. This study combines histology with laboratory-based and synchrotron-sourced Fourier-transform infrared microspectroscopy (FTIRM) to characterise bone microanatomical and biochemical degradation in bones from underwater (wet) and dry cave environments. We investigate fossil and historic mammal (ovicaprids, macropodids) bone diagenesis from Mount Gambier, South Australia, intra-skeletally and between depositional conditions. FTIRM analysis revealed greater collagen-associated amide signatures in historic wet bones than dry counterparts, while fossils showed no detectable signal. Despite biochemical differences, bone histology did not present noticeable differences between specimens, with similar birefringence levels and no radial micro-fractures across the secondary osteon border. Wet and fossil bones also exhibited comparable carbonate content despite differences in organic composition and mineral recrystallisation, suggesting submerged cave environments facilitate preservation of carbonate-bearing mineral phases within relatively closed (trapped) diagenetic systems. These findings demonstrate that underwater cave conditions support fossil preservation through complex and heterogenous organic and mineral diagenetic pathways.

## 1 Introduction

Skeletal remains are often found in forensic, archaeological, and palaeontological underwater contexts [1–6]. However, compared to terrestrial settings, less is known about the processes of bone decay and preservation underwater [5]. Since drowning is a common cause of death, and entrapment underwater has been recorded for modern and ancient humans and animals [7–11], understanding the difference between wet and dry depositional environments is key for archaeological and forensic investigations. In palaeontology, differentiating bone from wet and dry deposition in underwater cave systems may prove useful for reconstructing regional environments [5]. Bone diagenesis, the process of degradation and modification of mineral, organic, and structural content by biological and physiochemical processes after death [12, 13], is strongly influenced by burial or depositional environments [14, 15]. The impact and extent of diagenesis can be measured by assessing bone histological integrity alongside alterations to its crystallinity, collagen quality and quantity, and chemical composition measured through the carbonate-to-phosphate ratio [13, 16–19].

Alterations of bone mineral and organic components during diagenesis can improve their long-term preservation. During fossilisation in dry terrestrial cave and open environments, unstable bioapatite transforms into a more stable mineral phase through ionic substitutions, and dissolution of carbonates, is followed by recrystallisation [20]. The replacement of phosphate and hydroxyl ions with carbonate ions [14, 15, 21, 22] is suggested to be the most distinctive measure of preservation [23]. Under the right chemical conditions [24], such ionic substitutions in diagenesis result in recrystallisation (changes in crystal structure) of the bone mineral, increasing atomic order and size of the bioapatite lattice [25]. These mineral changes are linked to enzyme and chemical hydrolysis of collagen type-1 [12, 18, 26, 27], and its secondary order molecular structures, specifically amide I and amide II [28, 29]. In some conditions, amide can be preserved in palaeontological assemblages [30, 31]. However, there is a general trend in historic and archaeological bone where the proportion and quality of amide reduces with increasing time since deposition, subject to changes in local depositional conditions [15]. Nevertheless, changes to bone biomolecular structures are not consistent across different time periods, depositional environment or climatic conditions [15, 30, 32–34].

In aquatic landscapes, a theoretical model of bone diagenetic change was developed to distinguish early from late diagenesis [35]. Early and late diagenesis are not identified across timescales but are defined by the types of modification to bone. Early diagenesis of bones in water is dominated by enzyme-mediated hydrolysis induced by microbial communities that dissolve and displace discrete pockets of bone, accompanied by chemical hydrolysis and gelatinisation of collagen structure. Ionic exchange of chemical elements between bone mineral and the aquatic or soil depositional environment also begins in early diagenesis [20, 25, 35]. Inorganic reactions and structural changes to the bioapatite lattice continue into the late diagenetic period (long-term phase of fossilisation) [20, 35]. The degree of water exposure and hydrological regimes, however, play key roles in collagen decay as chemical hydrolysis breaks down peptide bonds [26, 36]. Whilst some studies have investigated the impact and timing of early bone diagenesis in water [37, 38], the cumulative modifications observed in late diagenesis have not been studied.

Diagenesis does not occur uniformly in bone [21, 39, 40]. Cortical bone contains two anatomically and compositionally different structural elements: (1) secondary osteons, which are branched cylinders of material surrounding vascular canals, and (2) interstitial bone, which lies between the secondary osteons. Theoretically, with intact (i.e., not fragmented after death) bones, water should first come into contact with bone at the bone exterior surface, then diffuse from this periosteal surface into the cortical bone, or enter via bone foramina and disperse through the blood vessel system. This may vary, particularly if fractures exist that expose endosteal surfaces to water, and if other decomposition and defleshing conditions are present. Higher mineral diagenesis and recrystallisation of the outer sub-periosteal bone region, compared to the inner and middle bone cortices, has been observed in biomolecular studies of bones from terrestrial environments [39]. A similar pattern may be observed for wet environments, mirroring diffusion of water into the cortex from the bone surface. Further, possibly different timings of chemical collagen hydrolysis in secondary osteons and interstitial bone, could lead to creation of radial microfractures across the border between these two structures (cement lines) to relieve water pressure and stress at that site (Figure 1) [41–43].

**Figure 1.**
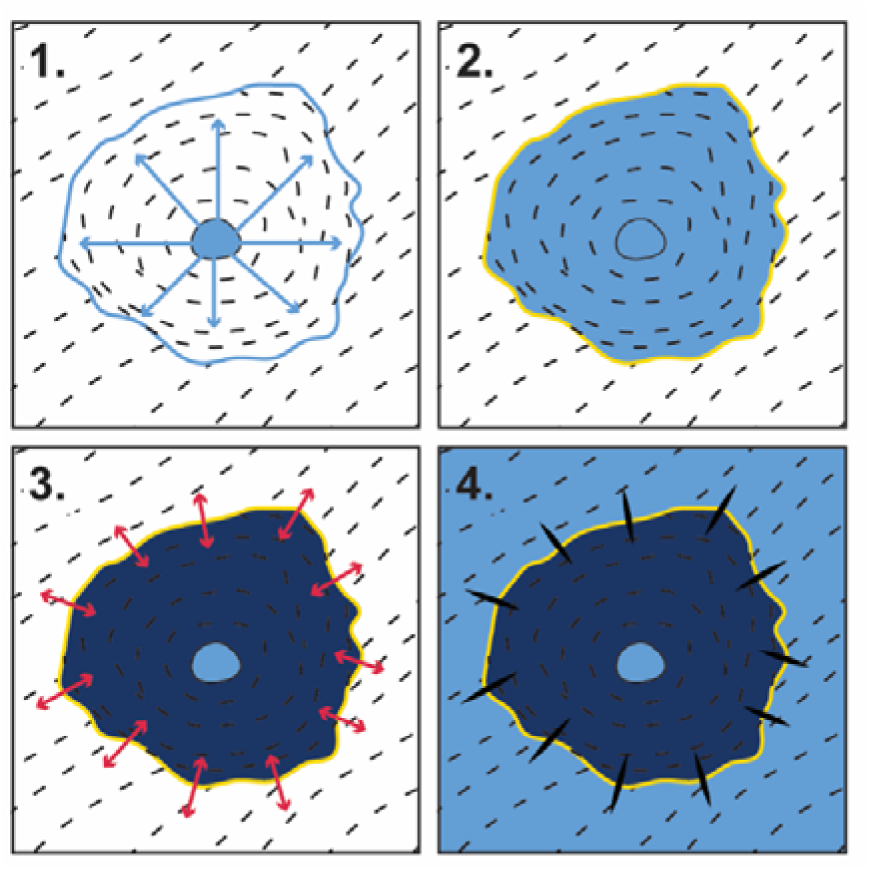
Theoretical diagenesis within a secondary osteon in submerged bone samples. 1) water moves (blue arrows) through the central secondary osteon blood canal (central circular structure) and diffuses across the osteonal wall (light blue shading). 2) Water fills the secondary osteon and is temporarily trapped by cement lines (yellow boundary). 3) Collagen hydrolysis in the secondary osteon results in a pocket of swollen bone (dark blue shading), building pressure on the cement lines (red arrows). 4) Microfractures crack open across the cement line to relieve the pressure, and water fills the surrounding bone tissue.

In underwater environments, current knowledge about the impact of chemical diagenesis on bone is limited, with research focusing on bone histological tunnelling patterns by microbial communities that are frequently, but not consistently, thought to be produced by aquatic micro-organisms [38, 44, 45]. In diagenesis over two years in 72 femora and tibiae of domesticated sheep, Guareschi et al. [46] showed more bone diagenetic porosity and less collagen content in aquatic environments (saltwater, freshwater, artificial seawater) compared to terrestrial conditions. Pig femora (n=52) deposited in simulated Australian aquatic environments for 12 months showed that the water chemistry in weir and river environments had the most impact on increasing diagenetic porosity compared to estuarine or marine environments [47]. Under longer time scales, Chadefaux et al. [48] found lower collagen alpha-helix and higher unordered structure (random coils) content in Neolithic mammalian bone (n=4; wild boar humerus and unidentifiable mammals) from soils with fluctuating hydrology, compared to those from sub-aquatic zones (n=2; bovine radius and deer ‘flat bone’) from the Chalain lake site, France. Comparisons between cow femora (n=940) deposited in fresh and brackish rivers, and a saline coastline environment over 18 months, with control samples (n=156) in fresh water tanks, however, showed limited changes to bone crystallinity but lower carbonate content in samples placed in natural environments [49]. In a Brazilian limestone cave setting, a fossil ground sloth vertebra that experienced long-term exposure to dripping cave water, saturated in calcium carbonate, presented with higher crystallinity and lower carbonate, compared to another ground sloth vertebra specimen from a consistently dry cave environment [50]. However, no research has tackled bone diagenesis in underwater cave environments, which differ from open water and dry terrestrial environments in the rate and nature of skeletal element deposition [5]. Studies specifically investigating open system underwater diagenesis are further limited by short experimental timescales, or small archaeological sample sizes.

In this study we aim to improve current understanding of the microstructural and biomolecular changes associated with diagenesis underwater. A historical assemblage of non-human faunal bones from submerged (wet) and dry cave environments is investigated using bone histology and Fourier-transform infrared microspectroscopy (FTIRM). Diagenesis patterns across the bone cortex and in different tissue types are examined in bones from wet environments to test the impact of water filtration starting at bone surfaces (periosteal and endosteal) and from the vascular canals.

We ask the following research questions:

1. Are there any microstructural and biochemical differences in diagenesis between bones from wet and dry cave conditions?
2. Are there any microstructural and biochemical diagenetic differences between different regions of wet bone, specifically the secondary osteon and surrounding interstitial bone?

Based on these, we then explore a palaeontological assemblage from one of the same underwater sites and theorise that early diagenesis will help inform the fossils’ original depositional environment (wet or dry).

## 2 Materials and methods

### 2.1 Sample selection

Non-human faunal bone specimens were collected from the surface of cave floors across two underwater caves systems in Mount Gambier, South Australia: Green Waterhole (5L81; [51] and Gouldens Sinkhole (5L8; [52]. These caves are filled with fresh ground water enriched in calcium carbonate from the surrounding karstic geology [53, 54]. Cave waters are consistently cool (approximately 16°C), chemically neutral (pH 7.4), and have very low hydrological flow rates. Three depositional environment conditions are here defined as historical wet, historical dry, and geological fossil (Figure 2).

**Figure 2.**
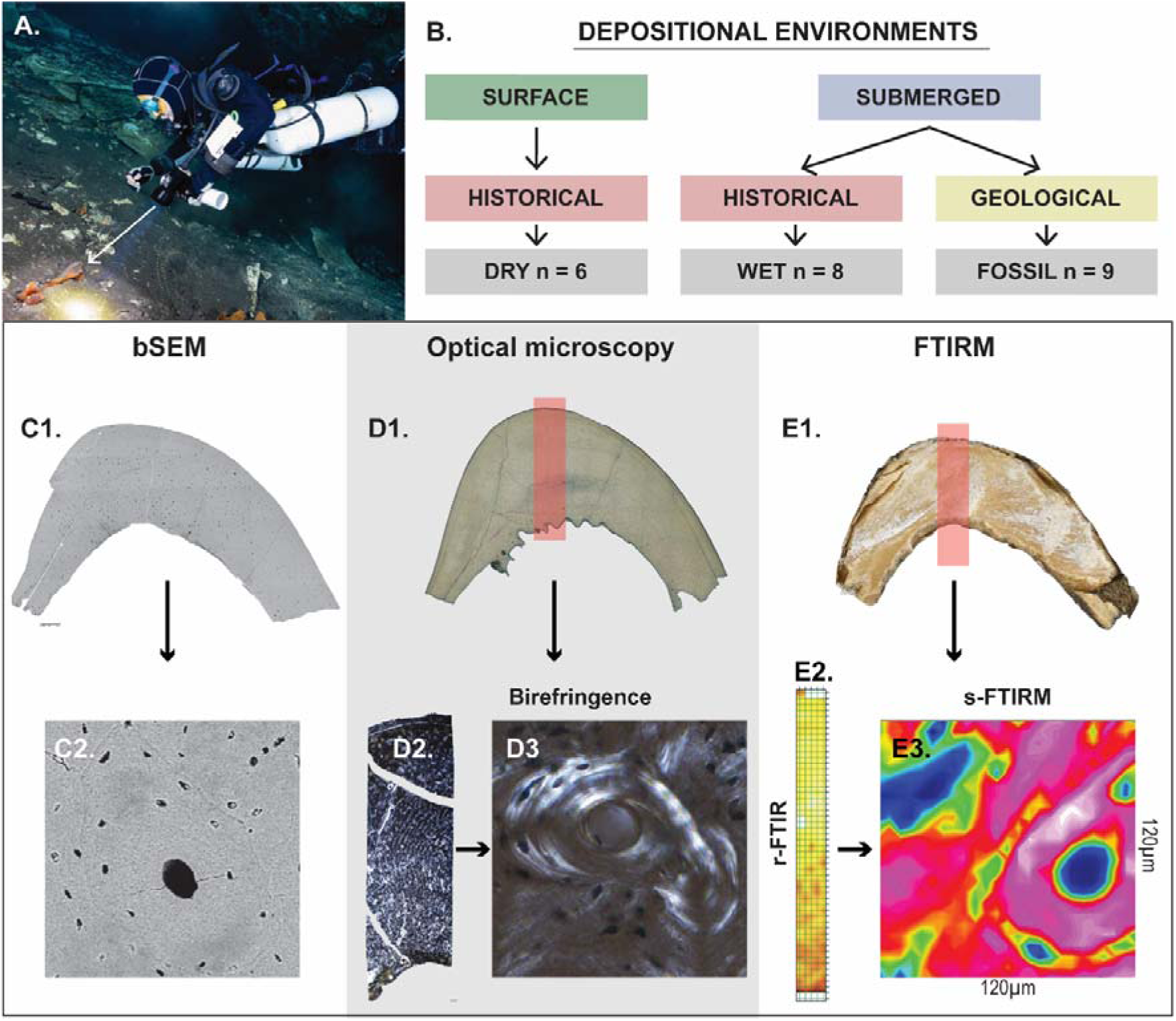
Study design summarising the assemblage depositional environments and methods applied. A. Author Julien Louys collecting bones from the underwater cave Green Waterhole, white arrow indicating a fossil in situ; B summary of depositional environments and their environmental and chronological contexts; C. backscatter scanning electron microscopy (bSEM) cross section (C1) and osteon (C2); D. optical microscopy with full cross section under transmitted light and indicating cortical strip location with red box (D1), cortical strip under polarised light (D2) and secondary osteon under polarised light (D3); E. Fourier transform infrared microscopy (FTIRM) measurements of bone block and cortical strip area indicated by red box (E1), false-colour chemical map of cortical strip acquired using reflectance FTIRM (E2) and false colour chemical map of secondary osteon obtained using synchrotron source FTIRM in ATR-FTIR mode (E3).

We assess n=14 historically dated specimens from ovicaprid limb bones, and *Macropus* limb and rib bones, which were collected in 2023 and selected from underwater (wet; n=8) and dry (n=6) conditions (Table 1). Intact shafts were targeted for analysis but not always available due to the nature of archaeological and palaeontological assemblages (Table 1). Taphonomic histories suggested domesticated faunal remains from Green Waterhole were likely the result of historic refuse dumping of butchered animals [38, 55, 56]. Those from Gouldens Hole did not show butchery marks, and it was inferred that individuals either drowned in the sinkhole or their remains were transported to the site from nearby terrestrial settings shortly after death [38]. Bone surface modifications and microscopic tunnelling patterns in bones from the wet and dry sections of the caves also differed, further suggesting distinct depositional histories between these settings [38].

**Table 1.**
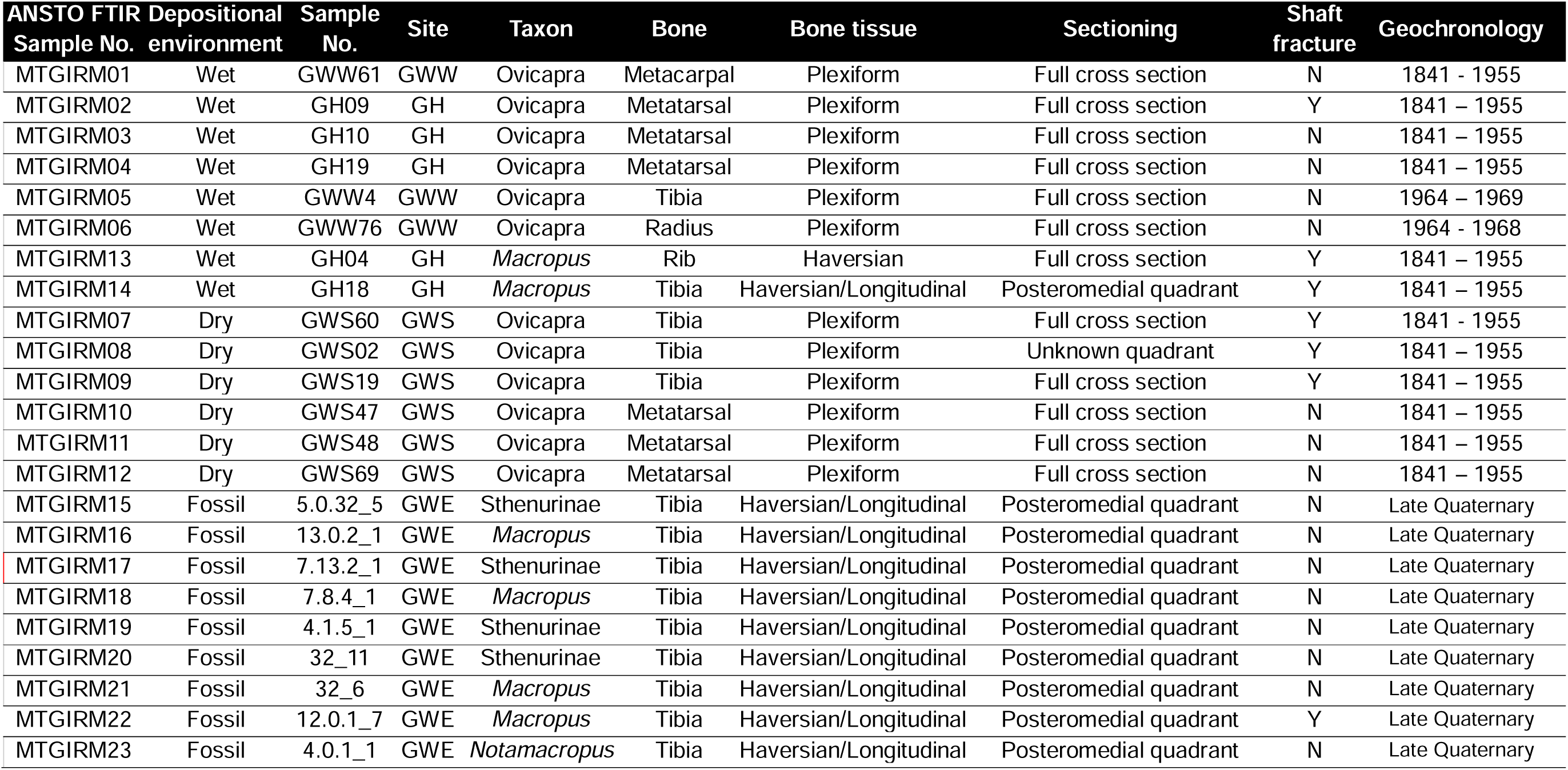
Historic and fossil specimens and associated depositional environments analysed using histology and Fourier Transform infrared microscopy (FTIRM) techniques. Specimens were collected from South Australia: Green Waterhole east lake (GWE), Green Waterhole west lake (GWW), Green Waterhole dry surface (GWS) and Gouldens Hole (GH). Geochronology data are based on results from Walker et al (2026).

| ANSTO FTIR Sample No. | Depositional environment | Sample No. | Site | Taxon | Bone | Bone tissue | Sectioning | Shaft fracture | Geochronology |
| --- | --- | --- | --- | --- | --- | --- | --- | --- | --- |
| MTGIRM01 | Wet | GWW61 | GWW | Ovicapra | Metacarpal | Plexiform | Full cross section | N | 1841 - 1955 |
| MTGIRM02 | Wet | GH09 | GH | Ovicapra | Metatarsal | Plexiform | Full cross section | Y | 1841 – 1955 |
| MTGIRM03 | Wet | GH10 | GH | Ovicapra | Metatarsal | Plexiform | Full cross section | N | 1841 – 1955 |
| MTGIRM04 | Wet | GH19 | GH | Ovicapra | Metatarsal | Plexiform | Full cross section | N | 1841 – 1955 |
| MTGIRM05 | Wet | GWW4 | GWW | Ovicapra | Tibia | Plexiform | Full cross section | N | 1964 – 1969 |
| MTGIRM06 | Wet | GWW76 | GWW | Ovicapra | Radius | Plexiform | Full cross section | N | 1964 - 1968 |
| MTGIRM13 | Wet | GH04 | GH | *Macropus* | Rib | Haversian | Full cross section | Y | 1841 – 1955 |
| MTGIRM14 | Wet | GH18 | GH | *Macropus* | Tibia | Haversian/Longitudinal | Posteromedial quadrant | Y | 1841 – 1955 |
| MTGIRM07 | Dry | GWS60 | GWS | Ovicapra | Tibia | Plexiform | Full cross section | Y | 1841 - 1955 |
| MTGIRM08 | Dry | GWS02 | GWS | Ovicapra | Tibia | Plexiform | Unknown quadrant | Y | 1841 – 1955 |
| MTGIRM09 | Dry | GWS19 | GWS | Ovicapra | Tibia | Plexiform | Full cross section | Y | 1841 – 1955 |
| MTGIRM10 | Dry | GWS47 | GWS | Ovicapra | Metatarsal | Plexiform | Full cross section | N | 1841 – 1955 |
| MTGIRM11 | Dry | GWS48 | GWS | Ovicapra | Metatarsal | Plexiform | Full cross section | N | 1841 – 1955 |
| MTGIRM12 | Dry | GWS69 | GWS | Ovicapra | Metatarsal | Plexiform | Full cross section | N | 1841 – 1955 |
| MTGIRM15 | Fossil | 5.0.32_5 | GWE | Sthenurinae | Tibia | Haversian/Longitudinal | Posteromedial quadrant | N | Late Quaternary |
| MTGIRM16 | Fossil | 13.0.2_1 | GWE | *Macropus* | Tibia | Haversian/Longitudinal | Posteromedial quadrant | N | Late Quaternary |
| MTGIRM17 | Fossil | 7.13.2_1 | GWE | Sthenurinae | Tibia | Haversian/Longitudinal | Posteromedial quadrant | N | Late Quaternary |
| MTGIRM18 | Fossil | 7.8.4_1 | GWE | *Macropus* | Tibia | Haversian/Longitudinal | Posteromedial quadrant | N | Late Quaternary |
| MTGIRM19 | Fossil | 4.1.5_1 | GWE | Sthenurinae | Tibia | Haversian/Longitudinal | Posteromedial quadrant | N | Late Quaternary |
| MTGIRM20 | Fossil | 32_11 | GWE | Sthenurinae | Tibia | Haversian/Longitudinal | Posteromedial quadrant | N | Late Quaternary |
| MTGIRM21 | Fossil | 32_6 | GWE | *Macropus* | Tibia | Haversian/Longitudinal | Posteromedial quadrant | N | Late Quaternary |
| MTGIRM22 | Fossil | 12.0.1_7 | GWE | *Macropus* | Tibia | Haversian/Longitudinal | Posteromedial quadrant | Y | Late Quaternary |
| MTGIRM23 | Fossil | 4.0.1_1 | GWE | *Notamacropus* | Tibia | Haversian/Longitudinal | Posteromedial quadrant | N | Late Quaternary |

Fossil Macropodidae are represented in our study by nine tibiae. These were collected in 1979 from Green Waterhole east lake (Table 1) [57]. Preliminary dates suggest the deposit is 60 ka [58], however, there may be considerable time-averaging of the fossil assemblage across the cave site, such that the nine bones examined here may not be coeval.

Sampling permissions for histological and FTIRM analysis of fossil specimens were provided by Flinders University, Palaeontology Department. Collection and analysis permissions for historic samples were provided by the Department for Environment and Water (DEW), the South Australian Heritage Council (Permit No. 0001/23), and Cave Divers Association of Australia.

### 2.2 Sample preparation

Standard methods for preparing bones and fossils for histology, backscatter scanning electron microscopy (bSEM) and FTIRM were followed [38, 59, 60]. Cortical bone mid-shafts of load bearing long bones, and a single rib, were targeted to ensure secondary osteon presence [61]. Unless otherwise noted (Table 1) full cross-sections were extracted from the modern bones, and a roughly 1 x 2 cm section was extracted from the posteromedial quadrant of tibial midshaft in the fossils [62]. This was done using a Dremel^®^ Variable-Speed Rotary Tool with Dremel^®^ cut-off wheel. The complete cross-sections were approximately 1 cm thick and either extracted from an exposed shaft (where bone was already fragmented post-mortem) or from the cortical midshaft.

Bone sections were embedded in Buehler epoxy resin (EpoxiCure^TM^). From these blocks, further sections were cut using an Allied Techcut 4TM low speed saw and 150 µm Kemet Diamond Wheel. These were separated for histology, FTIRM and bSEM (Figure 2). Samples for FTIRM and bSEM were ground to an even surface using 600/P1200 grit wet/dry sandpaper and then polished using Al203 powder on a wet polishing pad. Histology samples were ground evenly and polished as above and mounted onto glass microscope slides using Selleys^®^ ultra-clear Araldite epoxy adhesive. An approximately 100 +/- 90 µm thin section was developed through manual grinding using a series of abrasive grinding pads and polished. A temporary coverslip mounted with glycerine was applied for imaging.

### 2.3 Microscopy

Histotaphonomy (the study of bone diagenesis using microscopic methods, Bell, 2011) of the fossil samples was analysed under bSEM using a Hitachi TM3030 SEM_EDS model, using BSE mode (standard or reduced charge up) under high charge 15kv voltage. Stepwise images were taken across the bone samples, between 50× and 80× magnification to later generate a composite image of cross sections in Photoshop v26. Additional images were taken at higher magnification to highlight bone structures and taphonomic features of interest. Samples selected for analysis showed high levels of preservation, with at least 80 percent of original histological structures visible. Composite bSEM images were then assessed for presence/absence of radial microfractures across secondary osteons. If present, they were counted using the point and click tool in ImageJ (1.54g). Consistently good preservation allowed for a later cross-comparison between histological structural modifications and FTIRM biochemical alteration (see Section 2.4 for FTIRM methodology).

Histology samples were imaged using an Olympus BX63 high powered microscope under transmitted and polarised light. A ‘strip’ of bone (spanning cortical bone from the endosteal to the periosteal surface) was imaged and automatically stitched under 10× magnification. Birefringence was scored based on prior studies where it is assessed as to whether it is comparable to either fresh bone (1), reduced (0.5) or absent (0) birefringence [60, 64, 65]. Only bone areas with no obvious bacterial diagenesis were assessed to check birefringence in preserved bone structures.

### 2.4 Fourier-transform infrared microscpectroscopy (FTIRM)

Fourier-transform infrared microspectroscopy (FTIRM) was used to characterise both the organic and inorganic components of bone diagenesis. Conventional reflectance (r-)FTIRM measurements were performed using the Griffith Analytical Facility (GAF) at Griffith University (Brisbane, Australia). Synchrotron (s-)FTIRM measurements were also conducted on the Infrared Microspectroscopy (IRM) beamline at the Australian Synchrotron (Victoria, Australia) using macro attenuated total reflectance infrared microspectroscopy (ATR-FTIRM).

#### 2.4.1 Reflectance FTIRM

Complete cortical bone cross section strips, extending from the endosteal to periosteal surface and at least 400 µm wide, were analysed using a Perkin Elmer Spotlight 400 microscope and Spectrum 3 spectrometer in reflectance mode (Figure 3). Background spectra were acquired from a gold (Au) mirror using 8 co-added scans at a spectral resolution of 4 cm^-1^ over the spectral range of 4000-500 cm^-1^. Spectral maps were collected using a 100 µm projected aperture and a 100 µm step interval, resulting in 252-380 measurement points per specimen depending on bone size and sampling width. False coloured chemical maps were generated across the region of the cortex using Spectrum IMAGE software (Perkin Elmer). Reflectance spectra were subsequently subjected to Kramers–Kronig transformation, followed by data tune-up, which includes a baseline correction, using Spectrum IR software (Perkin Elmer). Data were exported into individual spectra files and re-combined in Python for further analysis (see Section 2.4.3 for spectral analysis method).

**Figure 3.**
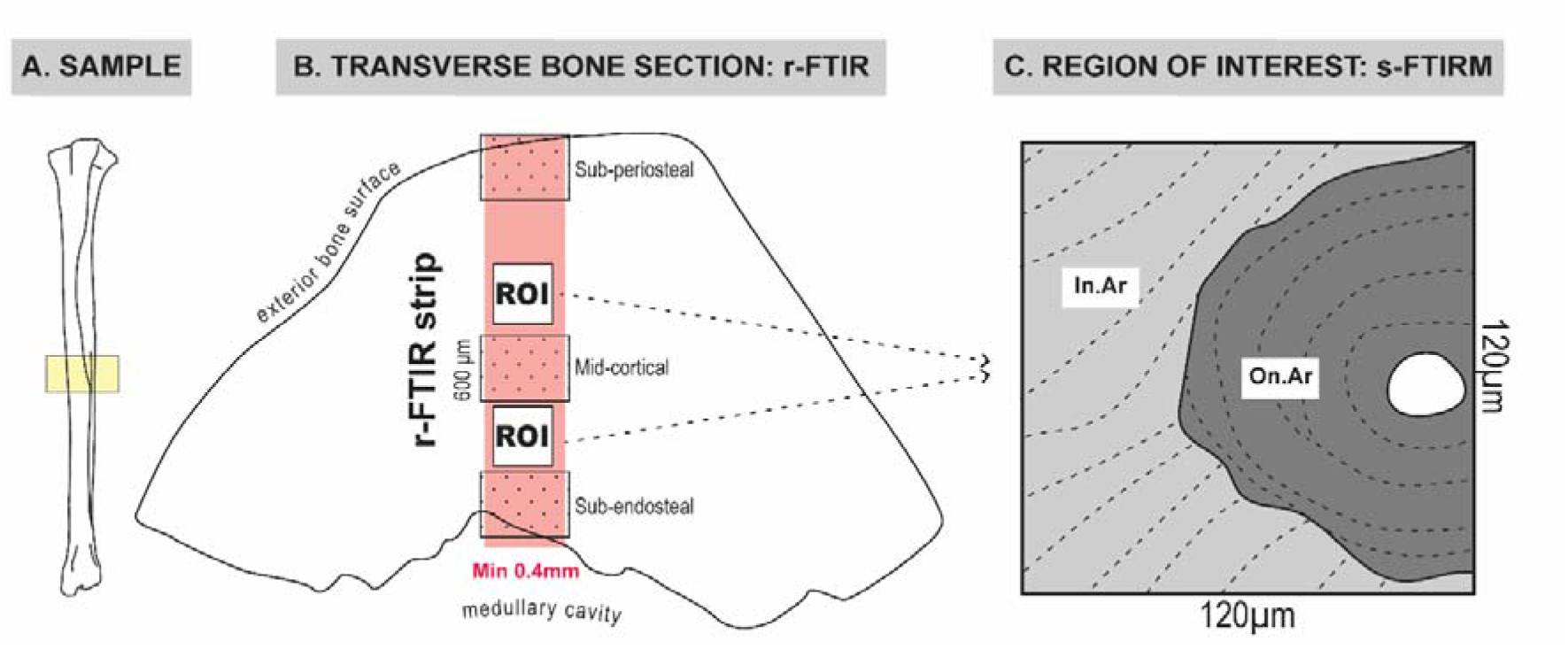
Fourier-transform Infrared Microspectroscopy (FTIRM) sampling strategy. (A) Cortical bone specimens were primarily extracted from the midshaft of load bearing long bones (yellow box). (B) Following preparation as polished blocks, a cortical strip extending from the periosteal to endosteal surface (≥0.4mm wide) was analysed by conventional reflectance (r-FTIRM) (red box). Three cortical regions spanning the width of the strip and 600-700 µm were averaged to generate datasets representing sub-periosteal, mid-cortical, and sub-endosteal cortex. Two secondary osteons were then selected as regions of interest (ROIs) from discrete locations within the same specimens for analysis using synchrotron source (s-)FTIRM in ATR-FTIR mode. (C) Spectral data were extracted from the entire ROI, interstitial bone (light grey) and secondary osteon bone (dark grey) for subsequent analysis.

#### 2.4.2 Synchrotron-FTIRM in macro ATR-FTIR mode

Secondary osteons and their surrounding interstitial bone were analysed using high-resolution s-FTIRM chemical imaging technique, to characterise biochemical variations across secondary osteon cement line (Figure 3). Two secondary osteons were sampled in each bone specimen. One secondary osteon was selected close to the endosteal border, and one from the mid-cortical bone region. A secondary osteon within the sub-periosteal cortex was not selected because this region exhibited microbial degradation and typically contains relatively few secondary osteons in macropod cortical bone [62].

The s-FTIRM experiments were conducted on the IRM beamline at the Australian Synchrotron (Clayton, Victoria, Australia), using a Bruker Vertex 80v spectrometer coupled to a Hyperion 3000 FTIR microscope and a liquid nitrogen-cooled narrow-band mercury cadmium telluride (MCT) detector (Bruker Optik GmbH, Ettlingen, Germany). All the s-FTIRM spectra were recorded over the spectral range of 3800_700 cm^-1^ at a spectral resolution of 4-cm^-1^ with Blackman-Harris 3-Term apodization, Mertz phase correction, and zero-filling factor of 2, using OPUS 8 software suite (Bruker Optik GmbH, Ettlingen, Germany).

High-resolution chemical ‘heat’ maps of secondary osteons were acquired in ATR-FTIR mapping mode, using an in-house developed macro ATR-FTIR accessory incorporating a hemispherical germanium (Ge) ATR crystal with a 250-μm-diameter sensing facet (*n*_Ge_ = 4.0) and a 20× IR objective (NA = 0.60) [59, 66, 67]. Background calibrations were conducted in air using a 3.1 µm aperture at 256 co-added scans. The Ge crystal was then lowered onto the polished bone surface, and high-resolution chemical maps were collected over 120 x 120µm region of interests (ROI) encompassing the selected secondary osteon, using 5 μm step interval and 16 co-added scans per spectrum. Optical microscopic images of each targeted secondary osteon were taken at 4x and 20x magnification before and after crystal-sample contact to verify accurate alignment between the selected optical region and the mapped ATR area.

#### 2.4.3 FTIRM data analysis

The extent of bone diagenesis was assessed using established spectral indices describing mineral composition, crystallinity, and collagen-associated molecular signatures (Table 2). Peak absorbance was determined as the height between selected band maximum and its corresponding baseline, following previously published methods (Figure 3, Figure 4, Supplementary 1) [15, 21, 40].

**Figure 4.**
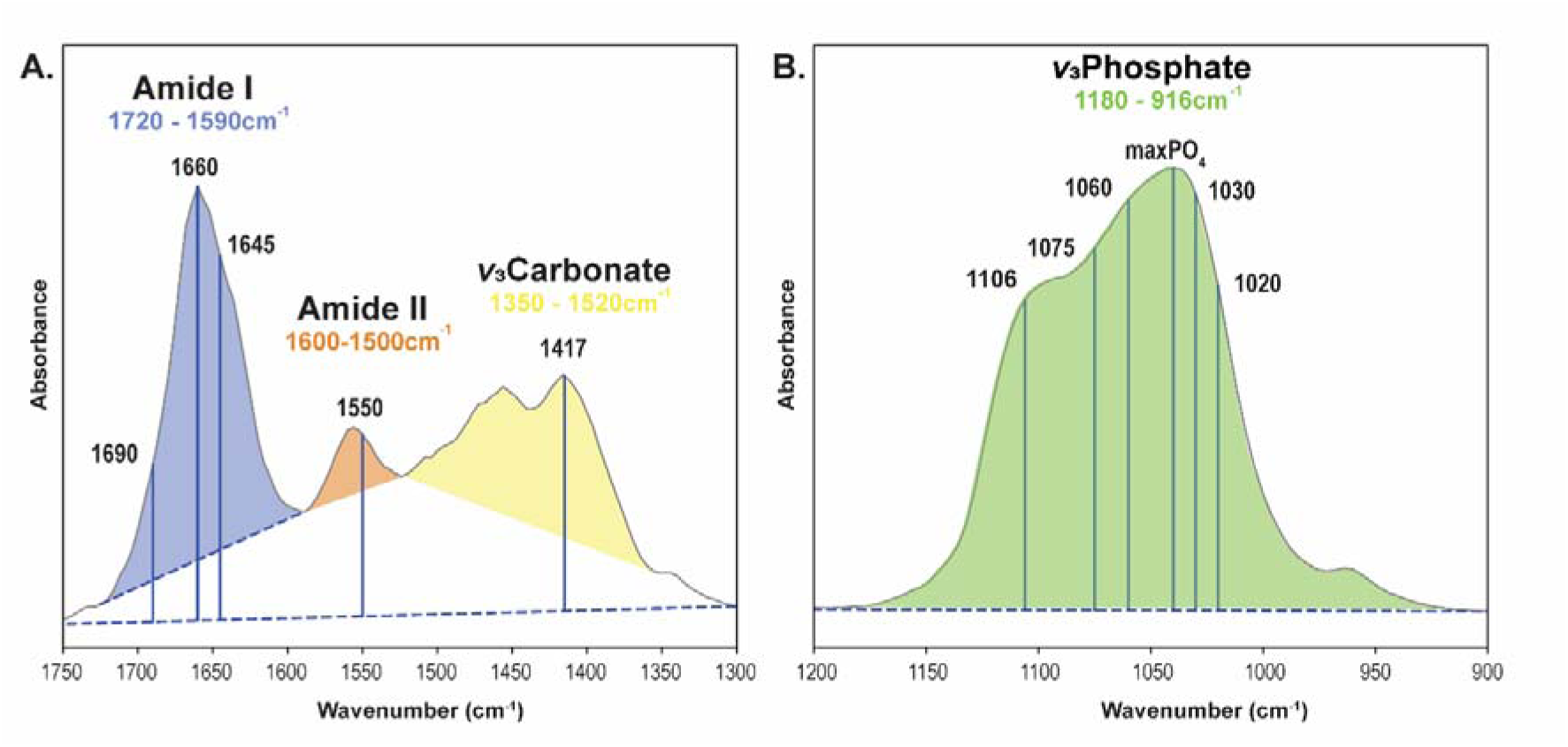
Spectral regions and baseline definitions used for FTIR analysis. (A) Amide I (blue), Amide II (orange), ν_₃_ carbonate (yellow) spectral regions showing the characteristic peak positions and baselines used to calculate collagen-associated and carbonate-related spectral indices. (B) ν□ phosphate (PO_4_; green) spectral region showing the peak positions and baseline used to determine phosphate-related indices, including hydrogen phosphate and crystallinity measures.

**Table 2.** Diagenetic measures analysed based on prior published studies.

| Diagenesis index | Abbreviation | Wavelengths ($\text{cm}^{-1}$) | r-FTIRM baseline maxima and minima wavelengths ($\text{cm}^{-1}$) | s-FTIRM baselines wavelengths ($\text{cm}^{-1}$) | Reference |
| --- | --- | --- | --- | --- | --- |
| **Inorganic biomineral diagenesis** |  |  |  |  |  |
| Carbonate to phosphate | C/P | $I_{1417}/I_{\text{maxPO}_4}$ | max 1700-1800,<br>min 1300-1400 /<br>max 1150-1250,<br>min 850-950 | 1800 - 1300 /<br>1200 - 900 | [40] |
| Hydrogen phosphate to phosphate | $\text{HPO}_4/\text{P}$ | $I_{1106}/I_{\text{maxPO}_4}$ | max 1150-1250,<br>min 850-950 | 1200 - 900 | [40] |
| Mineral Crystallinity Index 1 and 2 | CI1<br>CI2 | $I_{1030}/I_{1020}$<br>$I_{1060}/I_{1075}$ | max 1150-1250,<br>min 850-950 | 1175 - 900<br>1060 - 900 | [21]<br>Lebon et al., 2011) |
| **Organic diagenesis** |  |  |  |  |  |
| Amide I to Matrix (collagen content) | $A_{\text{ml}}/\text{P}$ | $I_{1660}/I_{\text{maxPO}_4}$ | max 1700-1800,<br>min 1300-1400 / | 1800 - 1300 /<br>1200 - 900 | [40] |
|  |  |  | max 1150-1250,<br>min 850-950 |  |  |
| Amide II to Matrix | AmlI/P | $I_{1550}/I_{\max \text{PO}_4}$ | max 1700-1800,<br>min 1300-1400 /<br>max 1150-1250,<br>min 850-950 | 1800 – 1300 /<br>1200 - 900 | [15] |
| Collagen denaturation ratio | Aml/AmlI | $I_{1660}/I_{1550}$ | max 1700-1800,<br>min 1300-1400 / | 1800 - 1300 | [15] |
| Collagen cross links in collagen ratio | Collagen Cross Link Integrity | $I_{1690}/I_{1660}$ | max 1700-1800,<br>min 1300-1400 / | 1720 - 1590 | [21] |
| Random coils to α-helix ratio | Random Coils ($\alpha$) | $I_{1645}/I_{1660}$ | max 1700-1800,<br>min 1300-1400 / | 1720 - 1590 | [21] |

Mineral composition alterations were evaluated using the carbonate-to-phosphate ratio (C/P; ν_₃_CO_₃_/ν_₃_PO_₄_) and hydrogen phosphate-to-phosphate ratio (HPO_₄_/P). Two crystallinity indices (CI1 and CI2) derived from the ν_₃_PO_₄_ region were measured to understand changes in the crystalline atomic order [21, 40]. Analysis of *v*_4_ phosphate region was not possible because it fell outside the measured spectral range.

Organic alterations were assessed using the amide I-to-phosphate (AmI/P) and amide II-to-phosphate (AmII/P) ratios as indicators of collagen-associated amide spectral signatures to assess degradation of collagen [15, 40]. Additional indices describing collagen-associated secondary structural features were calculated, specifically: the amide I-to-amide II ratio (AmI/AmII), collagen integrity ratio (1690/1660 cm^-1^), and random coil-to-α-helix ratio (1645/1660 cm^-1^) (Table 2) [15, 19, 21, 48].

Twenty-three hyperspectral images obtained by r-FTIRM were processed in Python (v 3.9.25) using Specio (v 0.1.0) [68]. The pre-processing python script can be found at Zenodo [69]. Baselines were defined as a linear tangent between bounding peak minima and maxima (Table 2) [15]. Ratio heat maps were generated for each diagenetic index (Table 2), and spectra-graphs were interpolated by linear curve fitting between neighbouring measurement points. Pixels contaminated by epoxy or biodegredation were excluded using a phosphate peak absorbance threshold (maxPO_4_, 1250-850 cm^-1^) of less than 0.3. Pixels with negative absorbance were further removed as outliers.

To quantify regional diagenetic differences within samples and between depositional groups, averages were calculated from pixels located 600-700µm within the sub-endosteal, mid-cortical, and sub-periosteal rows per ratio image (Figure 3). Row averaged values from each ratio image were also subject to a linear regression to assess trends between spectral indices and cortical position between the periosteal to endosteal bone limits; testing possible inward water diffusion pathways from bone surfaces. Statistical significance was evaluated at *p*<0.05, 0.01, and 0.001.

Synchrotron-FTIRM datasets were pre-processed in OPUS 8, using atmospheric compensation to remove the interference of atmospheric water vapour and carbon dioxide from the acquired spectra, followed by baseline correction. Wavenumber absorbance values (Table 2) were extracted from the entire mapped region (ROI; 625 spectra), the interstitial bone area (In.Ar), and the secondary osteon bone area (On.Ar) (Figure 3). The number of spectra within each tissue type are sample dependant (Supplementary 2). Spectra quality was assessed based on various peak absorbance thresholds to identify and remove empty pores, potential artefacts, contamination and taphonomy (Supplementary 2). Sample mean values for each diagenesis measure were calculated for bone type group (ROI, In.Ar, On.Ar) and outliers removed based on *z*-scores for each burial condition.

#### 2.4.4 Statistical analyses

Data are first summarised descriptively and visualised using R (version 3.6.0+) and RStudio (2026.05.0+218) using ggplot2 package [70] to produce box plots, kernel density violin plots, and principal component analysis plots. Violin plots are restricted to true values due to the small sample sizes. Non-parametric statistical tests were applied in SPSS (version 29) to determine if distribution and medians of the bone type groups differ between the three depositional environment conditions, and if there is a difference between the interstitial bone and secondary osteon bone areas for each burial context. Distributions were compared using Kruskal-Wallis (more than two groups compared) or Mann-Whitney U tests (for two-sample comparisons), and medians compared using Independent-Samples Median Test.

Principal component analyses (PCA) were conducted to identify if diagenesis measures significantly statistically varied across depositional environments, or bone tissue types. Principal component values were further tested using non-parametric distribution and median tests to identify significant data differences. Quality of models were assessed based on the Kaiser-Meyer-Olkin (KMO) test for suitability (>0.5), Bartlett’s test of sphericity (<0.05), Eigenvalues greater than one, and cumulative variance (>70%).

## 3 Results

Bone histology was well preserved across all specimens analysed. Polarised light microscopic images are available in Supplementary 2.

Organic and mineral diagenetic indices were quantified for the sub-periosteal, mid-cortical, and sub-endosteal bone regions, and averages produced for the different depositional environment conditions (Supplementary 3). Cleaned, r-FTIRM chemical heat maps for each specimen and diagenetic index are available from Zenodo [69] Group averages and descriptive statistics for the s-FTIRM micron scale ROIs, In.Ar, and On.Ar of the Haversian systems are further provided in Supplementary 3.

Of the 42 secondary osteons measured from 21 specimens analysed using s-FTIRM, three were excluded because of imaging artefacts or incomplete crystal-sample contact during ATR mapping, leaving 39 secondary osteons for analysis (Supplementary 1). Thus, fossil samples 12.0.1_7 and 32_11 were excluded from the analysis. After spectral quality screening, 19,886 spectra (81.4% of the acquired dataset) were retained for analysis, including 8,493 spectra from On.Ar and 8,250 from In.Ar (Supplementary 1).

### 3.1 Diagenesis markers compared between historic wet and dry bone assemblages

#### 3.1.1 Histotaphonomy of historic samples: birefringence

Birefringence was apparent in bone areas unaffected by biodegradation in both wet and dry depositional environments [65] (Figure 5; Supplementary 2). Most wet samples showed birefringence comparable to fresh bone (BI(1) = 50.0%%; BI(0.5) = 37.5%; BI(0) = 12.5%), whereas dry samples presented with a mix of birefringence equivalent to reduced (BI(0.5) = 50%) and fresh bone (BI(1) = 50%) levels. Although reduced birefringence was locally heterogeneous, this variation was not reflected in the overall BI. Bones showed absence of birefringence in regions with diagenetic bone staining resulting from the local burial conditions (Figure 5:C1-C2), or areas with microbial attack (Figure 5: B1-B2). Stained bone regions maintained bone microstructural integrity whereas bacterial attack removed bone mineral and collagen through enzyme hydrolysis [12].

**Figure 5.**
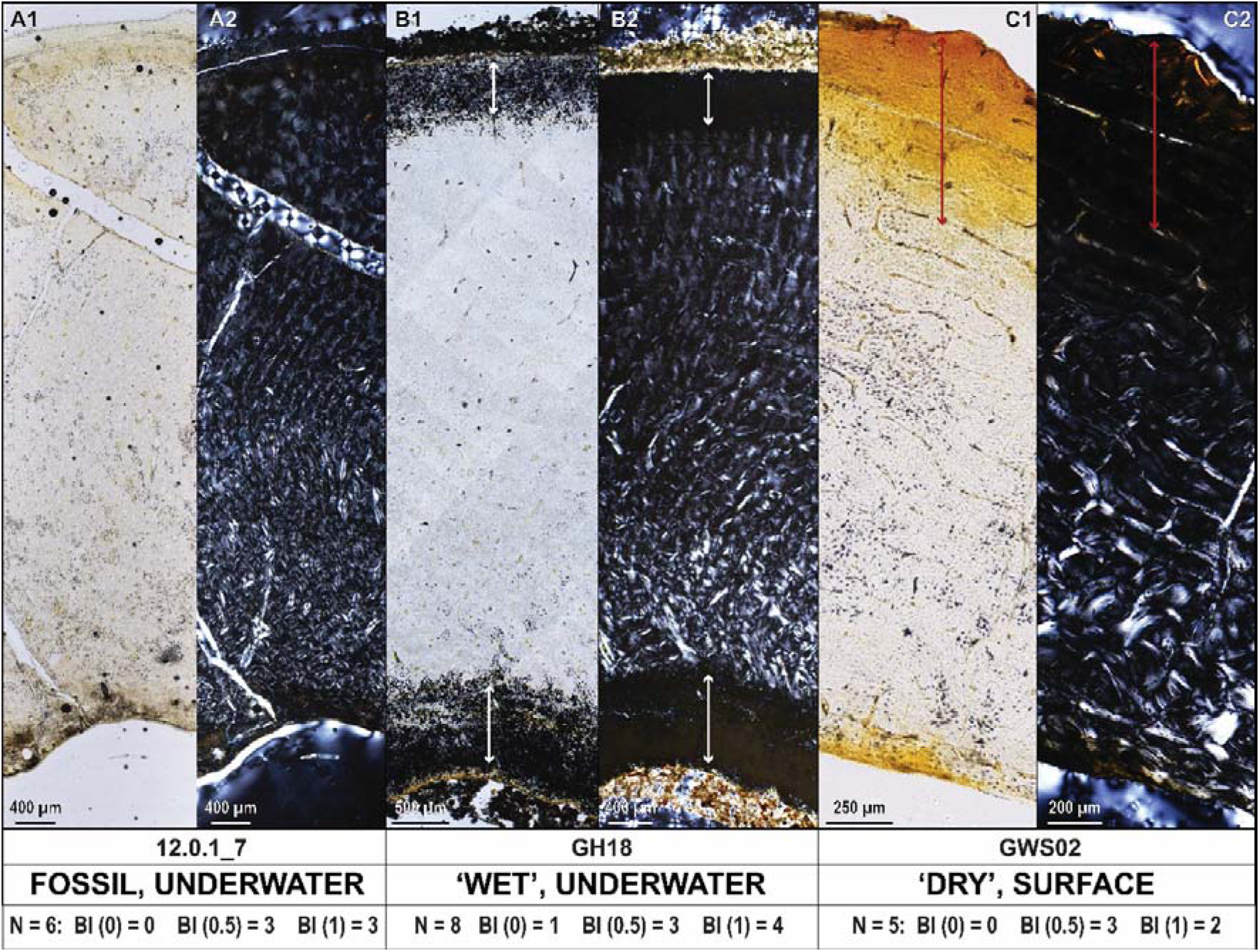
Bone preservation and birefringence under transmitted (1) and polarised (2) light from fossil (A), historic wet (B), and historic dry (C) conditions. Number of specimens analysed and birefringence index (BI) data are presented for each sub-assemblage. White arrows indicate regions of bioerosion, and red arrows indicate stained bone areas.

#### 3.1.2 FTIRM analysis

Dry and wet historical specimens showed different diagenetic patterns depending on the bone region analysed (Figure 6, Figure 7, Table 3). No significant differences in mineral or organic composition were detected in the mid-cortical region, although this region exhibited the greatest variability in collagen-associated spectral indices.

**Figure 6.**
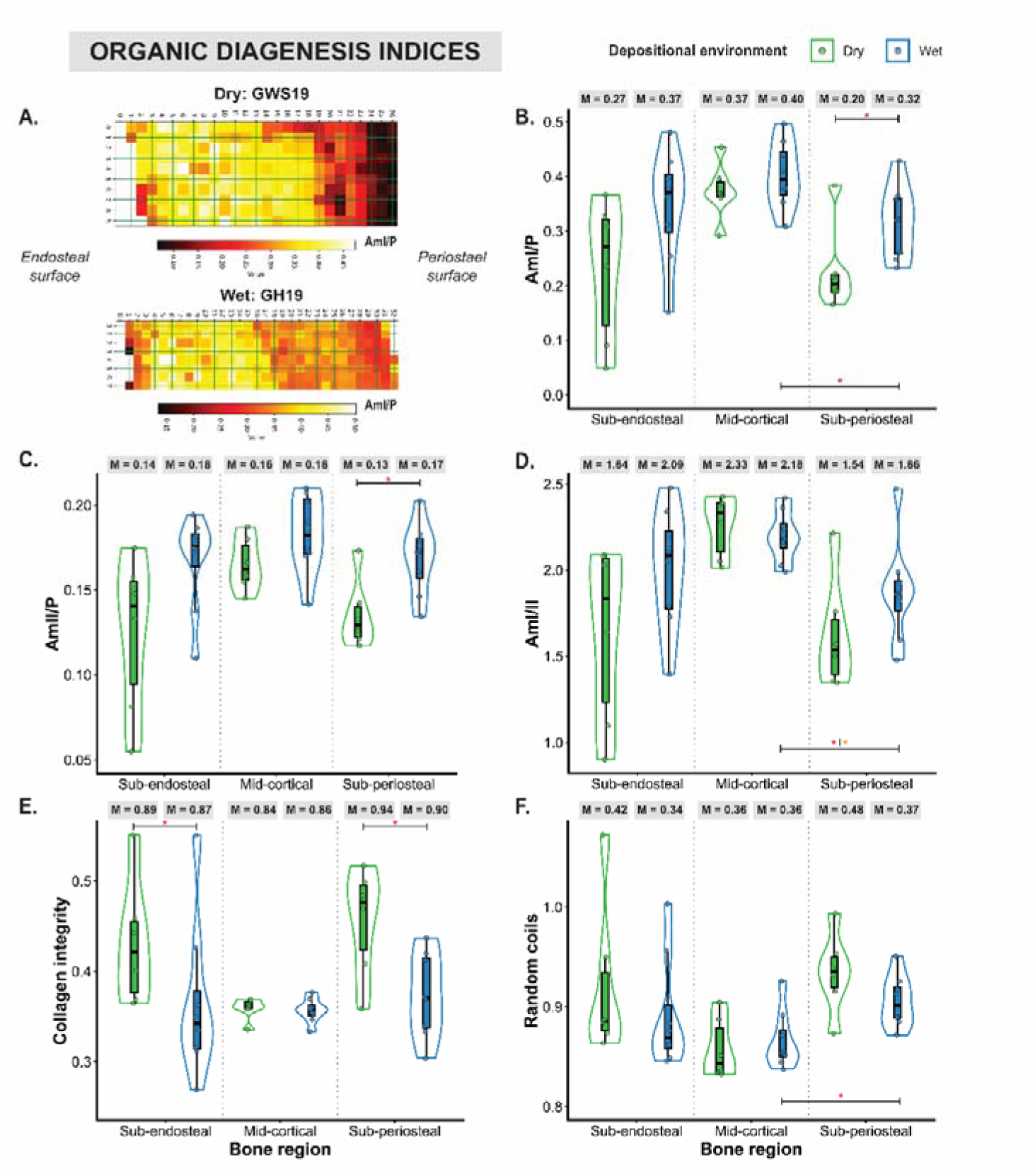
Organic diagenesis measured using reflectance Fourier-transform infrared microspectroscopy across bone regions and environments of historical samples: a.) AmI/P chemical mapping examples from each depositional environment, b.) amide I to phosphate (Am1/P), c.) amide II to phosphate (AmII/P), d.) amide I to amide II (AmI/II), e.) random coils to alpha helix, and f.) collagen cross links as collagen integrity. Median (M) values presented at the top of each plot. Statistically significant differences (CI 95%) are represented by red star for distributions, and orange for median comparisons. Only regional differences for wet environments were tested.

**Figure 7.**
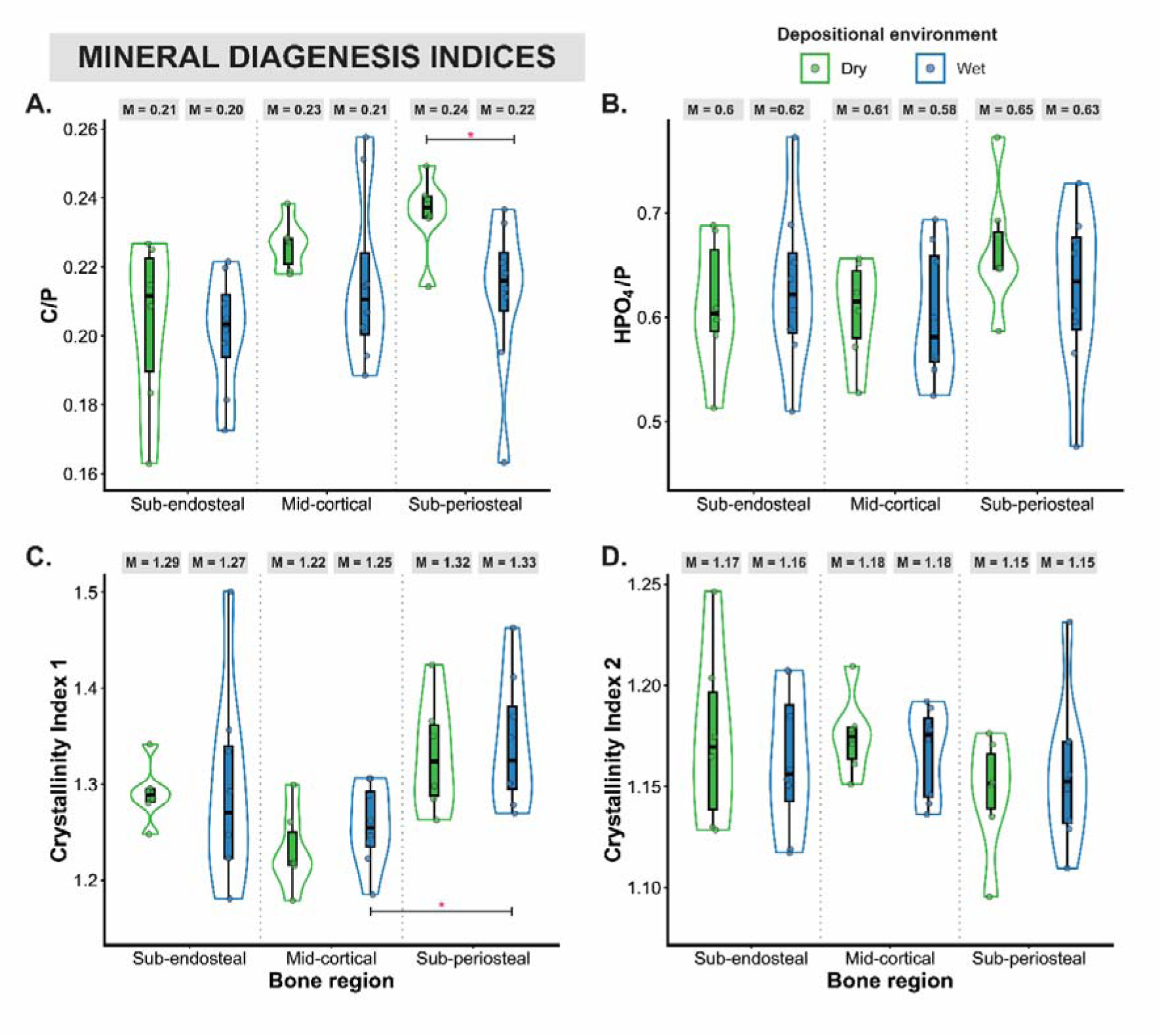
Mineral diagenesis measured using reflectance Fourier-transform infrared microscopy across bone regions, and wet and dry environments: a.) carbonate to phosphate (C/P), b.) hydrogen phosphate to phosphate (HPO_4_/P), c.) Crystallinity Index 1, and d.) Crystallinity Index 2. Median (M) values presented at the top of each plot. Statistically significant differences (CI 95%) are represented by red star for distribution comparisons. Only regional differences for wet environments were tested.

**Figure 8.**
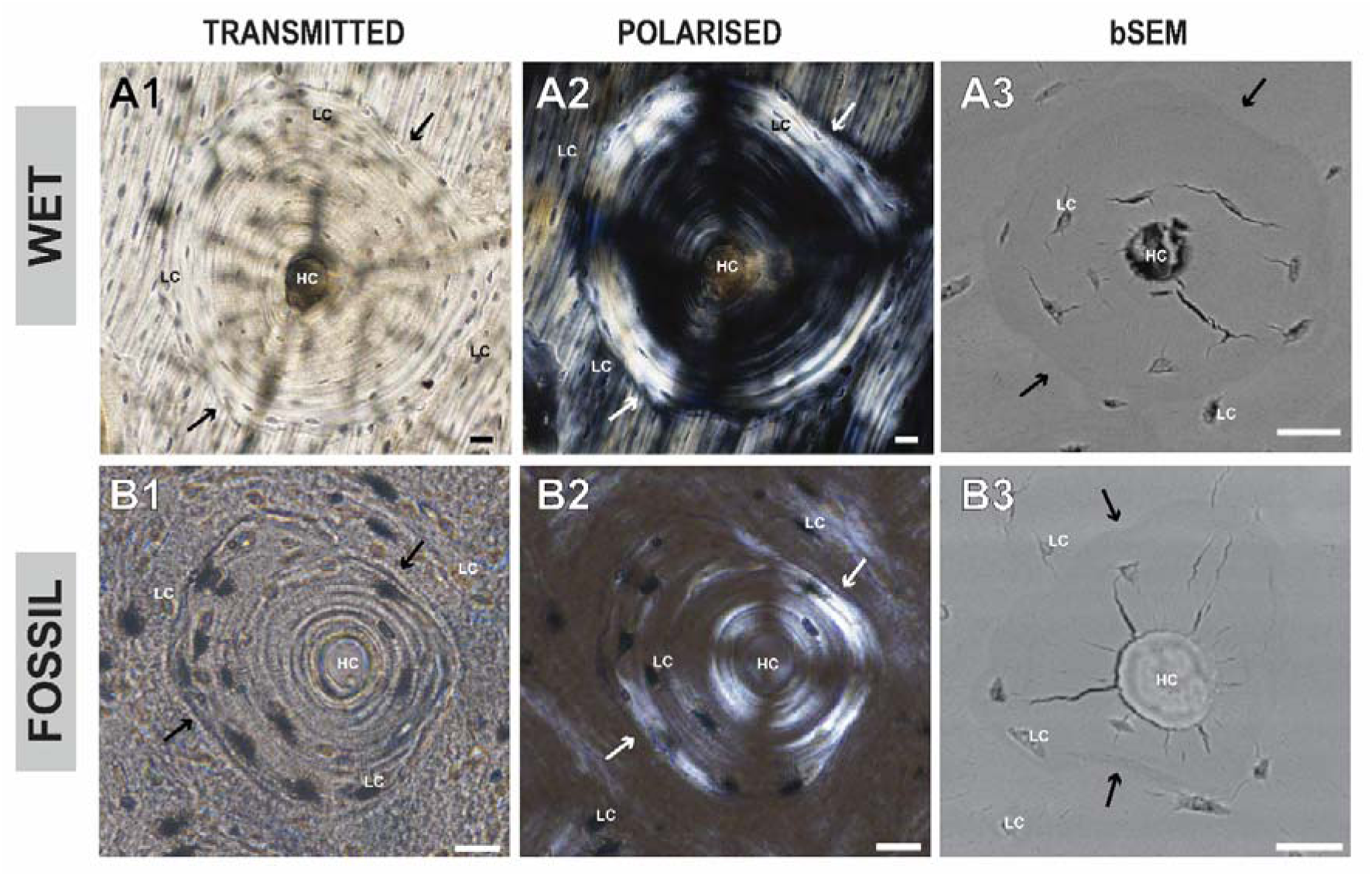
Example of secondary osteons with intact cement lines in wet (A1-A3) and fossil (B1-3) specimens under transmitted (A1,B1) and polarised (A2,B2) optical light microscopy, and backscatter scanning electron microscopy (bSEM; A3,B3). Cement lines are indicated by black and white arrows alongside Haversian canals (HC) and osteocyte lacunae (Lc). Scale bars measure 20 microns.

**Table 3.** Descriptive statistics of region of interest (ROI) data across wet, dry and fossil bone assemblages. Collagen quality indicators for the fossil condition are reported, but due to severe degradation are not considered a reliable indicator of collagen preservation.

|  | Wet |  |  |  |  | Dry |  |  |  |  | Fossil |  |  |  |  |
| --- | --- | --- | --- | --- | --- | --- | --- | --- | --- | --- | --- | --- | --- | --- | --- |
| Index | N | x̄ | s | σ | Range | N | x̄ | s | σ | Range | N | x̄ | s | σ | Range |
| C/P | 14 | 0.25 | 0.25 | 0.03 | 0.18 -0.31 | 11 | 0.26 | 0.27 | 0.03 | 0.22-0.30 | 14 | 0.22 | 0.22 | 0.03 | 0.18-0.29 |
| HPO<sub>4</sub>/P | 14 | 0.23 | 0.23 | 0.03 | 0.17-0.27 | 11 | 0.23 | 0.23 | 0.04 | 0.14-0.29 | 14 | 0.18 | 0.18 | 0.04 | 0.07-0.21 |
| CI1 | 14 | 0.95 | 0.95 | 0.02 | 0.91-1.00 | 11 | 0.94 | 0.94 | 0.02 | 0.89-0.97 | 14 | 0.99 | 0.99 | 0.02 | 0.95-1.01 |
| CI2 | 14 | 1.38 | 1.37 | 0.06 | 1.31-1.53 | 11 | 1.37 | 1.35 | 0.07 | 1.31-1.56 | 14 | 1.55 | 1.51 | 0.15 | 1.45-1.97 |
| Aml/P | 14 | 0.29 | 0.29 | 0.06 | 0.17-0.41 | 11 | 0.28 | 0.28 | 0.05 | 0.21-0.36 | 14 | 0.03 | 0.02 | 0.01 | 0.02-0.06 |
| AmlI/P | 14 | 0.23 | 0.24 | 0.04 | 0.14-0.31 | 11 | 0.23 | 0.22 | 0.03 | 0.19-0.28 | 14 | 0.06 | 0.05 | 0.01 | 0.05-0.09 |
| Aml/ AmlI | 14 | 1.43 | 1.42 | 0.12 | 1.29-1.71 | 11 | 1.48 | 1.48 | 0.05 | 1.14-1.59 | 14 | 0.69 | 0.70 | 0.27 | 0.32-1.24 |
| Collagen Integrity | 14 | 0.28 | 0.29 | 0.04 | 0.18-0.33 | 11 | 0.28 | 0.28 | 0.03 | 0.21-0.31 | 13 | 0.37 | 0.20 | 0.46 | 0.13 – 1.80 |
| Random Coils (α) | 14 | 0.95 | 0.95 | 0.03 | 0.90-1.03 | 11 | 0.95 | 0.95 | 0.03 | 0.90-1.01 | 14 | 4.04 | 3.32 | 2.75 | 0.95-8.25 |

Organic diagenetic patterns differed between wet and dry depositional environments, with the largest differences observed in the sub-periosteal cortex (Figure 6). The AmI/P (wet x□ = 0.32, dry x□ = 0.20, *p* = 0.03) and AmII/P (wet x□ = 0.17, dry x□ = 0.13, *p* = 0.01) ratios, collected by r-FTIRM analysis, were significantly higher in the sub-periosteal region of wet specimens, indicating greater preservation of collagen-associated amide spectral signatures. No differences in AmI/P or AmII/P were observed within the sub-endosteal region, consistent with the micron-scale ROI analysis, and thus there is a similarity across analytical scales (Table 10 in Supplementary 3). The collagen integrity ratio (1690/1660 cm_⁻_¹) was also greater in dry conditions in both the sub-periosteal (wet x□ = 0.37, dry x□ = 0.47, *p* = 0.04) and sub-endosteal (wet x□ = 0.34, dry x□ = 0.42, *p* = 0.04) regions. These differences were not observed in the ROI data, located in the sub-endosteal cortex, that showed consistently similar values. Among the remaining collagen-associated indices, only the AmI/AmII ratio differed at the micron scale, with higher values observed in dry specimens compared to the wet counterparts (*p*=0.047) (Table 3).

Mineral-related spectral indices were largely consistent between wet and dry conditions for each bone region (Figure 7), and at the micron-scale analysis of the Haversian system ROI (Table 10 in Supplementary 3). Only the C/P proportion distributions were significantly lower in the sub-periosteal region of wet the assemblage (Figure 7; *p* = 0.02). Although differences in the C/P values were not observed in the sub-endosteal region, micron-scale ROI analysis showed that the median proportion of carbonate is lower in the wet assemblage than the dry (Table 3, *p*=0.047). Levels of hydrogen phosphate and crystallinity indices were consistent between depositional conditions across all bone regions and the ROI.

### 3.2 Diagenesis markers examined within historic wet specimens

#### 3.2.1 Histotaphonomy of historic samples: microcracking

Under bSEM, and polarised and transmitted light optical microscopy of secondary osteons, no radial microfractures across cement lines were identified (Figure 6). However, other types of secondary osteon fracturing were identified across some samples under bSEM, including radial microfractures extending from Haversian canals, and shrinkage and separation of the secondary osteon at the cement line from the surrounding matrix. Whilst cracking was observed across secondary osteon cement lines, fractures either originated from the Haversian canal or were part of a broader generalised cracking pattern across the bone matrix. Microfractures were not identified under transmitted light microscopy and are thus likely an artefact of the pressurised SEM chamber.

#### 3.2.1 FTIRM analysis

There was limited variation in diagenesis markers across bone cortices of the wet samples, with statistically significant differences found only between the sub-periosteal and mid-cortical regions (Figure 6–Figure 7, Table 7 in Supplementary 3). Organic indices showed the greatest regional variation. The sub-periosteal cortex exhibited lower AmI/P values (*p* = 0.03), resulting in a corresponding increase in the AmI/AmII ratio (*p* = 0.01).Whilst the subperiosteal region also showed greater degradation in random coils-to-alpha-helix structure (*p*=0.03), the collagen integrity ratio (collagen cross-linking) remained unchanged. Among mineral indices, only CI1 differed significantly (*p* = 0.02), indicating greater crystalline order within the sub-periosteal cortex. No regional difference was observed for CI2.

In the wet bones, statistically significant regressions were observed between the periosteal and endosteal bone surfaces for 55.6% of the samples and diagenesis indices (Table 4). Although this may reflect a linear water diffusion pathway and associated diagenesis pathway from the bone surface into the cortex, of the statistically significant models, only 15% had correlation coefficients greater than 0.5, and *r* values were not consistently positive or negative across specific indices and burial conditions (Supplementary Table 1).

**Table 4.**
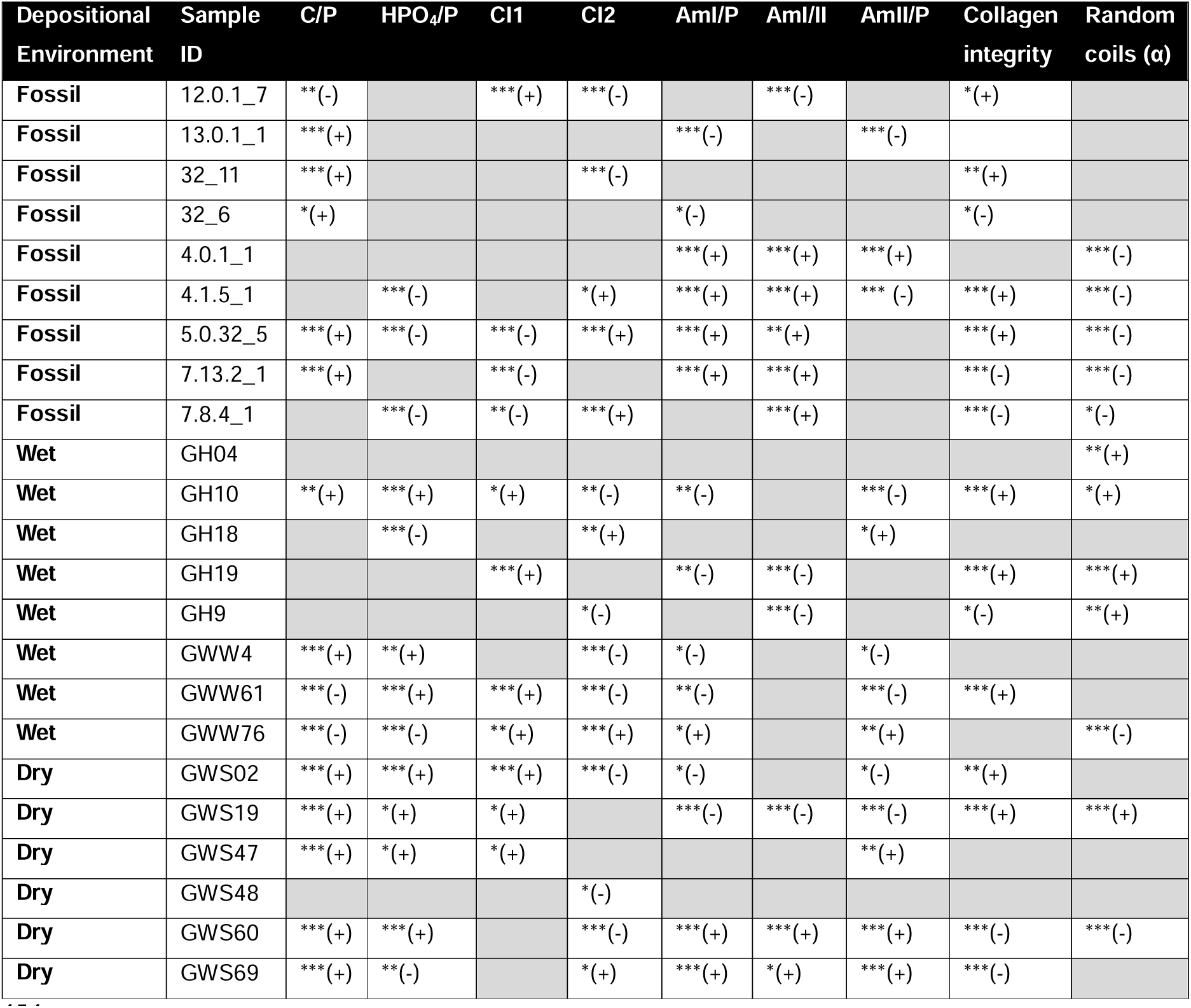
Pearson’s linear correlation analysis of diagenesis across bone cortex from periosteal to endosteal surfaces. Statistical significance indicated by asterisk to thresholds of 0.05 (*), 0.01 (**) and 0.001 (***). Directionality of significance indicated as either a negative (-) or positive (+) correlation.

Interstitial and secondary osteon bone were compared in wet condition specimens (Figure 9, Table 11 in Supplementary 3). Distributions of all mineral diagenetic indices were significantly different between interstitial and secondary osteon bone, (C/P *p*=0.04; HPO_4_ *p*=0.01; CI2 *p*=0.02), with additional median difference in CI1 (*p*=<0.01). Carbonate and hydrogen phosphate proportions were greater in secondary osteon bone (Figure 9c-d), but crystallinity indices varied (Figure 9e-f). No organic indices exhibited significant differences across bone tissue bounding the secondary osteon cement wall (Supplementary 3).

**Figure 9.**
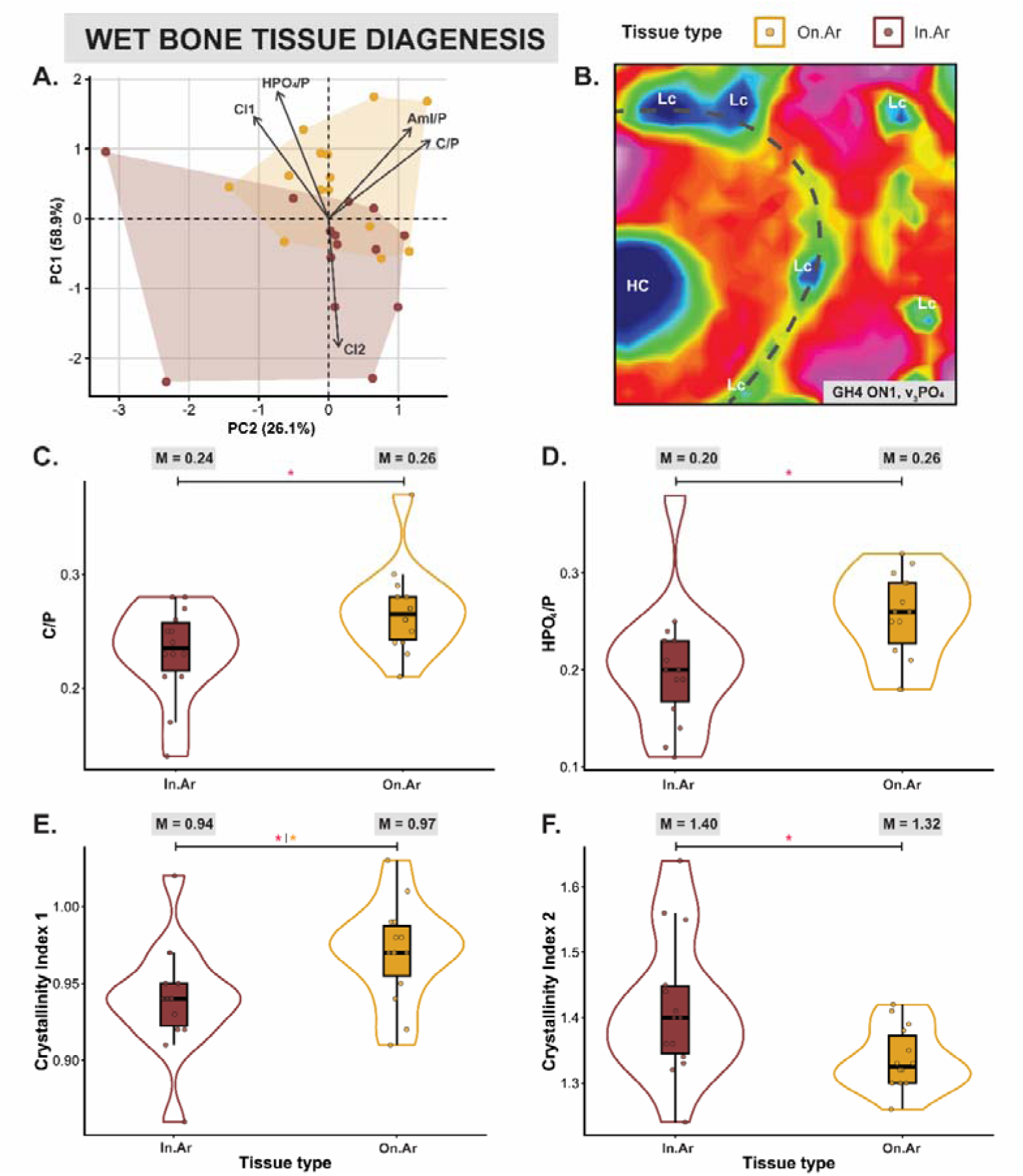
Synchrotron Fourier-transform infrared microspectroscopy (s-FTIRM) analysis of interstitial (In.Ar, n=14) and secondary osteon (On.Ar, n=14) bone tissues from the wet assemblage. (A) principal component analysis (PCA) based on the proportion of amide I to phosphate (AmI/P) and all other mineral diagenetic indices, component loadings represented by grey arrows; (B) example of secondary osteon (ID: GH64, Osteon 1) s-FTIRM chemical heat map for v_3_ PO_4_ curve area with grey dashed cement line, haversian canal (HC) and osteocyte lacunae (Lc) indicated; (C—F) bar charts and violin plots comparing In.Ar and On.Ar bone for carbonate to phosphate (C; C/P), hydrogen phosphate to phosphate (D; HPO4/P), crystallinity index 1 (E; CI1), and crystallinity index 2 (F; CI2). Median values are presented for each group, and statistical significance identified by either a red (distribution comparison) or orange (median comparison) star.

Principal component analysis data further supported the separation between interstitial and secondary osteon bone data in wet depositional conditions. Significant differences were observed for combined mineral and organic variables (PC1: *p*=0.02; PC2: *p*=0.03), mineral and AmI/P (PC1: *p*=0.004) and mineral only variables (PC1: *p*=0.005). The most robust PCA model included the mineral indices together with AmI/P content (Figure 9a; Table 12 in Supplementary 3) meeting the predefined suitability criteria (KMO = 0.543, sphericity = <0.01, cumulative variance = 85.02). In this model, On.Ar data are more tightly clustered compared to In.Ar data that are influenced by CI2 levels.

### 3.3 Diagenesis markers in fossil specimens

#### 3.3.1 Depositional environment

Bone structures examined using histological methods were well preserved across all fossil specimens (N=9), with only one specimen (ID 32_11) presenting with greater porosity throughout the mid-cortical region. Birefringence was similar to dry bone specimens, with mixed scores varying between fresh and reduced bone (BI(0.5) = 50%; BI(1) = 50%). As observed across dry and wet specimens, birefringence was heterogenous across the bone cortex. Birefringence was the strongest in the sub-endosteal region associated with a dense pocket secondary osteons (Figure 5).

Spectral data displayed strong carbonate and phosphate peaks, and a considerable loss of amide I and amide II in fossil samples compared to historical samples (Figure 10; Supplementary 1). One sample observed in the r-FTIRM analysis showed well preserved AmI/P and AmII/P, with average values greater than 0.1 (ID 12.0.1_7; Table 1 in Supplementary 3). This sample was excluded from s-FTIRM analysis through the data quality screening step and thus comparisons between scales are not attainable. Due to heavy amide I and II degradation, meaningful assessments of collagen content and quality between fossil and historic bones and between different fossil bone regions and tissues were not possible.

**Figure 10.**
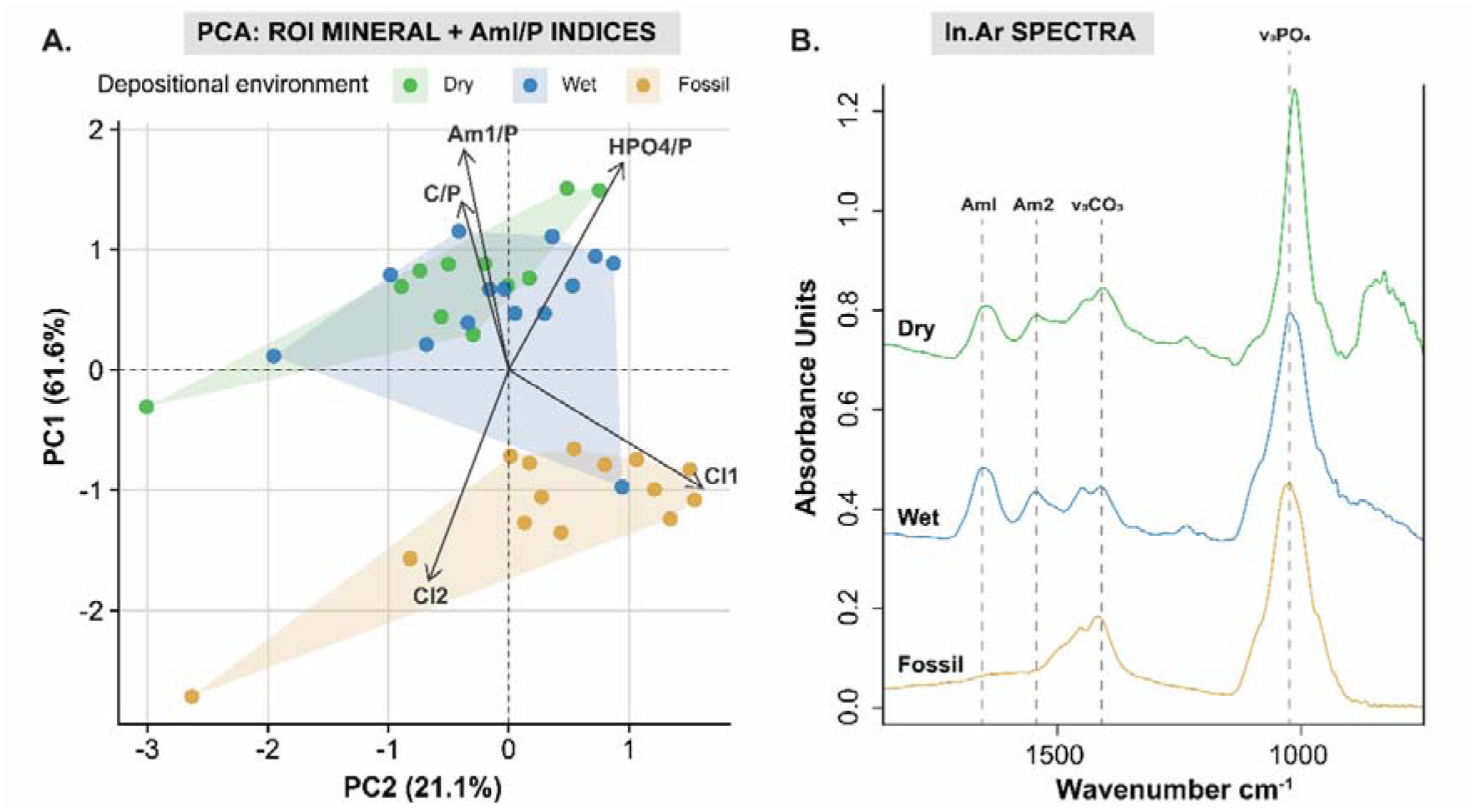
a. Principal component analysis (PCA) scree plot of region of interest data (ROI) for each depositional environment, based on mineral indices and the proportion of amide I to phosphate (AmI/P). Component loadings are represented by grey arrows. b. discrete spectra between ∼1800-750cm-1 extracted from interstitial area (In.Ar) bone tissue for dry (green; ID: GWS48 On2), wet (blue; ID: GWW4 On1) and fossil (yellow; ID: 32_6 On3). Amide I (AmI) amide II (AmII) carbonate (v3CO3) and phosphte (v3PO4) peaks are identified by dotted lines across each spectra. Note: spectra are shifted vertically and do not all start at 0 absorbance units.

Principal component analysis results of ROI data across the mineral and organic indices consistently isolated the fossil context from wet and dry conditions, driven by differences between C/P, AmI/P and HPO_4_/P values with crystallinity indicators in PC1 (Figure 10). Separation of conditions was best presented by the modelling across all mineral indices and only the Am/P organic diagenesis index (Figure 10; KMO = 0.597, sphericity = <0.001, cumulative variance = 84.25%, two eigenvalues >1). Statistical comparisons of the principal components between depositional environments separate the fossil and dry assemblage across both PC1 (*p*=<0.00) and PC2 (*p*=0.03), whereas only PC1 was significantly different between the wet and fossil assemblages (*p*=<0.00) (Table 13 in Supplementary 3).

Individual comparisons of mineral content diagenetic indices show considerable differences between the fossil and the dry and wet bone samples; however, the differences are more pronounced in comparisons between the fossil and dry group (Table 5). Dry and fossil bones were significantly different in all measures in the sub-periosteal and mid-cortical bone regions, and ROIs (Table 5). However, the sub-endosteal bone region showed the greatest similarities in mineral diagenesis between depositional environments (Table 5). Whilst median CI1 values were significantly elevated in fossil (x □=0.1.69) compared to dry (x□ =0.1.29) and wet (x□ =1.27) specimens, there were no differences in C/P or HPO_4_/P in any depositional environment comparison across the sub-endosteal bone region (Table 5, Supplementary 3).

**Table 5.** Bone regional differences in mineral diagenesis comparing the wet (n=8) and dry (n=6) specimens with the fossils (n=9). Bold values with an asterisk are significant at a 95% confidence interval.

|  |  | Sub-endosteal |  | Mid-cortical |  | Sub-periosteal |  | ROI |  |
| --- | --- | --- | --- | --- | --- | --- | --- | --- | --- |
| Diagenesis Index |  | Test statistic | p-value | Test statistic | p-value | Test statistic | p-value | Test statistic | p-value |
| **Wet vs. Fossil** |  |  |  |  |  |  |  |  |  |
| C/P | Distribution | 53.000 | 0.114 | 63.000 | **0.008\*** | 46.000 | 0.370 | 154.000 | **.009\*** |
|  | Median | 1.446 | 0.347 | 4.735 | 0.057 | 0.052 | 1.000 | 5.143 | .057 |
| HPO<sub>4</sub>/P | Distribution | 53.000 | 0.114 | 60.000 | **0.021\*** | 62.000 | **0.011\*** | 173.000 | **.000\*** |
|  | Median | 1.446 | 0.347 | 4.735 | 0.057 | 4.735 | 0.57 | 9.143a | **.007\*** |
| CI1 | Distribution | 6.000 | **0.002\*** | 7.000 | **0.004\*** | 4.000 | **0.001\*** | 24.000 | **.000\*** |
|  | Median | 7.244 | **0.015\*** | 13.432 | **0.000\*** | 7.244 | **0.015\*** | 14.286 | **.000\*** |
| CI2 | Distribution | 9.000 | **0.008\*** | 10.000 | **0.011\*** | 5.000 | **0.002\*** | 13.000 | **.000\*** |
|  | Median | 2.951 | 0.153 | 7.244 | **0.015\*** | 7.244 | **0.015\*** | 14.286 | **.000\*** |
| **Dry vs. Fossil** |  |  |  |  |  |  |  |  |  |
| C/P | Distribution | 15.000 | 0.181 | 0.000 | **0.000\*** | 7.000 | **0.018\*** | 23.000 | **.002\*** |
|  | Median | 1.607 | 0.315 | 11.429 | **0.001\*** | 5.402 | **0.041\*** | 4.812 | **.047\*** |
| HPO<sub>4</sub>/P | Distribution | 16.000 | 0.224 | 9.000 | **0.036\*** | 1.000 | **0.001\*** | 12.000 | **.000\*** |
|  | Median | 1.607 | 0.315 | 5.402 | **0.041\*** | 11.429 | **0.001\*** | 14.490 | **.000\*** |
| CI1 | Distribution | 48.000 | **0.012\*** | 49.000 | **0.008\*** | 53.000 | **0.001\*** | 149.000 | **.000\*** |
|  | Median | 8.750 | **0.007\*** | 8.750 | **0.007\*** | 8.750 | **0.007\*** | 18.132 | **.000\*** |
| CI2 | Distribution | 43.000 | 0.066 | 47.000 | **0.018\*** | 53.000 | **0.001\*** | 142.000 | **.000\*** |
|  | Median | 0.714 | 0.608 | 8.750 | **0.007\*** | 8.750 | **0.007\*** | 1.464 | **.000\*** |

Carbonate content showed the greatest similarities between the fossil and historic wet assemblages in the sub-periosteal and sub-endosteal bone (Table 5). Wet and fossil bones contained similar proportions of carbonate to phosphate in the sub-periosteal bone region (distribution *p*=0.37; median *p*=1.00), both exhibiting significantly lower levels compared to the dry bone assemblage (wet *vs* dry: distribution *p*=0.02; dry v fossil: distribution *p*=0.02; median *p*=0.04; Supplementary 3). Here, median C/P proportions are greatest in the dry assemblage (x□ =0.24), followed by wet (x□=0.22) and then fossil bones (x□=0.21) (Supplementary 3). Median ROI was further statistically similar between wet and fossil depositional environments, however distributions were consistent (Table 5).

Biomineral alterations across the fossil samples show significantly greater crystallinity for all bone regions than wet and dry conditions (Table 5). However, crystallinity indicator 1 and CI2 were not consistent, with statistically significant similarities in CI2 but differences in CI1 between all environments in the sub-endosteal bone region (Table 5).

#### 3.3.2 Bone tissue specific diagenesis

In the fossil samples, r-FTIRM digenesis markers were consistent across all bone regions (Table 8 in Supplementary 3). Differences in wet bone tissue degradation across the cortex were observed in the organic fraction, which cannot be tested in the fossil samples due to the loss of amide I and II. The only distribution to vary significantly is the proportion of hydrogen phosphate to phosphate, with greater values in the sub-endosteal region compared to the mid-cortex (*p*=0.05). This difference in HPO_4_/P was not observed in either wet or dry assemblages.

Diagenetic trends across the bone cortex were not consistent across the fossil samples (Table 4). Regression analysis was only statistically significant in 62.5% of cases, with 17.8% of these associated with correlation coefficients above 0.5 (Supplementary Table 1). Similarly to wet conditions, *r* values were not consistently positive or negative and thus do not exhibit a pattern that reflect a slow diffusion of water from the exterior bone surface into the cortex.

No structural or biomolecular differences were identified when comparing bone tissue types in the fossil samples. No micro-fracturing was identified across secondary osteon cement lines (Figure 6), although cracking from the Haversian canal was identified as a product of the pressurised SEM chamber. Unlike the mineralogical differences observed between tissue types for the wet bone assemblage, no differences were observed between interstitial and secondary osteon bone in the fossil samples in the individual index value comparisons or PCA analysis (Table 14 in Supplementary 3).

## 4 Discussion

In our study, bone organic and mineral preservation varied intra-skeletally depending on whether the depositional environment was wet or dry, and whether the specimen was from a historic and geologic timeline. Our analyses showed that wet historic specimens presented greater levels of organic preservation compared to those from historic dry depositional environment, but preservation was heterogenous across the bone cortex. However, organic preservation was consistent between secondary osteonal and interstitial bone tissues, and no association with radial microfractures across the secondary osteon cement line was observed. Instead, proportions of carbonate and hydrogen phosphate were greater in the secondary osteon tissue. Amide in fossil specimens was completely degraded, however bone birefringence levels were consistent across the depositional environments. Fossil samples were most like historic bone from wet environments, with similar proportions of carbonate despite greater crystallinity in the fossil groups. Below we discuss these key findings.

### 4.1 Diagenetic trajectories in wet and dry cave environments

Historical bones from underwater deposits showed greater preservation of amide I (AmI/P) and II (AmII/P) compared to bones from the dry cave environment. However, this was only observed across the r-FTIRM analyses and not at the s-FTIRM micron scale. Collagen quality observed using both FTIRM analyses was also better preserved in wet cave landscapes compared to dry conditions which showed more collagen structural decay despite similar levels of organic content preservation.

Preservation of collagen and its structure in cave waters is atypical compared to data from other aquatic sites [46] and modelling of collagen decay in water [12, 26]. These studies led to the suggestion that aquatic environments increase collagen hydrolysis and gelatinisation, increasing degradation during bone diagenesis [12, 26]. This process is accelerated by higher temperatures and extreme pH levels [18], and increases with hydraulic flow and cyclical inundation that refreshes water chemistry and modifies the aquatic environment [36]. In a separate experimental study of alligator bones, all samples submerged in synthetic wetland solutions varying from pH 7 to 11 showed collagen loss after 24 hours [25]. Although a modern bone benchmark was not analysed in this study, our data nevertheless indicate that collagen is well preserved in cave waters. The underwater caves investigated here are characterised by constant temperatures (∼16°C), neutral pH (7.4), and extremely low to no flow rates, possibly aiding the preservation of collagen.

Bones from wet environments presented with lower carbonate to phosphate ratios (C/P) in the outer cortex compared to dry environments. These differences are, however, likely a result of elevated C/P levels in dry cave conditions that also show a significant linear decline in carbonate from the exterior to interior bone surface, which was not observed in wet bone specimens. Research on archaeological human femoral bone shafts from Egypt [39] showed preferential mineral degradation of exterior cortical bone surfaces compared to inner cortices and trabecular bone, which was not observed here. Ionic uptake from the surrounding soils may instead artificially increase or preserve carbonate content in the outer bone regions of dry bone samples, but not wet bones that are in a constant aquatic environment.

Historic wet and dry bone assemblages present no differences in crystallinity measures in any bone region or tissue types. Bone dissolution is limited in neutral water [71, 72], however, bone from experimental studies simulating wetland conditions show recrystallisation and mineral uptake within one year of deposition under various pH levels [25]. Data shown here suggest that even if early recrystallisation is occurring, wet and dry assemblages from caves follow similar diagenetic patterns at the decadal scale (1841 – 1969).

### 4.2 Diagenesis across bone tissue types in underwater caves

Sub-periosteal bone regions in wet samples showed more collagen degradation and recrystallisation than the mid-cortical or sub-endosteal bone regions. Since degradation of collagen exposes the mineral crystalline lattice to modification, our results follow diagenetic patterns observed in other archaeological assemblages [15, 21]. Targeted modification of the exterior bone region further follows research that show the outer cortex is more impacted than inner cortical and trabecular structures due to the direct interactions between bone surfaces and local depositional environments [39, 73, 74]. In aquatic environments, modification of bone organics and mineral may reflect the heterogenous diffusion of water across bone cortices [35]. However, with similarities between the mid-cortical and sub-endosteal bone regions, greater mineral and organic diagenesis in the outer region may be induced by changing water chemistry in the local environment [36].

Microfractures were not identified across secondary osteon cement lines across either wet or dry historical samples or fossil specimens in our study. Such microfractures have been used in palaeontological and archaeological research as an indicator of underwater collagen gelatinisation patterns due to burial conditions [35, 43]. Whilst our FTIRM data show preservation of collagen and its structures in historical specimens, differential collagen decay is theoretically linked to early diagenesis and the rate of water diffusion across structures [42]. After decades of submersion, our wet specimens were highly saturated. Thus, actualistic and observational taphonomic data presented here, spanning ∼60,000 years, shows no relationship between aquatic cave submersion, collagen decay, and the production of radial microfractures. Instead, wet burial conditions show differences in mineral content across secondary osteon and interstitial bone tissues.

Interstitial and secondary osteon bone in wet conditions was best differentiated using PCA analysis using a model that incorporated mineral and AmI/P inputs. Heterogeneous diagenesis has been noted across micro-scale bone tissue analyses but has not been linked to specific tissue structures [21, 40]. Here we observed that secondary osteons presented greater carbonate and hydrogen phosphate values compared to interstitial bone. This may be associated with greater preservation [15, 23] or the deposition of secondary calcite that overlaps with the *v*_3_CO_3_ vibrational band [32.]. Crystallinity indicators in our data indicated differences between tissues, but CI1 was higher in secondary osteon bone and CI2 higher in interstitial bone. These measures were developed to mirror the infrared splitting factor (IRSF) in the 500 – 675cm^-1^ spectral range not measured here [21, 74, 75], but may not be a direct proxy for this traditional method. Nevertheless, diffusion of water through blood vessels may thus have some impact on bone diagenesis in the underwater cave environments examined here, however, differences are in the mineral content modifications, not collagen degradation.

Taken together, the histological and s-FTIRM results demonstrate that preservation of bone microarchitecture does not necessarily correspond to preservation of its original biochemical composition. Although birefringence and secondary osteon architecture remained largely intact across wet, dry and fossil specimens, s-FTIRM revealed substantial differences in collagen-associated amide spectral signatures and mineral composition. These findings suggest that histological integrity alone may overestimate molecular preservation in submerged cave assemblages and highlight the value of combining histological and spectroscopic approaches when reconstructing diagenetic histories.

### 4.3 Diagenesis of fossil specimens

Long term submergence represented by the palaeontological assemblage clearly results in the complete degradation of amide I and II, similar to other archaeological [32], and palaeontological cave specimens [76]. Bones may be buffered from organic degradation in these sites across decadal and centennial timescales; however, the fossils suggest amide structures are heavily impacted over tens of thousands of years of inundation. It is also possible that the remains experienced cyclical hydraulic events as changing glacial phases though the Late Pleistocene modified both sea and ground water levels [5, 77]. While historic specimens were collected from known wet or dry burial conditions, the original hydrological conditions of the palaeontological specimens are unknown, and changing depositional environments would likely increase collagen diagenesis [18, 36]. Organic loss may also be partially explained by the impact of museum curation over 30 years [78], but total loss is unlikely as shown by the well-preserved organics presented by one fossil specimen (ID 12.0.1_7).

Diagenesis of fossil sample ID 12.0.1_7 mirrored those from the historic wet and dry samples. This specimen was collected from the deepest point in the cave, and furthest from the entrance with an overhead environment. Based on current water levels and the caves structure, it is unlikely that drowning and decomposition would result in the deposition of this bone in the location where it was found [5]. It is possible that long-term water level changes did not recede to expose this bone to dry conditions, thus preserving collagen content and structures, however, it is also possible that prior cave diving ventures or other taphonomic processes moved a more modern specimen to the location where it was collected. Direct dating of this specimen may reveal its antiquity.

Collagen integrity and fibre orientation in bone is commonly identified in bone histology and histotaphonomy research through birefringence under polarised light [65, 79]. Data presented here show that despite preserved birefringence in all samples, amide I and II degradation varied from well-preserved to heavily degraded in the fossil specimens. Birefringence was instead associated with degradation of bone structure by microbial attack and staining. Although medical and archaeological studies show complementary histological and biomolecular changes in collagen quantity and structures [80, 81], data emerging from forensics and palaeontological research are not aligned [60, 82].

Although birefringence is associated with collagen fibril structural preservation, it does not necessarily reflect the quantity of collagen in bone [64]. Breakdown of amide results in individual products, cleaved peptide bonds [26], and destruction of structural integrity that theoretically results in lack of birefringence. However, in this study we show a distinct lack of amide I and II across the palaeontological samples coupled with retention of histological birefringence. As the structural backbone of collagen, amide I content by ATR-FTIR can suggest content preservation [28, 83]. Our data contradict the traditional birefringence framework, and further research into the links between birefringence and diagenesis is necessary.

Carbonate dissolution over time depends on depositional environments, however, whole cortex r-FTIRM and ROI s-FTIRM analysis here show few differences between historical wet and dry environments, and the palaeontological assemblage. ROI data produced here fall within the crossover range of well-preserved modern (0.14 – 0.42), archaeological (0.12 – 0.37), and bone from dry karst (0.14 – 0.23) conditions [23, 76, 84]. Bones collected from the underwater caves show less diagenesis across historical and palaeontological timelines compared to those from dry karstic sediments from a Spanish cave where carbonate loss was partially attributed to continuous wetting and drying phases (Del Valle et al 2025), and in Early Neolithic to Middle Palaeolithic fossil rodent bones collected from Moroccan dry cave stratigraphic units representing different climatic periods [85]. Cave waters reflect the minerals of the surrounding rock [86], and at the limestone sites studied here, waters are saturated in calcium carbonates that could limit ionic transfer between hydroxyapatite and water [53]. Reactions are further limited by the mild alkalinity of the cave waters in this study that reduces calcium carbonate solubility [87]. A buffering effect may thus explain the limited carbonate leeching across the wet and palaeontological assemblages, aiding overall preservation in these environments. Our samples suggest that submersion in cave waters may buffer long term carbonate dissolution compared to karstic soils.

In this study, crystallinity in fossil samples was greatest across all bone regions and tissue types compared to those from historic assemblages. This is the standard diagenetic process observed across other archaeological and palaeontological assemblages [15, 23, 40], supporting hydroxyapatite stabilisation and preservation through time [14, 25, 27]. Loss of organics in early diagenesis exposes the biomineral lattice structure to modification, resulting in the replacement or modification of carbonate ions and increased crystallinity, which becomes uncoupled in later diagenesis [34]. Maintenance of carbonate in the fossil specimens here suggests that recrystallisation is occurring in a “closed” system, with localised dissolution and reorganisation of carbonate ions [36]. Substitution with other elements from the surrounding groundwaters cannot be excluded [20, 34], and requires further analysis.

### 4.4 Limitations

Differences in bone type and tissue age should be considered in the interpretation of these data. Mammalian bone structures vary across taxa and limb elements [88, 89], which may alter diagenetic pathways as bone structural units differ in their construction, orientation, and composition. During decay underwater, water will penetrate the three-dimensional blood vessel systems that may impact mineral and organic degradation. As ovicaprid limb bones primarily comprise plexiform bone structures compared to the radial and Haversian systems observed in macropodid limb bones, different water diffusion pathways through vasculature may impact degradation patterns. Further, in bones structures where vessel structures are not contained by a secondary osteon, such as primary canals, water may diffuse freely into the interstitial bone, preventing differential gelatinisation of collagen across the secondary osteons in other regions of the bone. Nevertheless, our analyses across only ovicaprids show differences in biomineral changes in wet environments compared to dry settings. Other biological confounders include an increase in crystallinity with bone tissue and animal age [74, 90], also found in some pathological conditions [59, 91], and biomineral differences between different bone elements [92] which could not be controlled for in this study.

Analysis of archaeological and palaeontological assemblages further restricts interpretations of data, as the ‘true’ depositional histories of bones cannot be known. The historic assemblage analysed is both naturally and culturally deposited (Walker et al., 2026). At Green Waterhole, bones were butchered and thus the time between death of the animal and deposition is not known. Further, some bones were fractured post-mortem, possibly altering the diagenetic pathway. This is however restricted mostly to the dry specimens. Despite these limitations, FTIRM analysis of amide clearly shows different wet and dry decomposition signatures.

## 5 Conclusion

Structural and biomolecular analysis of bone and fossils from different cave environments show that submerged bones follow a different diagenetic pathway to those deposited in dry caves. Unlike previous studies and modelling of aquatic submergence, collagen quantity and structures represented by the amide I and II wavelengths are better preserved in underwater caves compared to dry caves. The stable chemical, thermal, and hydraulic environment in the underwater caves tested here may thus limit the impact of chemical hydrolysis on bones in early diagenesis.

Tissue-specific pathways were observed in the historically-deposited wet bone assemblage, with increased modification of collagen and associated recrystallisation of mineral lattice in the outer bone surfaces. Patterns of external degradation support current theoretical models of preferential hydrolysis in the outer bone layers in early diagenesis. However, the proposed differential impact of collagen chemical hydrolysis, assessed by comparing the presence of secondary osteon microfractures with changes to biomolecular collagen patterns, was not observed in samples from historic or geologic timelines. Instead, changes to bone mineral content and structures were identified between interstitial and secondary osteon bone tissues. Importantly, preservation of bone microstructure, as assessed by birefringence and secondary osteon micro-cracking, did not necessarily correspond to preservation of collagen-associated molecular signatures measured by FTIRM, demonstrating that histological integrity alone is not a reliable indicator of biochemical preservation.

There were few overlaps between the fossil and wet historic bone specimens. In fossils, collagen structures were heavily reduced, crystallinity elevated, and specific bone tissues were similarly modified, suggesting long term modifications override some evidence of early diagenesis. Proportions of carbonate were however similar across wet assemblages, reflecting early diagenesis, and the fossil specimens, representing late diagenesis. Preservation of carbonate, possibly reflecting local cave waters and a ‘closed’ mineral diagenesis system, could highlight a long-term diagenetic trend associated with bones in underwater caves. Further work is required to determine the original depositional environments associated with the fossil assemblage to identify the complete early to late diagenetic pathway associated with bone preservation in underwater caves.

## Acknowledgements

We thank the Burrandies Aboriginal Corporation on whose land this work was conducted, the Department for Environment and Water (DEW) for land access permissions, the Cave Divers Association of Australia specifically Joseph Monks, Damian Bishop, Kelvyn Ball, Steve Trewavas, Hiro Yoshida, Ellyse Klein and Tanya Yarra for supporting this project in the retrieval of historic bone assemblages, Keith Bambery at the Australian Synchrotron for assisting with the beamtime experiment, Aaron Camens, Gavin Prideaux and Carey Burke for access and extracting fossil specimens, Kitrim Dhakal, Vikral Neelesh Vakil, Tanya Smith and Emma Sudron for facilitating SEM and histological processing and imaging, and Leon Manuel and Alan White for their support at the Grifith Analytical Facility. The synchrotron FTIRM measurement was performed on the Infrared Microspectroscopy (IRM) beamline at the Australian Synchrotron, part of ANSTO, through merit-based beamtime access grant (Proposal ID. 23940).

## AI disclosure

An artificial intelligence program was used to assist developing code for R statistics program that visualised quantitative data. All outputs were corroborated and checked by the authors for accuracy.

## Funding information

Australian Research Council (LP210200704), Merit-Based Australian Synchrotron Beamline Access Grant (Proposal ID. 23940), Women Divers Hall of Fame Cecelia Connelly Memorial Graduate Scholarship, JM is supported by an Australian Research Council Future Fellowship (FT240100030), and NS is supported by a National Health and Medical Research Council Investigator grant (2025750).

## Notes

https://zenodo.org/records/20922391

### Competing Interest Statement

The authors have declared no competing interest.

