## Supplementary 1 for "Underwater caves preserve biochemical composition and histological structure in modern and fossil mammalian bones"

**Supplementary 1: Spectral processing, cleaning, and raw data for: *Underwater caves preserve biochemical composition and histological structure in modern and fossil mammalian bones***

Walker, MM, Miszkiewicz, JJ, Rowe, JM, Matheson, CD, Vongsvivut, J, Sims, NA and Louys J

### 1 Data processing and cleaning

Data acquisition integration methods were developed following previously published methodologies adapted to the current assemblage (Lebon et al., 2011, Reiche et al., 2010, Scaggion et al., 2024). Wavenumber absorbance values were collected from baselines generated for each spectra (Table 1).

Python analysis of reflectance FTIRM (r-FTIRM) data supported baseline calculations using minima and maxima on either side of the regional peak (Table 1). However, this was not possible for the s-FTIR data, and thus baselines were determined by previously published literature. As diagenesis indices methodologies use different baseline points, and therefore multiple values for the same wavenumbers were produced (Table 1). For example, Rieche et al (2010) calculate Amide 1 (1660cm^-1^) using a baseline that only covers the Amide 1 peak (1720 – 1590 cm^-1^), whilst Lebon et al. (2010) calculate the same wavenumber using a baseline across the region used to assess Amide I, Amide II and carbonate type v3 region (1800 – 1300 cm^-1^). Values were thus not interchangeable, and used specifically for the diagenesis index associated with each methodology.

Table 1 Absorbance wavenumbers collected and associated baselines

| Vibrational mode and functional group | Wavenumber (cm-1) | r-FTIRM Baseline max range (cm-1) | r-FTIRM Baseline min range (cm-1) | s-FTIRM Wavelength baseline max (cm-1) | s-FTIRM Wavelength baseline min (cm-1) | s-FTIR Reference |
| --- | --- | --- | --- | --- | --- | --- |
| Mineral | | | | | | |
| *v*_3_ PO_4_^3^ (antisymmetric *v*_3_ stretching) | maximum | 1150-1250 | 850-950 | 1200 | 900 | (Lebon et al., 2011) |
| *v*_3_ PO_4_^3^ | 1060 | 1150-1250 | 850-950 | 1200 | 900 | (Lebon et al., 2011) |
| *v*_3_ PO_4_^3^ | 1075 | 1150-1250 | 850-950 | 1200 | 900 | (Lebon et al., 2011) |
| *v*_3_ PO_4_^3^ (non- stoichiometric environment) | 1020 | 1150-1250 | 850-950 | 1175 | 900 | (Reiche et al., 2010) |
| *v*_3_ PO_4_^3^ (stoichiometric environment) | 1030 | 1150-1250 | 850-950 | 1175 | 900 | (Reiche et al., 2010) |
| HPO4 | 1106 | 1150-1250 | 850-950 | 1200 | 900 | (Lebon et al., 2011) |
| *v*_3_ CO_3_ | 1417 | 1700-1800 | 1300-1400 | 1800 | 1300 | (Lebon et al., 2011) |
| Organic | | | | | | |
| Amide II (C-N) stretch and delta (N-H) bend | 1550 | 1700-1800 | 1300-1400 | 1800 | 1300 | (Scaggion et al., 2024) |
| Amide 1 (C=O) stretch | 1660 | 1700-1800 | 1300-1400 | 1800 | 1300 | (Lebon et al., 2011) |
| Amide 1 (C=O) stretch | 1660 | NA | NA | 1720 | 1590 | (Reiche et al., 2010) |
| Amide 1 (reducible cross-links) | 1690 | 1700-1800 | 1300-1400 | 1720 | 1590 | (Reiche et al., 2010) |
| Amide 1 (random coils) | 1645 | 1700-1800 | 1300-1400 | 1720 | 1590 | (Reiche et al., 2010) |

### 1.1 s-FTIR data cleaning

Once exported into Microsoft Excel, thresholds were used to determine poor quality spectrum. Bad quality spectra met one or more of the following absorbance conditions: 1) Full Range value was greater than -50 absorbance, 2) maximum v3PO4 was less than 1060 wavelength (v3PO4), 3) maximum v3PO4 was less than 1660 wavelength (Amide 1), 4) maximum v3PO4 was less than 0.1 absorbance, 5) wavelength 1550 (CO3) was less than 0.05 absorbance, 5) any collected v3PO4 value (Table 1) was less than 0 absorbance. Amide absorbance was not used as a threshold due to the severe general degradation of organics across the fossil assemblage These threshold values were developed based on an assessment of individual spectra and heat maps for each specimen. They consistently identified taphonomic modifications, empty spaces, epoxy, and areas with poor ART contact on the specimen. Once poor-quality spectra were removed, the sub-assemblage datasets (On.Ar, In.Ar) were cross checked for replication against the ROI dataset to ensure data consistency. Total spectra collected for each dataset, and pre- and post-cleaning are reported for each specimen in Table 2.

Table 2 Total spectra collected for each region of interest and sub-groups (In.Ar, On.Ar) before and after cleaning.

|  |  |  | **Total spectra collected** | | | | **Total spectra after cleaning** | | | |
| --- | --- | --- | --- | --- | --- | --- | --- | --- | --- | --- |
| **ANSTO #** | **Sample name** | **Environment** | **ROI** | **In.Ar** | **On.Ar** | **Total** | **ROI** | **In.Ar** | **On.Ar** | **Total** |
| MTGIRM16 | 13.0.2_1_On3 | Fossil | 625 | 233 | 296 | 1154 | 605 | 229 | 280 | 1114 |
| MTGIRM16 | 13.0.2_1_On4 | Fossil | 625 | 433 | 142 | 1200 | 485 | 392 | 51 | 928 |
| MTGIRM21 | 32_6_On3 | Fossil | 625 | 300 | 270 | 1195 | 491 | 293 | 143 | 927 |
| MTGIRM21 | 32_6_On4 | Fossil | 625 | 310 | 276 | 1211 | 330 | 272 | 28 | 630 |
| MTGIRM23 | 4.0.1_1_On1 | Fossil | 625 | 243 | 308 | 1176 | 524 | 243 | 209 | 976 |
| MTGIRM23 | 4.0.1_1_On2 | Fossil | 625 | 321 | 187 | 1133 | 564 | 272 | 180 | 1016 |
| MTGIRM19 | 4.1.5_1_On3 | Fossil | 625 | 370 | 188 | 1183 | 392 | 151 | 180 | 723 |
| MTGIRM19 | 4.1.5_1_On4 | Fossil | 625 | 299 | 223 | 1147 | 603 | 287 | 217 | 1107 |
| MTGIRM15 | 5.0.32_5_On3 | Fossil | 625 | 334 | 202 | 1161 | 518 | 326 | 119 | 963 |
| MTGIRM15 | 5.0.32_5_On4 | Fossil | 625 |  |  | 625 | 573 |  |  | 573 |
| MTGIRM17 | 7.13.2_1_On3 | Fossil | 625 | 263 | 298 | 1186 | 506 | 214 | 240 | 960 |
| MTGIRM17 | 7.13.2_1_On4 | Fossil | 625 | 293 | 217 | 1135 | 582 | 270 | 197 | 1049 |
| MTGIRM18 | 7.8.4_1_On1 | Fossil | 625 | 373 | 178 | 1176 | 201 | 30 | 141 | 372 |
| MTGIRM18 | 7.8.4_1_On2 | Fossil | 625 | 326 | 216 | 1167 | 430 | 324 | 37 | 791 |
| MTGIRM03 | GH_10_On4 | Wet | 625 | 220 | 313 | 1158 | 556 | 212 | 268 | 1036 |
| MTGIRM14 | GH_18_On3 | Wet | 625 | 231 | 306 | 1162 | 531 | 191 | 261 | 983 |
| MTGIRM14 | GH_18_On4 | Wet | 625 | 375 | 182 | 1182 | 557 | 351 | 143 | 1051 |
| MTGIRM04 | GH_19_On1 | Wet | 625 | 240 | 321 | 1186 | 593 | 229 | 304 | 1126 |
| MTGIRM04 | GH_19_On2 | Wet | 625 | 257 | 313 | 1195 | 585 | 239 | 291 | 1115 |
| MTGIRM13 | GH_4_On1 | Wet | 625 | 340 | 194 | 1159 | 512 | 276 | 150 | 938 |
| MTGIRM13 | GH_4_On2 | Wet | 625 | 382 | 209 | 1216 | 240 | 122 | 97 | 459 |
| MTGIRM02 | GH_9_On1 | Wet | 625 | 197 | 333 | 1155 | 588 | 188 | 321 | 1097 |
| MTGIRM02 | GH_9_On2 | Wet | 625 | 312 | 227 | 1164 | 552 | 278 | 197 | 1027 |
| MTGIRM05 | GWW_4_On1 | Wet | 625 | 211 | 346 | 1182 | 594 | 203 | 326 | 1123 |
| MTGIRM05 | GWW_4_On2 | Wet | 625 | 228 | 272 | 1125 | 297 | 15 | 220 | 532 |
| MTGIRM01 | GWW_61_On1 | Wet | 625 | 217 | 356 | 1198 | 589 | 198 | 340 | 1127 |
| MTGIRM06 | GWW_76_On1 | Wet | 625 | 231 | 249 | 1105 | 435 | 119 | 229 | 783 |
| MTGIRM06 | GWW_76_On2 | Wet | 625 | 329 | 190 | 1144 | 546 | 309 | 186 | 1041 |
| MTGIRM08 | GWS_02_On1 | Dry | 625 | 113 | 210 | 948 | 551 | 110 | 185 | 846 |
| MTGIRM08 | GWS_02_On2 | Dry | 625 | 60 | 469 | 1154 | 585 | 60 | 459 | 1104 |
| MTGIRM09 | GWS_19_On1 | Dry | 625 | 157 | 393 | 1175 | 611 | 153 | 386 | 1150 |
| MTGIRM09 | GWS_19_On2 | Dry | 625 | 315 | 268 | 1208 | 571 | 271 | 264 | 1106 |
| MTGIRM10 | GWS_47_On1 | Dry | 625 | 222 | 341 | 1188 | 347 | 129 | 188 | 664 |
| MTGIRM10 | GWS_47_On2 | Dry | 625 | 300 | 269 | 1194 | 272 | 38 | 223 | 533 |
| MTGIRM11 | GWS_48_On1 | Dry | 625 | 385 | 206 | 1216 | 589 | 375 | 182 | 1146 |
| MTGIRM11 | GWS_48_On2 | Dry | 625 | 213 | 301 | 1139 | 583 | 209 | 269 | 1061 |
| MTGIRM07 | GWS_60_On3 | Dry | 625 | 237 | 333 | 1195 | 602 | 218 | 331 | 1151 |
| MTGIRM07 | GWS_60_On4 | Dry | 625 | 297 | 266 | 1188 | 602 | 290 | 257 | 1149 |
| MTGIRM12 | GWS_69_On4 | Dry | 625 | 418 | 171 | 1214 | 594 | 407 | 151 | 1152 |
| **TOTAL** | | | **24375** | **10585** | **10039** | **44999** | **19886** | **8493** | **8250** | **36629** |
| **Historic Wet (N=14)** | | | 8750 | 3370 | 3811 | 16331 | 7175 | 2930 | 3333 | 13438 |
| **Historic Dry (N=11)** | | | 6875 | 2717 | 3227 | 12819 | 5907 | 2260 | 2895 | 11062 |
| **Fossil (N=14)** | | | 8750 | 4098 | 3001 | 15849 | 6804 | 3303 | 2022 | 12129 |

### 2 Wavelength data

Raw wavelength data for each group were extracted to determine diagenetic index values. Tables 3 and 4 summarise wavelength data for bone tissue regions across environmental deposition conditions using r-FTIRM methods. Tables 5 and 6 summarise wavelength data for the ROI area and sub-assemblages (In.Ar and On.Ar) across environmental deposition conditions using s-FTIRM methods.

### 2.1 r-FTIRM wavelength data summaries

Table 3 Raw spectral absorbance data for carbonate and phosphate wavelengths collected across region of interest datasets from r-FTIRM methods

| **Wavelength cm^-1^** | | **Sub-periosteal** | | | **Mid-cortical** | | | **Sub-endosteal** | | |
| --- | --- | --- | --- | --- | --- | --- | --- | --- | --- | --- |
|  |  | **Wet** | **Dry** | **Fossil** | **Wet** | **Dry** | **Fossil** | **Wet** | **Dry** | **Fossil** |
| 1415  (*v*_3_ CO_3_) | x̄ | 0.208 | 0.259 | 0.292 | 0.252 | 0.271 | 0.292 | 0.176 | 0.206 | 0.227 |
|  | σ | 0.085 | 0.074 | 0.109 | 0.037 | 0.039 | 0.086 | 0.067 | 0.060 | 0.080 |
|  | Min | 0.045 | 0.050 | 0.046 | 0.073 | 0.112 | 0.059 | 0.031 | 0.043 | 0.042 |
|  | Max | 0.361 | 0.358 | 0.486 | 0.317 | 0.361 | 0.458 | 0.295 | 0.349 | 0.410 |
| Maximum  (*v*_3_ PO_4_^3^) | x̄ | 0.979 | 1.090 | 1.431 | 1.137 | 1.195 | 1.548 | 0.860 | 1.016 | 1.203 |
|  | σ | 0.387 | 0.290 | 0.507 | 0.191 | 0.170 | 0.432 | 0.266 | 0.246 | 0.391 |
|  | Min | 0.300 | 0.302 | 0.305 | 0.346 | 0.476 | 0.309 | 0.301 | 0.316 | 0.300 |
|  | Max | 1.641 | 1.496 | 2.145 | 1.584 | 1.550 | 2.331 | 1.329 | 1.616 | 1.917 |
| 1106  (HPO_4_) | x̄ | 0.601 | 0.703 | 0.712 | 0.690 | 0.721 | 0.768 | 0.534 | 0.599 | 0.655 |
|  | σ | 0.211 | 0.157 | 0.219 | 0.084 | 0.104 | 0.193 | 0.142 | 0.122 | 0.175 |
|  | Min | 0.150 | 0.209 | 0.171 | 0.249 | 0.351 | 0.184 | 0.144 | 0.197 | 0.163 |
|  | Max | 1.048 | 0.952 | 1.194 | 0.882 | 0.994 | 1.084 | 0.837 | 0.991 | 0.965 |
| 1060  (*v*_3_ PO_4_^3^) | x̄ | 0.874 | 0.997 | 1.302 | 1.004 | 1.059 | 1.397 | 0.763 | 0.897 | 1.108 |
|  | σ | 0.330 | 0.245 | 0.445 | 0.146 | 0.141 | 0.374 | 0.214 | 0.192 | 0.341 |
|  | Min | 0.219 | 0.281 | 0.287 | 0.325 | 0.469 | 0.262 | 0.231 | 0.296 | 0.198 |
|  | Max | 1.478 | 1.332 | 2.071 | 1.359 | 1.382 | 1.985 | 1.296 | 1.438 | 1.720 |
| 1075  (*v*_3_ PO_4_^3^) | x̄ | 0.753 | 0.863 | 1.050 | 0.863 | 0.902 | 1.121 | 0.655 | 0.759 | 0.900 |
|  | σ | 0.272 | 0.200 | 0.343 | 0.116 | 0.122 | 0.289 | 0.175 | 0.155 | 0.261 |
|  | Min | 0.192 | 0.251 | 0.253 | 0.308 | 0.424 | 0.266 | 0.188 | 0.251 | 0.192 |
|  | Max | 1.250 | 1.130 | 1.717 | 1.159 | 1.200 | 1.563 | 1.150 | 1.238 | 1.356 |
| 1020  (*v*_3_ PO_4_^3^) | x̄ | 0.686 | 0.773 | 0.748 | 0.863 | 0.936 | 0.774 | 0.629 | 0.757 | 0.599 |
|  | σ | 0.308 | 0.252 | 0.338 | 0.181 | 0.158 | 0.325 | 0.252 | 0.227 | 0.313 |
|  | Min | 0.051 | 0.092 | 0.006 | 0.194 | 0.296 | 0.014 | 0.071 | 0.082 | 0.020 |
|  | Max | 1.228 | 1.173 | 1.429 | 1.249 | 1.251 | 1.429 | 1.038 | 1.298 | 1.373 |
| 1030  (*v*_3_ PO_4_^3^) | x̄ | 0.904 | 1.006 | 1.176 | 1.085 | 1.150 | 1.250 | 0.800 | 0.959 | 0.943 |
|  | σ | 0.384 | 0.297 | 0.458 | 0.203 | 0.176 | 0.401 | 0.282 | 0.261 | 0.388 |
|  | Min | 0.159 | 0.191 | 0.099 | 0.275 | 0.397 | 0.055 | 0.160 | 0.191 | 0.033 |
|  | Max | 1.590 | 1.418 | 1.910 | 1.543 | 1.510 | 1.994 | 1.273 | 1.574 | 1.716 |

Table 4 Raw spectral absorbance data for carbonate and phosphate wavelengths collected across region of interest datasets from r-FTIRM methods

| **Wavelength cm^-1^** | | **Sub-periosteal** | | | **Mid-cortical** | | | **Sub-endosteal** | | |
| --- | --- | --- | --- | --- | --- | --- | --- | --- | --- | --- |
|  |  | **Wet** | **Dry** | **Fossil** | **Wet** | **Dry** | **Fossil** | **Wet** | **Dry** | **Fossil** |
| 1660 (Amide I) | x̄ | 0.309 | 0.261 | 0.082 | 0.467 | 0.444 | 0.083 | 0.310 | 0.236 | 0.072 |
|  | σ | 0.163 | 0.150 | 0.066 | 0.064 | 0.071 | 0.128 | 0.157 | 0.175 | 0.102 |
|  | Min | 0.034 | 0.024 | 0.001 | 0.148 | 0.207 | 0.002 | 0.041 | 0.023 | 0.001 |
|  | Max | 0.565 | 0.585 | 0.390 | 0.611 | 0.614 | 0.517 | 0.583 | 0.549 | 0.533 |
| 1550 (Amide II) | x̄ | 0.159 | 0.147 | 0.079 | 0.213 | 0.197 | 0.085 | 0.146 | 0.127 | 0.082 |
|  | σ | 0.069 | 0.054 | 0.037 | 0.030 | 0.028 | 0.052 | 0.062 | 0.066 | 0.144 |
|  | Min | 0.031 | 0.026 | 0.002 | 0.062 | 0.098 | 0.013 | 0.017 | 0.026 | 0.001 |
|  | Max | 0.263 | 0.247 | 0.220 | 0.269 | 0.250 | 0.258 | 0.251 | 0.241 | 2.199 |
| 1645 (Amide I) | x̄ | 0.272 | 0.236 | 0.084 | 0.403 | 0.381 | 0.082 | 0.271 | 0.208 | 0.073 |
|  | σ | 0.138 | 0.126 | 0.061 | 0.054 | 0.059 | 0.110 | 0.134 | 0.149 | 0.093 |
|  | Min | 0.040 | 0.031 | 0.001 | 0.128 | 0.161 | 0.003 | 0.034 | 0.028 | 0.002 |
|  | Max | 0.484 | 0.506 | 0.361 | 0.524 | 0.504 | 0.444 | 0.497 | 0.490 | 0.568 |
| 1690 (Amide I) | x̄ | 0.114 | 0.108 | 0.042 | 0.167 | 0.159 | 0.040 | 0.109 | 0.091 | 0.036 |
|  | σ | 0.057 | 0.049 | 0.029 | 0.023 | 0.027 | 0.044 | 0.051 | 0.059 | 0.044 |
|  | Min | 0.000 | 0.010 | 0.000 | 0.061 | 0.086 | 0.000 | 0.010 | 0.014 | 0.000 |
|  | Max | 0.211 | 0.212 | 0.170 | 0.226 | 0.227 | 0.201 | 0.210 | 0.210 | 0.372 |

### 2.2 ATR-sFTIR wavelength data summaries

Table 5 Raw spectral absorbance data for carbonate and phosphate wavelengths collected across region of interest (ROI), interstitial area (In.Ar) and osteonal area (On.Ar) datasets from ATR-sFTIR methods

| **Wavelength cm^-1^** | | **Sub-periosteal** | | | **Mid-cortical** | | | **Sub-endosteal** | | |
| --- | --- | --- | --- | --- | --- | --- | --- | --- | --- | --- |
|  |  | **Wet** | **Dry** | **Fossil** | **Wet** | **Dry** | **Fossil** | **Wet** | **Dry** | **Fossil** |
| 1417  (*v*_3_ CO_3_) | x̄ | 0.098 | 0.111 | 0.105 | 0.101 | 0.112 | 0.106 | 0.099 | 0.113 | 0.101 |
|  | σ | 0.029 | 0.027 | 0.033 | 0.030 | 0.026 | 0.030 | 0.025 | 0.026 | 0.038 |
|  | Min | 0.010 | 0.010 | 0.000 | 0.011 | 0.010 | 0.000 | 0.014 | 0.011 | 0.000 |
|  | Max | 0.179 | 0.190 | 0.352 | 0.179 | 0.190 | 0.187 | 0.175 | 0.186 | 0.352 |
| Maximum  (*v*_3_ PO_4_^3^) | x̄ | 0.398 | 0.432 | 0.493 | 0.426 | 0.440 | 0.508 | 0.388 | 0.428 | 0.491 |
|  | σ | 0.115 | 0.120 | 0.188 | 0.118 | 0.119 | 0.198 | 0.107 | 0.119 | 0.182 |
|  | Min | 0.100 | 0.102 | 0.103 | 0.116 | 0.141 | 0.103 | 0.100 | 0.102 | 0.103 |
|  | Max | 0.827 | 0.877 | 1.106 | 0.827 | 0.877 | 1.106 | 0.756 | 0.833 | 0.983 |
| 1106  (HPO_4_) | x̄ | 0.086 | 0.093 | 0.077 | 0.078 | 0.096 | 0.084 | 0.093 | 0.091 | 0.073 |
|  | σ | 0.036 | 0.038 | 0.036 | 0.037 | 0.040 | 0.032 | 0.034 | 0.036 | 0.037 |
|  | Min | 0.000 | 0.000 | 0.000 | 0.000 | 0.000 | 0.000 | 0.000 | 0.000 | 0.000 |
|  | Max | 0.194 | 0.199 | 0.210 | 0.194 | 0.199 | 0.184 | 0.193 | 0.195 | 0.210 |
| 1060  (*v*_3_ PO_4_^3^) | x̄ | 0.206 | 0.221 | 0.238 | 0.202 | 0.229 | 0.252 | 0.213 | 0.214 | 0.224 |
|  | σ | 0.055 | 0.058 | 0.061 | 0.060 | 0.062 | 0.056 | 0.049 | 0.054 | 0.060 |
|  | Min | 0.024 | 0.018 | 0.022 | 0.027 | 0.021 | 0.042 | 0.025 | 0.018 | 0.022 |
|  | Max | 0.373 | 0.396 | 0.441 | 0.373 | 0.386 | 0.430 | 0.344 | 0.396 | 0.441 |
| 1075  (*v*_3_ PO_4_^3^) | x̄ | 0.154 | 0.167 | 0.160 | 0.148 | 0.173 | 0.172 | 0.162 | 0.161 | 0.150 |
|  | σ | 0.049 | 0.052 | 0.051 | 0.053 | 0.056 | 0.047 | 0.044 | 0.048 | 0.050 |
|  | Min | 0.003 | 0.000 | 0.009 | 0.003 | 0.001 | 0.017 | 0.007 | 0.000 | 0.009 |
|  | Max | 0.291 | 0.315 | 0.318 | 0.291 | 0.308 | 0.302 | 0.291 | 0.315 | 0.318 |
| 1020  (*v*_3_ PO_4_^3^) | x̄ | 0.397 | 0.429 | 0.480 | 0.424 | 0.435 | 0.495 | 0.387 | 0.426 | 0.479 |
|  | σ | 0.115 | 0.119 | 0.194 | 0.116 | 0.118 | 0.204 | 0.108 | 0.118 | 0.187 |
|  | Min | 0.097 | 0.099 | 0.083 | 0.097 | 0.142 | 0.099 | 0.101 | 0.099 | 0.083 |
|  | Max | 0.804 | 0.862 | 1.099 | 0.804 | 0.862 | 1.099 | 0.755 | 0.816 | 0.985 |
| 1030  (*v*_3_ PO_4_^3^) | x̄ | 0.373 | 0.398 | 0.463 | 0.390 | 0.405 | 0.478 | 0.369 | 0.395 | 0.460 |
|  | σ | 0.091 | 0.093 | 0.156 | 0.091 | 0.091 | 0.163 | 0.086 | 0.092 | 0.152 |
|  | Min | 0.092 | 0.090 | 0.093 | 0.110 | 0.120 | 0.095 | 0.103 | 0.090 | 0.093 |
|  | Max | 0.690 | 0.743 | 0.952 | 0.685 | 0.743 | 0.952 | 0.684 | 0.695 | 0.876 |

Table 6 Raw spectral absorbance data for amide region wavelengths collected across region of interest datasets from s-FTIRM methods

| **Wavelength cm^-1^** | | **ROI** | | | **In.Ar** | | | **On.Ar** | | |
| --- | --- | --- | --- | --- | --- | --- | --- | --- | --- | --- |
|  |  | **Wet** | **Dry** | **Fossil** | **Wet** | **Dry** | **Fossil** | **Wet** | **Dry** | **Fossil** |
| 1660  (Amide I)  (L) | x̄ | 0.112 | 0.119 | 0.012 | 0.110 | 0.118 | 0.014 | 0.119 | 0.123 | 0.010 |
|  | σ | 0.035 | 0.033 | 0.012 | 0.036 | 0.032 | 0.012 | 0.032 | 0.032 | 0.012 |
|  | Min | 0.000 | 0.009 | 0.000 | 0.000 | 0.009 | 0.000 | 0.017 | 0.014 | 0.000 |
|  | Max | 0.214 | 0.223 | 0.173 | 0.196 | 0.202 | 0.173 | 0.214 | 0.223 | 0.063 |
| 1660  (Amide I)  (R) | x̄ | 0.094 | 0.101 | 0.003 | 0.091 | 0.100 | 0.004 | 0.101 | 0.105 | 0.002 |
|  | σ | 0.028 | 0.025 | 0.006 | 0.028 | 0.024 | 0.007 | 0.027 | 0.024 | 0.005 |
|  | Min | 0.001 | 0.007 | 0.000 | 0.001 | 0.007 | 0.000 | 0.014 | 0.016 | 0.000 |
|  | Max | 0.169 | 0.158 | 0.151 | 0.148 | 0.158 | 0.151 | 0.169 | 0.156 | 0.033 |
| 1550  (Amide II) | x̄ | 0.082 | 0.084 | 0.024 | 0.082 | 0.083 | 0.023 | 0.086 | 0.087 | 0.025 |
|  | σ | 0.027 | 0.025 | 0.018 | 0.030 | 0.024 | 0.016 | 0.022 | 0.025 | 0.021 |
|  | Min | 0.000 | 0.000 | 0.000 | 0.000 | 0.000 | 0.000 | 0.005 | 0.005 | 0.000 |
|  | Max | 0.182 | 0.169 | 0.129 | 0.182 | 0.166 | 0.129 | 0.154 | 0.169 | 0.102 |
| 1690  (Amide I) | x̄ | 0.028 | 0.029 | -0.001 | 0.028 | 0.028 | 0.000 | 0.030 | 0.030 | 0.000 |
|  | σ | 0.011 | 0.011 | 0.003 | 0.011 | 0.011 | 0.003 | 0.011 | 0.011 | 0.003 |
|  | Min | 0.000 | 0.000 | 0.000 | 0.000 | 0.000 | 0.000 | 0.000 | 0.000 | 0.000 |
|  | Max | 0.073 | 0.076 | 0.045 | 0.066 | 0.060 | 0.045 | 0.073 | 0.076 | 0.014 |
| 1645  (Amide I) | x̄ | 0.089 | 0.095 | 0.004 | 0.086 | 0.095 | 0.005 | 0.096 | 0.100 | 0.003 |
|  | σ | 0.027 | 0.023 | 0.006 | 0.027 | 0.022 | 0.007 | 0.026 | 0.022 | 0.004 |
|  | Min | 0.002 | 0.007 | 0.000 | 0.002 | 0.007 | 0.000 | 0.013 | 0.017 | 0.000 |
|  | Max | 0.153 | 0.143 | 0.191 | 0.141 | 0.143 | 0.191 | 0.153 | 0.139 | 0.030 |

### 3 Index data summary per sample

Diagenetic indices were developed based on wavelength absorbance data following ratios provided in Table 1. Table 7 and 8 provide the mean and standard deviation data collected using r-FTIRM methods for each specimen and bone region whilst Tables 9-12 summarise the same indices for ROI, In.Ar and On.Ar collected using s-FTIRM techniques.

### 3.1 r-FTIRM data

Table 7 Mineral diagenetic index data per sample and bone region collected by r-FTIRM methods.

|  |  |  |  | **C/P** | | **HPO_4_/P** | | **CI1** | | **CI2** | |
| --- | --- | --- | --- | --- | --- | --- | --- | --- | --- | --- | --- |
| **Sample ID** | **Bone region** | **Environment** | **n. pixels** | **x̄** | **σ** | **x̄** | **σ** | **x̄** | **σ** | **x̄** | **σ** |
| GWS02 | Sub-endosteal | Dry | 29 | 0.215 | 0.017 | 0.583 | 0.065 | 1.248 | 0.047 | 1.204 | 0.035 |
| GWS19 | Sub-endosteal | Dry | 51 | 0.227 | 0.009 | 0.609 | 0.071 | 1.285 | 0.105 | 1.175 | 0.030 |
| GWS47 | Sub-endosteal | Dry | 41 | 0.183 | 0.015 | 0.683 | 0.089 | 1.295 | 0.090 | 1.129 | 0.032 |
| GWS48 | Sub-endosteal | Dry | 43 | 0.209 | 0.020 | 0.598 | 0.078 | 1.280 | 0.098 | 1.165 | 0.038 |
| GWS60 | Sub-endosteal | Dry | 40 | 0.163 | 0.008 | 0.513 | 0.099 | 1.342 | 0.203 | 1.246 | 0.044 |
| GWS69 | Sub-endosteal | Dry | 16 | 0.225 | 0.013 | 0.688 | 0.099 | 1.293 | 0.196 | 1.130 | 0.052 |
| GWS02 | Mid-cortical | Dry | 42 | 0.228 | 0.006 | 0.656 | 0.038 | 1.261 | 0.032 | 1.161 | 0.017 |
| GWS19 | Mid-cortical | Dry | 60 | 0.227 | 0.007 | 0.606 | 0.045 | 1.299 | 0.041 | 1.180 | 0.018 |
| GWS47 | Mid-cortical | Dry | 48 | 0.228 | 0.007 | 0.624 | 0.070 | 1.218 | 0.058 | 1.171 | 0.028 |
| GWS48 | Mid-cortical | Dry | 48 | 0.219 | 0.007 | 0.572 | 0.026 | 1.215 | 0.026 | 1.178 | 0.010 |
| GWS60 | Mid-cortical | Dry | 48 | 0.218 | 0.007 | 0.527 | 0.039 | 1.179 | 0.024 | 1.210 | 0.021 |
| GWS69 | Mid-cortical | Dry | 60 | 0.238 | 0.006 | 0.651 | 0.041 | 1.218 | 0.032 | 1.151 | 0.014 |
| GWS02 | Sub-periosteal | Dry | 46 | 0.234 | 0.009 | 0.693 | 0.062 | 1.350 | 0.083 | 1.151 | 0.026 |
| GWS19 | Sub-periosteal | Dry | 70 | 0.240 | 0.008 | 0.648 | 0.043 | 1.366 | 0.057 | 1.171 | 0.020 |
| GWS47 | Sub-periosteal | Dry | 48 | 0.235 | 0.034 | 0.772 | 0.095 | 1.425 | 0.127 | 1.095 | 0.034 |
| GWS48 | Sub-periosteal | Dry | 53 | 0.214 | 0.019 | 0.646 | 0.095 | 1.298 | 0.146 | 1.135 | 0.040 |
| GWS60 | Sub-periosteal | Dry | 55 | 0.241 | 0.007 | 0.587 | 0.081 | 1.263 | 0.144 | 1.176 | 0.035 |
| GWS69 | Sub-periosteal | Dry | 68 | 0.249 | 0.006 | 0.647 | 0.034 | 1.285 | 0.070 | 1.152 | 0.015 |
| 12.0.1_7 | Sub-endosteal | Fossil | 24 | 0.195 | 0.017 | 0.560 | 0.048 | 1.244 | 0.062 | 1.187 | 0.013 |
| 13.0.1_1 | Sub-endosteal | Fossil | 13 | 0.194 | 0.012 | 0.648 | 0.050 | 1.435 | 0.110 | 1.185 | 0.033 |
| 32_11 | Sub-endosteal | Fossil | 26 | 0.166 | 0.010 | 0.498 | 0.062 | 1.592 | 0.152 | 1.288 | 0.025 |
| 32_6 | Sub-endosteal | Fossil | 26 | 0.207 | 0.020 | 0.508 | 0.082 | 1.668 | 1.079 | 1.256 | 0.048 |
| 4.0.1_1 | Sub-endosteal | Fossil | 60 | 0.203 | 0.009 | 0.576 | 0.080 | 1.694 | 0.158 | 1.203 | 0.049 |
| 4.1.5_1 | Sub-endosteal | Fossil | 24 | 0.170 | 0.005 | 0.614 | 0.051 | 2.388 | 0.638 | 1.223 | 0.039 |
| 5.0.32_5 | Sub-endosteal | Fossil | 18 | 0.160 | 0.015 | 0.611 | 0.108 | 2.199 | 0.758 | 1.191 | 0.079 |
| 7.13.2_1 | Sub-endosteal | Fossil | 24 | 0.171 | 0.010 | 0.579 | 0.103 | 1.803 | 0.251 | 1.238 | 0.041 |
| 7.8.4_1 | Sub-endosteal | Fossil | 18 | 0.217 | 0.086 | 0.542 | 0.105 | 1.649 | 0.220 | 1.213 | 0.064 |
| 12.0.1_7 | Mid-cortical | Fossil | 24 | 0.200 | 0.005 | 0.530 | 0.039 | 1.197 | 0.036 | 1.183 | 0.015 |
| 13.0.1_1 | Mid-cortical | Fossil | 23 | 0.198 | 0.014 | 0.671 | 0.080 | 1.727 | 0.999 | 1.134 | 0.058 |
| 32_11 | Mid-cortical | Fossil | 30 | 0.185 | 0.007 | 0.452 | 0.025 | 1.600 | 0.107 | 1.283 | 0.013 |
| 32_6 | Mid-cortical | Fossil | 30 | 0.199 | 0.009 | 0.484 | 0.049 | 1.629 | 0.149 | 1.265 | 0.030 |
| 4.0.1_1 | Mid-cortical | Fossil | 66 | 0.200 | 0.006 | 0.478 | 0.019 | 1.628 | 0.068 | 1.258 | 0.009 |
| 4.1.5_1 | Mid-cortical | Fossil | 24 | 0.155 | 0.005 | 0.563 | 0.026 | 2.942 | 0.229 | 1.219 | 0.018 |
| 5.0.32_5 | Mid-cortical | Fossil | 24 | 0.174 | 0.007 | 0.486 | 0.026 | 1.708 | 0.207 | 1.260 | 0.015 |
| 7.13.2_1 | Mid-cortical | Fossil | 24 | 0.177 | 0.011 | 0.552 | 0.056 | 1.840 | 0.193 | 1.226 | 0.037 |
| 7.8.4_1 | Mid-cortical | Fossil | 24 | 0.190 | 0.005 | 0.470 | 0.025 | 1.605 | 0.097 | 1.265 | 0.015 |
| 12.0.1_7 | Sub-periosteal | Fossil | 27 | 0.178 | 0.017 | 0.574 | 0.080 | 1.382 | 0.131 | 1.173 | 0.037 |
| 13.0.1_1 | Sub-periosteal | Fossil | 23 | 0.243 | 0.018 | 0.604 | 0.064 | 1.432 | 0.129 | 1.189 | 0.050 |
| 32_11 | Sub-periosteal | Fossil | 31 | 0.195 | 0.022 | 0.558 | 0.114 | 1.663 | 0.806 | 1.205 | 0.076 |
| 32_6 | Sub-periosteal | Fossil | 33 | 0.218 | 0.010 | 0.464 | 0.080 | 1.474 | 0.183 | 1.264 | 0.052 |
| 4.0.1_1 | Sub-periosteal | Fossil | 73 | 0.209 | 0.012 | 0.524 | 0.077 | 1.771 | 0.321 | 1.222 | 0.054 |
| 4.1.5_1 | Sub-periosteal | Fossil | 28 | 0.176 | 0.009 | 0.555 | 0.049 | 2.424 | 0.259 | 1.219 | 0.038 |
| 5.0.32_5 | Sub-periosteal | Fossil | 28 | 0.190 | 0.008 | 0.486 | 0.034 | 1.665 | 0.166 | 1.254 | 0.018 |
| 7.13.2_1 | Sub-periosteal | Fossil | 28 | 0.217 | 0.007 | 0.510 | 0.061 | 1.528 | 0.116 | 1.257 | 0.032 |
| 7.8.4_1 | Sub-periosteal | Fossil | 28 | 0.214 | 0.005 | 0.421 | 0.048 | 1.426 | 0.121 | 1.281 | 0.029 |
| GH04 | Sub-endosteal | Wet | 74 | 0.181 | 0.042 | 0.773 | 0.074 | 1.501 | 0.141 | 1.117 | 0.030 |
| GH10 | Sub-endosteal | Wet | 31 | 0.209 | 0.011 | 0.589 | 0.060 | 1.293 | 0.056 | 1.207 | 0.032 |
| GH18 | Sub-endosteal | Wet | 22 | 0.202 | 0.027 | 0.653 | 0.089 | 1.357 | 0.102 | 1.119 | 0.034 |
| GH19 | Sub-endosteal | Wet | 44 | 0.198 | 0.025 | 0.574 | 0.079 | 1.223 | 0.053 | 1.185 | 0.034 |
| GH9 | Sub-endosteal | Wet | 53 | 0.220 | 0.019 | 0.689 | 0.065 | 1.334 | 0.071 | 1.154 | 0.026 |
| GWW4 | Sub-endosteal | Wet | 29 | 0.173 | 0.011 | 0.637 | 0.064 | 1.248 | 0.066 | 1.158 | 0.027 |
| GWW61 | Sub-endosteal | Wet | 48 | 0.222 | 0.008 | 0.510 | 0.057 | 1.181 | 0.056 | 1.208 | 0.028 |
| GWW76 | Sub-endosteal | Wet | 32 | 0.205 | 0.006 | 0.607 | 0.043 | 1.223 | 0.038 | 1.151 | 0.019 |
| GH04 | Mid-cortical | Wet | 90 | 0.258 | 0.009 | 0.675 | 0.051 | 1.306 | 0.047 | 1.142 | 0.020 |
| GH10 | Mid-cortical | Wet | 42 | 0.214 | 0.011 | 0.600 | 0.071 | 1.288 | 0.033 | 1.189 | 0.032 |
| GH18 | Mid-cortical | Wet | 30 | 0.203 | 0.006 | 0.560 | 0.024 | 1.239 | 0.024 | 1.182 | 0.011 |
| GH19 | Mid-cortical | Wet | 48 | 0.194 | 0.012 | 0.563 | 0.043 | 1.262 | 0.041 | 1.173 | 0.017 |
| GH9 | Mid-cortical | Wet | 65 | 0.251 | 0.008 | 0.694 | 0.049 | 1.306 | 0.051 | 1.136 | 0.016 |
| GWW4 | Mid-cortical | Wet | 34 | 0.207 | 0.009 | 0.653 | 0.051 | 1.223 | 0.064 | 1.146 | 0.024 |
| GWW61 | Mid-cortical | Wet | 48 | 0.215 | 0.005 | 0.550 | 0.022 | 1.185 | 0.018 | 1.178 | 0.010 |
| GWW76 | Mid-cortical | Wet | 36 | 0.189 | 0.007 | 0.525 | 0.023 | 1.247 | 0.025 | 1.192 | 0.011 |
| GH04 | Sub-periosteal | Wet | 91 | 0.221 | 0.023 | 0.728 | 0.101 | 1.412 | 0.142 | 1.110 | 0.041 |
| GH10 | Sub-periosteal | Wet | 43 | 0.233 | 0.013 | 0.661 | 0.058 | 1.463 | 0.264 | 1.172 | 0.037 |
| GH18 | Sub-periosteal | Wet | 35 | 0.218 | 0.025 | 0.596 | 0.096 | 1.349 | 0.133 | 1.156 | 0.045 |
| GH19 | Sub-periosteal | Wet | 52 | 0.195 | 0.007 | 0.566 | 0.074 | 1.370 | 0.174 | 1.172 | 0.030 |
| GH9 | Sub-periosteal | Wet | 59 | 0.211 | 0.039 | 0.673 | 0.092 | 1.301 | 0.096 | 1.129 | 0.026 |
| GWW4 | Sub-periosteal | Wet | 38 | 0.237 | 0.014 | 0.687 | 0.059 | 1.270 | 0.109 | 1.133 | 0.022 |
| GWW61 | Sub-periosteal | Wet | 54 | 0.214 | 0.014 | 0.607 | 0.062 | 1.278 | 0.115 | 1.149 | 0.028 |
| GWW76 | Sub-periosteal | Wet | 42 | 0.163 | 0.017 | 0.476 | 0.047 | 1.300 | 0.123 | 1.231 | 0.027 |

Table 8 Amide region diagenetic index data per sample and bone region collected by r-FTIRM methods.

|  |  |  |  | **AmI/P** | | **AmII/P** | | **AmI/II** | | **Collagen Integrity** | | **Random coils** | |
| --- | --- | --- | --- | --- | --- | --- | --- | --- | --- | --- | --- | --- | --- |
| **Sample ID** | **Bone region** | **Environment** | **n. pixels** | **x̄** | **σ** | **x̄** | **σ** | **x̄** | **σ** | **x̄** | **σ** | **x̄** | **σ** |
| GWS02 | Sub-endosteal | Dry | 29 | 0.326 | 0.088 | 0.157 | 0.024 | 2.030 | 0.348 | 0.368 | 0.066 | 0.884 | 0.054 |
| GWS19 | Sub-endosteal | Dry | 51 | 0.367 | 0.057 | 0.175 | 0.016 | 2.089 | 0.177 | 0.365 | 0.031 | 0.887 | 0.014 |
| GWS47 | Sub-endosteal | Dry | 41 | 0.091 | 0.030 | 0.081 | 0.018 | 1.100 | 0.225 | 0.458 | 0.096 | 0.950 | 0.084 |
| GWS48 | Sub-endosteal | Dry | 43 | 0.235 | 0.132 | 0.133 | 0.043 | 1.641 | 0.544 | 0.443 | 0.094 | 0.874 | 0.076 |
| GWS60 | Sub-endosteal | Dry | 40 | 0.049 | 0.008 | 0.055 | 0.009 | 0.901 | 0.099 | 0.551 | 0.080 | 1.072 | 0.072 |
| GWS69 | Sub-endosteal | Dry | 16 | 0.309 | 0.063 | 0.148 | 0.020 | 2.080 | 0.281 | 0.401 | 0.048 | 0.864 | 0.060 |
| GWS02 | Mid-cortical | Dry | 42 | 0.370 | 0.022 | 0.155 | 0.007 | 2.391 | 0.103 | 0.361 | 0.020 | 0.836 | 0.016 |
| GWS19 | Mid-cortical | Dry | 60 | 0.369 | 0.039 | 0.180 | 0.014 | 2.049 | 0.115 | 0.369 | 0.024 | 0.887 | 0.015 |
| GWS47 | Mid-cortical | Dry | 48 | 0.454 | 0.043 | 0.187 | 0.010 | 2.427 | 0.187 | 0.361 | 0.040 | 0.833 | 0.024 |
| GWS48 | Mid-cortical | Dry | 48 | 0.361 | 0.020 | 0.159 | 0.006 | 2.277 | 0.112 | 0.368 | 0.013 | 0.834 | 0.012 |
| GWS60 | Mid-cortical | Dry | 48 | 0.292 | 0.023 | 0.145 | 0.010 | 2.014 | 0.109 | 0.335 | 0.022 | 0.904 | 0.023 |
| GWS69 | Mid-cortical | Dry | 60 | 0.396 | 0.022 | 0.166 | 0.009 | 2.389 | 0.087 | 0.358 | 0.019 | 0.852 | 0.016 |
| GWS02 | Sub-periosteal | Dry | 46 | 0.188 | 0.079 | 0.121 | 0.026 | 1.502 | 0.404 | 0.484 | 0.066 | 0.953 | 0.072 |
| GWS19 | Sub-periosteal | Dry | 70 | 0.166 | 0.068 | 0.117 | 0.022 | 1.357 | 0.340 | 0.499 | 0.049 | 0.994 | 0.083 |
| GWS47 | Sub-periosteal | Dry | 48 | 0.190 | 0.095 | 0.142 | 0.076 | 1.346 | 0.262 | 0.517 | 0.079 | 0.932 | 0.065 |
| GWS48 | Sub-periosteal | Dry | 53 | 0.217 | 0.105 | 0.133 | 0.033 | 1.569 | 0.497 | 0.469 | 0.120 | 0.916 | 0.110 |
| GWS60 | Sub-periosteal | Dry | 55 | 0.222 | 0.038 | 0.126 | 0.017 | 1.759 | 0.115 | 0.408 | 0.033 | 0.939 | 0.030 |
| GWS69 | Sub-periosteal | Dry | 68 | 0.384 | 0.027 | 0.173 | 0.007 | 2.216 | 0.120 | 0.359 | 0.019 | 0.873 | 0.016 |
| 12.0.1_7 | Sub-endosteal | Fossil | 24 | 0.221 | 0.089 | 0.123 | 0.036 | 1.729 | 0.245 | 0.443 | 0.072 | 0.928 | 0.045 |
| 13.0.1_1 | Sub-endosteal | Fossil | 13 | 0.025 | 0.010 | 0.067 | 0.016 | 0.356 | 0.131 | 0.453 | 0.176 | 1.114 | 0.479 |
| 32_11 | Sub-endosteal | Fossil | 26 | 0.047 | 0.019 | 0.048 | 0.009 | 0.942 | 0.247 | 0.473 | 0.126 | 1.087 | 0.182 |
| 32_6 | Sub-endosteal | Fossil | 26 | 0.070 | 0.039 | 0.086 | 0.129 | 1.019 | 0.273 | 0.532 | 0.069 | 1.038 | 0.080 |
| 4.0.1_1 | Sub-endosteal | Fossil | 60 | 0.024 | 0.006 | 0.044 | 0.009 | 0.582 | 0.323 | 0.555 | 0.134 | 1.224 | 0.208 |
| 4.1.5_1 | Sub-endosteal | Fossil | 24 | 0.028 | 0.004 | 0.062 | 0.008 | 0.446 | 0.051 | 0.550 | 0.103 | 1.182 | 0.066 |
| 5.0.32_5 | Sub-endosteal | Fossil | 18 | 0.022 | 0.006 | 0.044 | 0.010 | 0.519 | 0.149 | 0.561 | 0.156 | 1.180 | 0.095 |
| 7.13.2_1 | Sub-endosteal | Fossil | 24 | 0.028 | 0.005 | 0.050 | 0.008 | 0.571 | 0.083 | 0.624 | 0.081 | 1.170 | 0.072 |
| 7.8.4_1 | Sub-endosteal | Fossil | 18 | 0.074 | 0.195 | 0.258 | 0.859 | 0.519 | 0.144 | 0.657 | 0.082 | 1.140 | 0.119 |
| 12.0.1_7 | Mid-cortical | Fossil | 24 | 0.305 | 0.011 | 0.147 | 0.005 | 2.081 | 0.079 | 0.364 | 0.010 | 0.873 | 0.013 |
| 13.0.1_1 | Mid-cortical | Fossil | 23 | 0.027 | 0.014 | 0.063 | 0.016 | 0.451 | 0.245 | 0.581 | 0.271 | 1.191 | 0.199 |
| 32_11 | Mid-cortical | Fossil | 30 | 0.026 | 0.003 | 0.041 | 0.003 | 0.634 | 0.068 | 0.627 | 0.072 | 1.139 | 0.079 |
| 32_6 | Mid-cortical | Fossil | 30 | 0.027 | 0.004 | 0.046 | 0.007 | 0.597 | 0.077 | 0.655 | 0.050 | 1.146 | 0.048 |
| 4.0.1_1 | Mid-cortical | Fossil | 66 | 0.033 | 0.009 | 0.045 | 0.005 | 0.723 | 0.147 | 0.578 | 0.057 | 1.122 | 0.069 |
| 4.1.5_1 | Mid-cortical | Fossil | 24 | 0.029 | 0.003 | 0.056 | 0.004 | 0.515 | 0.046 | 0.708 | 0.045 | 1.108 | 0.036 |
| 5.0.32_5 | Mid-cortical | Fossil | 24 | 0.022 | 0.002 | 0.038 | 0.004 | 0.596 | 0.076 | 0.673 | 0.038 | 1.120 | 0.034 |
| 7.13.2_1 | Mid-cortical | Fossil | 24 | 0.028 | 0.006 | 0.050 | 0.011 | 0.558 | 0.093 | 0.612 | 0.119 | 1.160 | 0.068 |
| 7.8.4_1 | Mid-cortical | Fossil | 24 | 0.023 | 0.003 | 0.040 | 0.005 | 0.572 | 0.057 | 0.615 | 0.090 | 1.129 | 0.081 |
| 12.0.1_7 | Sub-periosteal | Fossil | 27 | 0.183 | 0.072 | 0.112 | 0.034 | 1.591 | 0.215 | 0.502 | 0.069 | 0.944 | 0.036 |
| 13.0.1_1 | Sub-periosteal | Fossil | 23 | 0.015 | 0.011 | 0.033 | 0.017 | 0.446 | 0.364 | 0.542 | 0.329 | 1.279 | 0.446 |
| 32_11 | Sub-periosteal | Fossil | 31 | 0.053 | 0.041 | 0.053 | 0.034 | 0.959 | 0.329 | 0.669 | 0.562 | 1.069 | 0.270 |
| 32_6 | Sub-periosteal | Fossil | 33 | 0.044 | 0.014 | 0.050 | 0.006 | 0.886 | 0.229 | 0.458 | 0.079 | 1.145 | 0.541 |
| 4.0.1_1 | Sub-periosteal | Fossil | 73 | 0.056 | 0.017 | 0.055 | 0.005 | 1.004 | 0.236 | 0.502 | 0.066 | 1.057 | 0.078 |
| 4.1.5_1 | Sub-periosteal | Fossil | 28 | 0.042 | 0.016 | 0.055 | 0.007 | 0.770 | 0.259 | 0.628 | 0.075 | 1.065 | 0.102 |
| 5.0.32_5 | Sub-periosteal | Fossil | 28 | 0.028 | 0.003 | 0.045 | 0.005 | 0.623 | 0.078 | 0.646 | 0.038 | 1.114 | 0.028 |
| 7.13.2_1 | Sub-periosteal | Fossil | 28 | 0.046 | 0.012 | 0.050 | 0.007 | 0.932 | 0.193 | 0.554 | 0.081 | 1.082 | 0.064 |
| 7.8.4_1 | Sub-periosteal | Fossil | 28 | 0.047 | 0.009 | 0.045 | 0.005 | 1.032 | 0.162 | 0.485 | 0.051 | 1.054 | 0.062 |
| GH04 | Sub-endosteal | Wet | 74 | 0.255 | 0.147 | 0.137 | 0.051 | 1.731 | 0.459 | 0.551 | 0.121 | 0.846 | 0.086 |
| GH10 | Sub-endosteal | Wet | 31 | 0.396 | 0.051 | 0.187 | 0.009 | 2.123 | 0.243 | 0.350 | 0.026 | 0.861 | 0.024 |
| GH18 | Sub-endosteal | Wet | 22 | 0.152 | 0.056 | 0.110 | 0.037 | 1.397 | 0.275 | 0.426 | 0.086 | 0.956 | 0.085 |
| GH19 | Sub-endosteal | Wet | 44 | 0.378 | 0.075 | 0.173 | 0.032 | 2.188 | 0.133 | 0.312 | 0.059 | 0.862 | 0.036 |
| GH9 | Sub-endosteal | Wet | 53 | 0.427 | 0.064 | 0.182 | 0.023 | 2.342 | 0.140 | 0.363 | 0.033 | 0.849 | 0.040 |
| GWW4 | Sub-endosteal | Wet | 29 | 0.481 | 0.053 | 0.194 | 0.015 | 2.477 | 0.214 | 0.335 | 0.038 | 0.876 | 0.047 |
| GWW61 | Sub-endosteal | Wet | 48 | 0.363 | 0.025 | 0.177 | 0.008 | 2.050 | 0.194 | 0.315 | 0.027 | 0.883 | 0.022 |
| GWW76 | Sub-endosteal | Wet | 32 | 0.312 | 0.028 | 0.175 | 0.009 | 1.787 | 0.114 | 0.269 | 0.033 | 1.003 | 0.025 |
| GH04 | Mid-cortical | Wet | 90 | 0.464 | 0.025 | 0.207 | 0.008 | 2.240 | 0.090 | 0.369 | 0.030 | 0.845 | 0.021 |
| GH10 | Mid-cortical | Wet | 42 | 0.439 | 0.047 | 0.202 | 0.007 | 2.174 | 0.230 | 0.358 | 0.016 | 0.854 | 0.014 |
| GH18 | Mid-cortical | Wet | 30 | 0.308 | 0.011 | 0.142 | 0.003 | 2.179 | 0.082 | 0.377 | 0.013 | 0.838 | 0.012 |
| GH19 | Mid-cortical | Wet | 48 | 0.378 | 0.040 | 0.190 | 0.010 | 1.989 | 0.145 | 0.358 | 0.022 | 0.892 | 0.016 |
| GH9 | Mid-cortical | Wet | 65 | 0.497 | 0.046 | 0.210 | 0.014 | 2.364 | 0.110 | 0.360 | 0.018 | 0.852 | 0.018 |
| GWW4 | Mid-cortical | Wet | 34 | 0.411 | 0.024 | 0.170 | 0.011 | 2.419 | 0.160 | 0.353 | 0.036 | 0.872 | 0.030 |
| GWW61 | Mid-cortical | Wet | 48 | 0.371 | 0.015 | 0.172 | 0.005 | 2.163 | 0.088 | 0.346 | 0.011 | 0.855 | 0.012 |
| GWW76 | Mid-cortical | Wet | 36 | 0.353 | 0.018 | 0.174 | 0.006 | 2.027 | 0.112 | 0.333 | 0.016 | 0.926 | 0.017 |
| GH04 | Sub-periosteal | Wet | 91 | 0.248 | 0.134 | 0.146 | 0.053 | 1.594 | 0.370 | 0.437 | 0.066 | 0.925 | 0.080 |
| GH10 | Sub-periosteal | Wet | 43 | 0.261 | 0.098 | 0.134 | 0.029 | 1.867 | 0.380 | 0.412 | 0.037 | 0.890 | 0.059 |
| GH18 | Sub-periosteal | Wet | 35 | 0.233 | 0.086 | 0.179 | 0.124 | 1.478 | 0.506 | 0.420 | 0.078 | 0.918 | 0.091 |
| GH19 | Sub-periosteal | Wet | 52 | 0.320 | 0.031 | 0.172 | 0.012 | 1.860 | 0.102 | 0.372 | 0.035 | 0.890 | 0.018 |
| GH9 | Sub-periosteal | Wet | 59 | 0.358 | 0.122 | 0.183 | 0.040 | 1.925 | 0.440 | 0.304 | 0.109 | 0.951 | 0.146 |
| GWW4 | Sub-periosteal | Wet | 38 | 0.429 | 0.061 | 0.173 | 0.018 | 2.472 | 0.243 | 0.339 | 0.038 | 0.885 | 0.050 |
| GWW61 | Sub-periosteal | Wet | 54 | 0.322 | 0.070 | 0.161 | 0.022 | 1.987 | 0.296 | 0.369 | 0.054 | 0.872 | 0.044 |
| GWW76 | Sub-periosteal | Wet | 42 | 0.365 | 0.057 | 0.202 | 0.039 | 1.820 | 0.172 | 0.333 | 0.073 | 0.913 | 0.045 |

### 3.2 s-FTIRM data

Table 9 Mineral diagenetic index data per sample ROI collected by s-FTIRM methods

|  |  |  | **C/P** | | **HPO_4_/P** | | **CI1** | | **CI2** | |
| --- | --- | --- | --- | --- | --- | --- | --- | --- | --- | --- |
| **Sample ID** | **Dataset** | **Environment** | **x̄** | **σ** | **x̄** | **σ** | **x̄** | **σ** | **x̄** | **σ** |
| GWS_02_On1 | ROI | Dry | 0.268 | 0.066 | 0.231 | 0.079 | 0.937 | 0.064 | 1.341 | 0.099 |
| GWS_02_On2 | ROI | Dry | 0.275 | 0.063 | 0.223 | 0.100 | 0.935 | 0.070 | 1.366 | 0.106 |
| GWS_19_On1 | ROI | Dry | 0.257 | 0.057 | 0.244 | 0.102 | 0.952 | 0.064 | 1.329 | 0.082 |
| GWS_19_On2 | ROI | Dry | 0.266 | 0.063 | 0.229 | 0.099 | 0.955 | 0.065 | 1.347 | 0.083 |
| GWS_47_On1 | ROI | Dry | 0.217 | 0.064 | 0.222 | 0.099 | 0.938 | 0.065 | 1.377 | 0.165 |
| GWS_47_On2 | ROI | Dry | 0.226 | 0.054 | 0.215 | 0.096 | 0.933 | 0.066 | 1.360 | 0.093 |
| GWS_48_On1 | ROI | Dry | 0.287 | 0.069 | 0.247 | 0.112 | 0.955 | 0.069 | 1.430 | 2.515 |
| GWS_48_On2 | ROI | Dry | 0.234 | 0.060 | 0.143 | 0.125 | 0.887 | 0.098 | 1.565 | 0.747 |
| GWS_60_On3 | ROI | Dry | 0.301 | 0.080 | 0.287 | 0.125 | 0.969 | 0.068 | 1.311 | 0.172 |
| GWS_60_On4 | ROI | Dry | 0.299 | 0.077 | 0.280 | 0.116 | 0.962 | 0.067 | 1.309 | 0.087 |
| GWS_69_On4 | ROI | Dry | 0.258 | 0.053 | 0.218 | 0.086 | 0.924 | 0.066 | 1.349 | 0.076 |
| GH_10_On4 | ROI | Wet | 0.254 | 0.065 | 0.167 | 0.095 | 0.912 | 0.075 | 1.458 | 0.189 |
| GH_18_On3 | ROI | Wet | 0.258 | 0.066 | 0.233 | 0.105 | 0.950 | 0.069 | 1.349 | 0.103 |
| GH_18_On4 | ROI | Wet | 0.243 | 0.061 | 0.200 | 0.101 | 0.938 | 0.076 | 1.401 | 0.300 |
| GH_19_On1 | ROI | Wet | 0.254 | 0.050 | 0.227 | 0.090 | 0.949 | 0.058 | 1.352 | 0.083 |
| GH_19_On2 | ROI | Wet | 0.228 | 0.054 | 0.235 | 0.106 | 0.954 | 0.065 | 1.335 | 0.100 |
| GH_4_On1 | ROI | Wet | 0.308 | 0.070 | 0.235 | 0.108 | 0.951 | 0.073 | 1.362 | 0.111 |
| GH_4_On2 | ROI | Wet | 0.180 | 0.056 | 0.196 | 0.114 | 0.995 | 0.071 | 1.527 | 0.432 |
| GH_9_On1 | ROI | Wet | 0.274 | 0.057 | 0.264 | 0.101 | 0.956 | 0.057 | 1.319 | 0.077 |
| GH_9_On2 | ROI | Wet | 0.251 | 0.055 | 0.268 | 0.113 | 0.962 | 0.058 | 1.308 | 0.091 |
| GWW_4_On1 | ROI | Wet | 0.228 | 0.057 | 0.251 | 0.116 | 0.964 | 0.066 | 1.356 | 0.201 |
| GWW_4_On2 | ROI | Wet | 0.228 | 0.049 | 0.218 | 0.088 | 0.944 | 0.056 | 1.387 | 0.173 |
| GWW_61_On1 | ROI | Wet | 0.276 | 0.066 | 0.213 | 0.094 | 0.930 | 0.072 | 1.372 | 0.130 |
| GWW_76_On1 | ROI | Wet | 0.257 | 0.070 | 0.255 | 0.130 | 0.985 | 0.057 | 1.373 | 0.214 |
| GWW_76_On2 | ROI | Wet | 0.239 | 0.063 | 0.220 | 0.097 | 0.959 | 0.059 | 1.372 | 0.108 |
| 13.0.2_1_On3 | ROI | Fossil | 0.221 | 0.080 | 0.210 | 0.095 | 0.971 | 0.065 | 1.464 | 0.129 |
| 13.0.2_1_On4 | ROI | Fossil | 0.219 | 0.059 | 0.175 | 0.094 | 0.979 | 0.070 | 1.520 | 0.159 |
| 32_6_On3 | ROI | Fossil | 0.208 | 0.060 | 0.153 | 0.063 | 0.992 | 0.058 | 1.526 | 0.115 |
| 32_6_On4 | ROI | Fossil | 0.183 | 0.060 | 0.204 | 0.095 | 0.990 | 0.066 | 1.452 | 0.112 |
| 4.0.1_1_On1.2 | ROI | Fossil | 0.247 | 0.090 | 0.176 | 0.105 | 0.983 | 0.073 | 1.520 | 0.209 |
| 4.0.1_1_On2 | ROI | Fossil | 0.211 | 0.054 | 0.153 | 0.074 | 0.982 | 0.070 | 1.531 | 0.129 |
| 4.1.5_1_On3 | ROI | Fossil | 0.196 | 0.049 | 0.214 | 0.106 | 0.973 | 0.082 | 1.453 | 0.128 |
| 4.1.5_1_On4 | ROI | Fossil | 0.232 | 0.064 | 0.208 | 0.104 | 0.994 | 0.066 | 1.472 | 0.135 |
| 5.0.32_5_On3 | ROI | Fossil | 0.201 | 0.072 | 0.209 | 0.117 | 1.007 | 0.081 | 1.495 | 0.161 |
| 5.0.32_5_On4 | ROI | Fossil | 0.287 | 0.083 | 0.119 | 0.096 | 0.998 | 0.080 | 1.772 | 0.377 |
| 7.13.2_1_On3 | ROI | Fossil | 0.219 | 0.058 | 0.180 | 0.091 | 1.018 | 0.044 | 1.526 | 0.156 |
| 7.13.2_1_On4 | ROI | Fossil | 0.240 | 0.077 | 0.182 | 0.102 | 0.970 | 0.088 | 1.496 | 0.155 |
| 7.8.4_1_On1 | ROI | Fossil | 0.197 | 0.035 | 0.066 | 0.046 | 0.949 | 0.073 | 1.971 | 0.444 |
| 7.8.4_1_On2 | ROI | Fossil | 0.241 | 0.078 | 0.201 | 0.095 | 1.015 | 0.058 | 1.472 | 0.139 |

Table 10 Amide region diagenetic index data per sample ROI collected by -FTIRM methods

|  |  |  | **AmI/P** | | **AmII/P** | | **AmI/AmII** | | **Collagen Integrity** | | **Random Coils**  **(α)** | |
| --- | --- | --- | --- | --- | --- | --- | --- | --- | --- | --- | --- | --- |
| **Sample Name** | **Dataset** | **Environment** | **x̄** | **σ** | **x̄** | **σ** | **x̄** | **σ** | **x̄** | **σ** | **x̄** | **σ** |
| GWS_02_On1 | ROI | Dry | 0.287 | 0.057 | 0.226 | 0.063 | 1.589 | 1.355 | 0.309 | 0.040 | 0.896 | 0.051 |
| GWS_02_On2 | ROI | Dry | 0.283 | 0.062 | 0.231 | 0.075 | 1.481 | 0.194 | 0.297 | 0.050 | 0.922 | 0.051 |
| GWS_19_On1 | ROI | Dry | 0.256 | 0.055 | 0.201 | 0.052 | 1.469 | 0.163 | 0.293 | 0.054 | 0.969 | 0.057 |
| GWS_19_On2 | ROI | Dry | 0.272 | 0.053 | 0.217 | 0.066 | 1.464 | 0.212 | 0.241 | 0.050 | 0.996 | 0.062 |
| GWS_47_On1 | ROI | Dry | 0.249 | 0.077 | 0.203 | 0.085 | 1.491 | 0.356 | 0.284 | 0.076 | 0.963 | 0.077 |
| GWS_47_On2 | ROI | Dry | 0.276 | 0.054 | 0.217 | 0.076 | 1.565 | 0.344 | 0.264 | 0.055 | 0.926 | 0.064 |
| GWS_48_On1 | ROI | Dry | 0.313 | 0.064 | 0.244 | 0.068 | 1.456 | 0.151 | 0.276 | 0.054 | 0.958 | 0.056 |
| GWS_48_On2 | ROI | Dry | 0.210 | 0.067 | 0.192 | 0.058 | 1.417 | 0.201 | 0.209 | 0.080 | 1.008 | 0.084 |
| GWS_60_On3 | ROI | Dry | 0.351 | 0.084 | 0.278 | 0.066 | 1.414 | 0.154 | 0.295 | 0.042 | 0.937 | 0.050 |
| GWS_60_On4 | ROI | Dry | 0.360 | 0.078 | 0.273 | 0.062 | 1.489 | 0.154 | 0.300 | 0.040 | 0.947 | 0.050 |
| GWS_69_On4 | ROI | Dry | 0.246 | 0.048 | 0.207 | 0.058 | 1.453 | 0.217 | 0.279 | 0.052 | 0.922 | 0.047 |
| GH_10_On4 | ROI | Wet | 0.223 | 0.051 | 0.222 | 0.075 | 1.291 | 0.197 | 0.282 | 0.055 | 0.967 | 0.060 |
| GH_18_On3 | ROI | Wet | 0.261 | 0.064 | 0.216 | 0.063 | 1.420 | 0.428 | 0.290 | 0.046 | 0.900 | 0.051 |
| GH_18_On4 | ROI | Wet | 0.217 | 0.053 | 0.189 | 0.063 | 1.409 | 0.901 | 0.253 | 0.057 | 0.929 | 0.063 |
| GH_19_On1 | ROI | Wet | 0.290 | 0.047 | 0.245 | 0.059 | 1.370 | 0.139 | 0.311 | 0.048 | 0.961 | 0.053 |
| GH_19_On2 | ROI | Wet | 0.254 | 0.048 | 0.207 | 0.055 | 1.429 | 0.207 | 0.321 | 0.053 | 0.954 | 0.054 |
| GH_4_On1 | ROI | Wet | 0.326 | 0.072 | 0.302 | 0.092 | 1.287 | 0.331 | 0.261 | 0.043 | 0.948 | 0.048 |
| GH_4_On2 | ROI | Wet | 0.166 | 0.108 | 0.142 | 0.082 | 1.332 | 0.617 | 0.180 | 0.080 | 1.032 | 0.164 |
| GH_9_On1 | ROI | Wet | 0.303 | 0.060 | 0.243 | 0.055 | 1.432 | 0.194 | 0.315 | 0.045 | 0.938 | 0.048 |
| GH_9_On2 | ROI | Wet | 0.294 | 0.067 | 0.229 | 0.066 | 1.535 | 0.457 | 0.299 | 0.048 | 0.956 | 0.051 |
| GWW_4_On1 | ROI | Wet | 0.356 | 0.081 | 0.254 | 0.064 | 1.565 | 0.186 | 0.321 | 0.044 | 0.961 | 0.051 |
| GWW_4_On2 | ROI | Wet | 0.293 | 0.062 | 0.224 | 0.095 | 1.712 | 1.541 | 0.235 | 0.051 | 0.984 | 0.045 |
| GWW_61_On1 | ROI | Wet | 0.288 | 0.057 | 0.257 | 0.060 | 1.330 | 0.179 | 0.308 | 0.046 | 0.901 | 0.047 |
| GWW_76_On1 | ROI | Wet | 0.409 | 0.112 | 0.309 | 0.101 | 1.449 | 0.643 | 0.264 | 0.049 | 0.990 | 0.053 |
| GWW_76_On2 | ROI | Wet | 0.312 | 0.067 | 0.251 | 0.083 | 1.462 | 0.341 | 0.333 | 0.039 | 0.940 | 0.048 |
| 13.0.2_1_On3 | ROI | Fossil | 0.018 | 0.010 | 0.066 | 0.033 | 0.486 | 0.132 | 0.142 | 0.099 | 4.013 | 15.108 |
| 13.0.2_1_On4 | ROI | Fossil | 0.018 | 0.011 | 0.046 | 0.031 | 0.454 | 0.176 | 0.781 | 0.914 | 5.424 | 50.823 |
| 32_6_On3 | ROI | Fossil | 0.022 | 0.010 | 0.046 | 0.029 | 0.690 | 0.486 | 0.162 | 0.135 | 2.127 | 2.661 |
| 32_6_On4 | ROI | Fossil | 0.036 | 0.052 | 0.054 | 0.040 | 1.243 | 0.536 | 0.198 | 0.114 | 1.241 | 0.987 |
| 4.0.1_1_On1.2 | ROI | Fossil | 0.064 | 0.025 | 0.073 | 0.033 | 1.192 | 1.556 | 0.286 | 0.088 | 0.952 | 0.199 |
| 4.0.1_1_On2 | ROI | Fossil | 0.028 | 0.011 | 0.053 | 0.029 | 0.739 | 0.385 | 0.148 | 0.135 | 1.360 | 0.783 |
| 4.1.5_1_On3 | ROI | Fossil | 0.018 | 0.012 | 0.050 | 0.048 | 0.850 | 2.085 | 1.805 | 3.805 | 6.131 | 11.833 |
| 4.1.5_1_On4 | ROI | Fossil | 0.015 | 0.010 | 0.051 | 0.032 | 0.490 | 0.177 | - | - | 8.203 | 14.945 |
| 5.0.32_5_On3 | ROI | Fossil | 0.016 | 0.011 | 0.045 | 0.030 | 0.507 | 0.179 | 0.270 | 0.178 | 8.212 | 37.214 |
| 5.0.32_5_On4 | ROI | Fossil | 0.030 | 0.015 | 0.085 | 0.043 | 0.532 | 0.129 | 0.278 | 0.291 | 2.406 | 2.406 |
| 7.13.2_1_On3 | ROI | Fossil | 0.027 | 0.011 | 0.050 | 0.023 | 0.702 | 0.276 | 0.135 | 0.098 | 2.645 | 4.301 |
| 7.13.2_1_On4 | ROI | Fossil | 0.037 | 0.014 | 0.063 | 0.025 | 0.734 | 0.347 | 0.161 | 0.102 | 1.497 | 3.558 |
| 7.8.4_1_On1 | ROI | Fossil | 0.024 | 0.020 | 0.061 | 0.039 | 0.317 | 0.180 | 0.264 | 0.150 | 8.248 | 28.338 |
| 7.8.4_1_On2 | ROI | Fossil | 0.023 | 0.011 | 0.046 | 0.023 | 0.729 | 0.771 | 0.155 | 0.138 | 4.060 | 35.923 |

Table 11 Mineral diagenetic index data per sample osteon area (On.Ar) and interstitial bone area (In.Ar) collected by s-FTIRM methods

|  |  |  | **C/P** | | | **HPO_4_/P** | | **CI1** | | **CI2** | |
| --- | --- | --- | --- | --- | --- | --- | --- | --- | --- | --- | --- |
| **Sample ID** | **Dataset** | **Environment** | **x̄** | **σ** | **x̄** | | **σ** | **x̄** | **σ** | **x̄** | **σ** |
| GWS_02_On1 | On.Ar | Dry | 0.269 | 0.039 | 0.183 | | 0.054 | 0.899 | 0.054 | 1.387 | 0.138 |
| GWS_02_On1 | In.Ar | Dry | 0.276 | 0.065 | 0.287 | | 0.074 | 0.978 | 0.053 | 1.295 | 0.047 |
| GWS_02_On2 | On.Ar | Dry | 0.275 | 0.059 | 0.206 | | 0.099 | 0.922 | 0.070 | 1.383 | 0.108 |
| GWS_02_On2 | In.Ar | Dry | 0.319 | 0.046 | 0.312 | | 0.060 | 1.004 | 0.029 | 1.281 | 0.045 |
| GWS_19_On1 | On.Ar | Dry | 0.261 | 0.045 | 0.235 | | 0.100 | 0.949 | 0.065 | 1.344 | 0.084 |
| GWS_19_On1 | In.Ar | Dry | 0.262 | 0.078 | 0.269 | | 0.113 | 0.962 | 0.062 | 1.302 | 0.080 |
| GWS_19_On2 | On.Ar | Dry | 0.250 | 0.032 | 0.187 | | 0.092 | 0.917 | 0.061 | 1.378 | 0.081 |
| GWS_19_On2 | In.Ar | Dry | 0.284 | 0.079 | 0.267 | | 0.089 | 0.990 | 0.047 | 1.320 | 0.074 |
| GWS_47_On1 | On.Ar | Dry | 0.225 | 0.073 | 0.256 | | 0.082 | 0.955 | 0.056 | 1.314 | 0.073 |
| GWS_47_On1 | In.Ar | Dry | 0.217 | 0.046 | 0.167 | | 0.101 | 0.911 | 0.071 | 1.479 | 0.219 |
| GWS_47_On2 | On.Ar | Dry | 0.238 | 0.040 | 0.201 | | 0.092 | 0.930 | 0.067 | 1.379 | 0.089 |
| GWS_47_On2 | In.Ar | Dry | 0.165 | 0.081 | 0.292 | | 0.094 | 0.941 | 0.062 | 1.264 | 0.049 |
| GWS_48_On1 | On.Ar | Dry | 0.274 | 0.041 | 0.156 | | 0.047 | 0.910 | 0.056 | 1.753 | 4.516 |
| GWS_48_On1 | In.Ar | Dry | 0.298 | 0.078 | 0.291 | | 0.109 | 0.976 | 0.064 | 1.283 | 0.077 |
| GWS_48_On2 | On.Ar | Dry | 0.252 | 0.075 | 0.188 | | 0.146 | 0.916 | 0.095 | 1.453 | 0.231 |
| GWS_48_On2 | In.Ar | Dry | 0.217 | 0.038 | 0.101 | | 0.089 | 0.858 | 0.096 | 1.694 | 1.174 |
| GWS_60_On3 | On.Ar | Dry | 0.327 | 0.092 | 0.321 | | 0.113 | 0.982 | 0.061 | 1.279 | 0.071 |
| GWS_60_On3 | In.Ar | Dry | 0.275 | 0.043 | 0.237 | | 0.128 | 0.949 | 0.077 | 1.359 | 0.262 |
| GWS_60_On4 | On.Ar | Dry | 0.338 | 0.090 | 0.323 | | 0.108 | 0.984 | 0.057 | 1.281 | 0.067 |
| GWS_60_On4 | In.Ar | Dry | 0.268 | 0.046 | 0.235 | | 0.109 | 0.938 | 0.069 | 1.340 | 0.092 |
| GWS_69_On4 | On.Ar | Dry | 0.288 | 0.058 | 0.254 | | 0.076 | 0.952 | 0.050 | 1.317 | 0.060 |
| GWS_69_On4 | In.Ar | Dry | 0.246 | 0.045 | 0.200 | | 0.084 | 0.910 | 0.068 | 1.365 | 0.078 |
| GH_10_On4 | On.Ar | Wet | 0.295 | 0.063 | 0.206 | | 0.089 | 0.950 | 0.056 | 1.393 | 0.171 |
| GH_10_On4 | In.Ar | Wet | 0.214 | 0.035 | 0.114 | | 0.077 | 0.860 | 0.072 | 1.551 | 0.187 |
| GH_18_On3 | On.Ar | Wet | 0.271 | 0.067 | 0.249 | | 0.088 | 0.967 | 0.057 | 1.331 | 0.073 |
| GH_18_On3 | In.Ar | Wet | 0.254 | 0.064 | 0.227 | | 0.126 | 0.938 | 0.081 | 1.361 | 0.134 |
| GH_18_On4 | On.Ar | Wet | 0.277 | 0.070 | 0.268 | | 0.085 | 0.993 | 0.038 | 1.320 | 0.071 |
| GH_18_On4 | In.Ar | Wet | 0.228 | 0.050 | 0.160 | | 0.088 | 0.908 | 0.076 | 1.448 | 0.365 |
| GH_19_On1 | On.Ar | Wet | 0.257 | 0.058 | 0.252 | | 0.081 | 0.965 | 0.050 | 1.322 | 0.065 |
| GH_19_On1 | In.Ar | Wet | 0.257 | 0.036 | 0.191 | | 0.091 | 0.924 | 0.062 | 1.399 | 0.086 |
| GH_19_On2 | On.Ar | Wet | 0.210 | 0.048 | 0.222 | | 0.106 | 0.943 | 0.068 | 1.347 | 0.114 |
| GH_19_On2 | In.Ar | Wet | 0.245 | 0.055 | 0.253 | | 0.105 | 0.967 | 0.061 | 1.318 | 0.082 |
| GH_4_On1 | On.Ar | Wet | 0.366 | 0.069 | 0.298 | | 0.087 | 0.992 | 0.047 | 1.302 | 0.067 |
| GH_4_On1 | In.Ar | Wet | 0.283 | 0.053 | 0.199 | | 0.107 | 0.923 | 0.078 | 1.398 | 0.116 |
| GH_4_On2 | On.Ar | Wet | 0.226 | 0.053 | 0.261 | | 0.107 | 1.029 | 0.036 | 1.382 | 0.114 |
| GH_4_On2 | In.Ar | Wet | 0.144 | 0.026 | 0.141 | | 0.090 | 0.972 | 0.083 | 1.637 | 0.536 |
| GH_9_On1 | On.Ar | Wet | 0.274 | 0.051 | 0.294 | | 0.110 | 0.973 | 0.053 | 1.297 | 0.074 |
| GH_9_On1 | In.Ar | Wet | 0.280 | 0.070 | 0.231 | | 0.073 | 0.936 | 0.052 | 1.341 | 0.069 |
| GH_9_On2 | On.Ar | Wet | 0.258 | 0.067 | 0.323 | | 0.091 | 0.985 | 0.039 | 1.259 | 0.066 |
| GH_9_On2 | In.Ar | Wet | 0.248 | 0.046 | 0.211 | | 0.099 | 0.937 | 0.064 | 1.358 | 0.086 |
| GWW_4_On1 | On.Ar | Wet | 0.236 | 0.063 | 0.288 | | 0.106 | 0.978 | 0.051 | 1.302 | 0.077 |
| GWW_4_On1 | In.Ar | Wet | 0.214 | 0.042 | 0.200 | | 0.116 | 0.941 | 0.082 | 1.442 | 0.306 |
| GWW_4_On2 | On.Ar | Wet | 0.245 | 0.037 | 0.181 | | 0.059 | 0.922 | 0.047 | 1.425 | 0.186 |
| GWW_4_On2 | In.Ar | Wet | 0.172 | 0.018 | 0.384 | | 0.049 | 1.017 | 0.021 | 1.245 | 0.037 |
| GWW_61_On1 | On.Ar | Wet | 0.280 | 0.057 | 0.182 | | 0.083 | 0.914 | 0.074 | 1.409 | 0.148 |
| GWW_61_On1 | In.Ar | Wet | 0.271 | 0.082 | 0.244 | | 0.094 | 0.945 | 0.067 | 1.330 | 0.081 |
| GWW_76_On1 | On.Ar | Wet | 0.287 | 0.066 | 0.306 | | 0.109 | 1.010 | 0.041 | 1.298 | 0.099 |
| GWW_76_On1 | In.Ar | Wet | 0.233 | 0.053 | 0.124 | | 0.071 | 0.929 | 0.051 | 1.560 | 0.280 |
| GWW_76_On2 | On.Ar | Wet | 0.254 | 0.079 | 0.259 | | 0.100 | 0.971 | 0.058 | 1.326 | 0.091 |
| GWW_76_On2 | In.Ar | Wet | 0.232 | 0.051 | 0.193 | | 0.090 | 0.949 | 0.061 | 1.406 | 0.111 |
| 13.0.2_1_On3 | On.Ar | Fossil | 0.251 | 0.094 | 0.238 | | 0.092 | 0.973 | 0.072 | 1.412 | 0.096 |
| 13.0.2_1_On3 | In.Ar | Fossil | 0.179 | 0.041 | 0.174 | | 0.080 | 0.971 | 0.051 | 1.526 | 0.134 |
| 13.0.2_1_On4 | On.Ar | Fossil | 0.153 | 0.024 | 0.185 | | 0.061 | 1.044 | 0.048 | 1.541 | 0.129 |
| 13.0.2_1_On4 | In.Ar | Fossil | 0.227 | 0.057 | 0.167 | | 0.098 | 0.966 | 0.069 | 1.528 | 0.165 |
| 32_6_On3 | On.Ar | Fossil | 0.158 | 0.031 | 0.118 | | 0.050 | 0.989 | 0.076 | 1.615 | 0.104 |
| 32_6_On3 | In.Ar | Fossil | 0.230 | 0.058 | 0.166 | | 0.062 | 0.991 | 0.049 | 1.491 | 0.101 |
| 32_6_On4 | On.Ar | Fossil | 0.113 | 0.065 | 0.327 | | 0.123 | 1.021 | 0.048 | 1.335 | 0.145 |
| 32_6_On4 | In.Ar | Fossil | 0.193 | 0.056 | 0.187 | | 0.082 | 0.983 | 0.067 | 1.469 | 0.102 |
| 4.0.1_1_On1.2 | On.Ar | Fossil | 0.190 | 0.038 | 0.143 | | 0.111 | 0.968 | 0.074 | 1.583 | 0.196 |
| 4.0.1_1_On1.2 | In.Ar | Fossil | 0.280 | 0.099 | 0.181 | | 0.095 | 0.984 | 0.076 | 1.503 | 0.225 |
| 4.0.1_1_On2 | On.Ar | Fossil | 0.215 | 0.037 | 0.116 | | 0.065 | 0.967 | 0.070 | 1.617 | 0.132 |
| 4.0.1_1_On2 | In.Ar | Fossil | 0.213 | 0.064 | 0.171 | | 0.075 | 0.985 | 0.074 | 1.492 | 0.113 |
| 4.1.5_1_On3 | On.Ar | Fossil | 0.189 | 0.038 | 0.134 | | 0.041 | 0.908 | 0.070 | 1.516 | 0.099 |
| 4.1.5_1_On3 | In.Ar | Fossil | 0.210 | 0.060 | 0.319 | | 0.082 | 1.032 | 0.040 | 1.356 | 0.092 |
| 4.1.5_1_On4 | On.Ar | Fossil | 0.229 | 0.046 | 0.154 | | 0.081 | 0.976 | 0.070 | 1.545 | 0.136 |
| 4.1.5_1_On4 | In.Ar | Fossil | 0.228 | 0.073 | 0.233 | | 0.104 | 1.000 | 0.060 | 1.436 | 0.119 |
| 5.0.32_5_On3 | On.Ar | Fossil | 0.200 | 0.086 | 0.287 | | 0.125 | 1.018 | 0.058 | 1.401 | 0.150 |
| 5.0.32_5_On3 | In.Ar | Fossil | 0.204 | 0.066 | 0.175 | | 0.104 | 1.003 | 0.088 | 1.540 | 0.156 |
| 7.13.2_1_On3 | On.Ar | Fossil | 0.206 | 0.045 | 0.183 | | 0.103 | 1.018 | 0.045 | 1.537 | 0.184 |
| 7.13.2_1_On3 | In.Ar | Fossil | 0.236 | 0.066 | 0.178 | | 0.082 | 1.019 | 0.044 | 1.516 | 0.130 |
| 7.13.2_1_On4 | On.Ar | Fossil | 0.267 | 0.066 | 0.209 | | 0.086 | 1.009 | 0.065 | 1.471 | 0.169 |
| 7.13.2_1_On4 | In.Ar | Fossil | 0.210 | 0.075 | 0.157 | | 0.109 | 0.930 | 0.092 | 1.520 | 0.156 |
| 7.8.4_1_On1 | On.Ar | Fossil | 0.198 | 0.035 | 0.070 | | 0.051 | 0.959 | 0.067 | 1.969 | 0.480 |
| 7.8.4_1_On1 | In.Ar | Fossil | 0.187 | 0.022 | 0.060 | | 0.027 | 0.945 | 0.088 | 1.862 | 0.221 |
| 7.8.4_1_On2 | On.Ar | Fossil | 0.150 | 0.032 | 0.108 | | 0.053 | 0.992 | 0.070 | 1.638 | 0.166 |
| 7.8.4_1_On2 | In.Ar | Fossil | 0.264 | 0.070 | 0.216 | | 0.094 | 1.023 | 0.049 | 1.454 | 0.130 |

Table 12 Amide region diagenetic index data per sample osteon area (On.Ar) and interstitial bone area (In.Ar) collected by s-FTIRM methods

|  |  |  | **AmI/P** | | **AmII/P** | | **AmI/AmII** | | **Collagen Integrity** | | | **Random Coils**  **(α)** | |
| --- | --- | --- | --- | --- | --- | --- | --- | --- | --- | --- | --- | --- | --- |
| **Sample ID** | **Dataset** | **Environment** | **x̄** | **σ** | **x̄** | **σ** | **x̄** | **σ** | | **x̄** | **σ** | **x̄** | **σ** |
| GWS_02_On1 | On.Ar | Dry | 0.302 | 0.042 | 0.264 | 0.070 | 1.504 | 0.174 | | 0.301 | 0.040 | 0.923 | 0.047 |
| GWS_02_On1 | In.Ar | Dry | 0.297 | 0.063 | 0.198 | 0.031 | 1.560 | 0.128 | | 0.324 | 0.026 | 0.886 | 0.029 |
| GWS_02_On2 | On.Ar | Dry | 0.286 | 0.055 | 0.241 | 0.077 | 1.472 | 0.187 | | 0.298 | 0.050 | 0.928 | 0.049 |
| GWS_02_On2 | In.Ar | Dry | 0.325 | 0.048 | 0.219 | 0.036 | 1.538 | 0.145 | | 0.291 | 0.030 | 0.905 | 0.021 |
| GWS_19_On1 | On.Ar | Dry | 0.261 | 0.051 | 0.202 | 0.057 | 1.498 | 0.152 | | 0.289 | 0.045 | 0.988 | 0.042 |
| GWS_19_On1 | In.Ar | Dry | 0.248 | 0.068 | 0.201 | 0.051 | 1.400 | 0.193 | | 0.296 | 0.068 | 0.936 | 0.063 |
| GWS_19_On2 | On.Ar | Dry | 0.272 | 0.031 | 0.219 | 0.075 | 1.563 | 0.176 | | 0.238 | 0.044 | 1.008 | 0.054 |
| GWS_19_On2 | In.Ar | Dry | 0.272 | 0.066 | 0.218 | 0.059 | 1.365 | 0.207 | | 0.245 | 0.053 | 0.984 | 0.066 |
| GWS_47_On1 | On.Ar | Dry | 0.264 | 0.086 | 0.202 | 0.061 | 1.475 | 0.264 | | 0.285 | 0.082 | 0.960 | 0.079 |
| GWS_47_On1 | In.Ar | Dry | 0.233 | 0.061 | 0.211 | 0.114 | 1.486 | 0.464 | | 0.286 | 0.068 | 0.975 | 0.077 |
| GWS_47_On2 | On.Ar | Dry | 0.285 | 0.047 | 0.227 | 0.074 | 1.514 | 0.212 | | 0.254 | 0.041 | 0.944 | 0.042 |
| GWS_47_On2 | In.Ar | Dry | 0.226 | 0.068 | 0.167 | 0.069 | 1.857 | 0.702 | | 0.333 | 0.079 | 0.819 | 0.072 |
| GWS_48_On1 | On.Ar | Dry | 0.292 | 0.032 | 0.243 | 0.089 | 1.509 | 0.148 | | 0.265 | 0.041 | 0.984 | 0.043 |
| GWS_48_On1 | In.Ar | Dry | 0.327 | 0.072 | 0.250 | 0.054 | 1.418 | 0.138 | | 0.282 | 0.060 | 0.944 | 0.057 |
| GWS_48_On2 | On.Ar | Dry | 0.235 | 0.076 | 0.207 | 0.059 | 1.402 | 0.212 | | 0.233 | 0.094 | 0.980 | 0.095 |
| GWS_48_On2 | In.Ar | Dry | 0.187 | 0.051 | 0.174 | 0.047 | 1.448 | 0.169 | | 0.182 | 0.049 | 1.039 | 0.047 |
| GWS_60_On3 | On.Ar | Dry | 0.383 | 0.093 | 0.303 | 0.058 | 1.378 | 0.161 | | 0.304 | 0.045 | 0.922 | 0.049 |
| GWS_60_On3 | In.Ar | Dry | 0.319 | 0.048 | 0.254 | 0.065 | 1.463 | 0.129 | | 0.283 | 0.032 | 0.963 | 0.041 |
| GWS_60_On4 | On.Ar | Dry | 0.400 | 0.087 | 0.304 | 0.056 | 1.426 | 0.128 | | 0.312 | 0.043 | 0.938 | 0.051 |
| GWS_60_On4 | In.Ar | Dry | 0.330 | 0.055 | 0.250 | 0.058 | 1.557 | 0.153 | | 0.290 | 0.034 | 0.957 | 0.049 |
| GWS_69_On4 | On.Ar | Dry | 0.275 | 0.053 | 0.223 | 0.037 | 1.384 | 0.139 | | 0.269 | 0.047 | 0.929 | 0.044 |
| GWS_69_On4 | In.Ar | Dry | 0.235 | 0.041 | 0.202 | 0.065 | 1.473 | 0.211 | | 0.282 | 0.054 | 0.920 | 0.046 |
| GH_10_On4 | On.Ar | Wet | 0.253 | 0.048 | 0.224 | 0.056 | 1.295 | 0.133 | | 0.278 | 0.045 | 0.980 | 0.057 |
| GH_10_On4 | In.Ar | Wet | 0.192 | 0.032 | 0.233 | 0.095 | 1.268 | 0.213 | | 0.290 | 0.061 | 0.950 | 0.058 |
| GH_18_On3 | On.Ar | Wet | 0.279 | 0.065 | 0.220 | 0.050 | 1.400 | 0.119 | | 0.291 | 0.041 | 0.904 | 0.045 |
| GH_18_On3 | In.Ar | Wet | 0.249 | 0.057 | 0.213 | 0.079 | 1.481 | 0.689 | | 0.289 | 0.047 | 0.895 | 0.055 |
| GH_18_On4 | On.Ar | Wet | 0.238 | 0.066 | 0.198 | 0.055 | 1.269 | 0.125 | | 0.237 | 0.047 | 0.926 | 0.055 |
| GH_18_On4 | In.Ar | Wet | 0.206 | 0.041 | 0.188 | 0.069 | 1.470 | 1.127 | | 0.262 | 0.058 | 0.928 | 0.066 |
| GH_19_On1 | On.Ar | Wet | 0.291 | 0.053 | 0.247 | 0.047 | 1.318 | 0.132 | | 0.309 | 0.053 | 0.958 | 0.061 |
| GH_19_On1 | In.Ar | Wet | 0.292 | 0.040 | 0.250 | 0.075 | 1.433 | 0.128 | | 0.311 | 0.040 | 0.971 | 0.038 |
| GH_19_On2 | On.Ar | Wet | 0.246 | 0.048 | 0.200 | 0.070 | 1.491 | 0.258 | | 0.319 | 0.059 | 0.954 | 0.062 |
| GH_19_On2 | In.Ar | Wet | 0.260 | 0.046 | 0.210 | 0.030 | 1.374 | 0.109 | | 0.325 | 0.048 | 0.953 | 0.045 |
| GH_4_On1 | On.Ar | Wet | 0.381 | 0.074 | 0.331 | 0.060 | 1.236 | 0.068 | | 0.258 | 0.033 | 0.949 | 0.037 |
| GH_4_On1 | In.Ar | Wet | 0.303 | 0.054 | 0.301 | 0.108 | 1.317 | 0.443 | | 0.266 | 0.044 | 0.944 | 0.048 |
| GH_4_On2 | On.Ar | Wet | 0.262 | 0.081 | 0.210 | 0.057 | 1.306 | 0.231 | | 0.200 | 0.046 | 1.001 | 0.055 |
| GH_4_On2 | In.Ar | Wet | 0.087 | 0.057 | 0.084 | 0.055 | 1.374 | 0.837 | | 0.166 | 0.098 | 1.049 | 0.217 |
| GH_9_On1 | On.Ar | Wet | 0.317 | 0.057 | 0.229 | 0.036 | 1.486 | 0.111 | | 0.310 | 0.036 | 0.951 | 0.038 |
| GH_9_On1 | In.Ar | Wet | 0.292 | 0.062 | 0.264 | 0.070 | 1.382 | 0.277 | | 0.328 | 0.056 | 0.918 | 0.058 |
| GH_9_On2 | On.Ar | Wet | 0.341 | 0.067 | 0.249 | 0.054 | 1.542 | 0.181 | | 0.305 | 0.051 | 0.966 | 0.056 |
| GH_9_On2 | In.Ar | Wet | 0.261 | 0.049 | 0.220 | 0.074 | 1.535 | 0.619 | | 0.296 | 0.048 | 0.949 | 0.046 |
| GWW_4_On1 | On.Ar | Wet | 0.387 | 0.084 | 0.260 | 0.046 | 1.609 | 0.185 | | 0.320 | 0.045 | 0.964 | 0.054 |
| GWW_4_On1 | In.Ar | Wet | 0.314 | 0.057 | 0.250 | 0.090 | 1.496 | 0.186 | | 0.321 | 0.043 | 0.960 | 0.046 |
| GWW_4_On2 | On.Ar | Wet | 0.317 | 0.043 | 0.255 | 0.086 | 1.537 | 0.161 | | 0.236 | 0.056 | 0.990 | 0.047 |
| GWW_4_On2 | In.Ar | Wet | 0.181 | 0.026 | 0.115 | 0.014 | 1.508 | 0.157 | | 0.204 | 0.031 | 0.968 | 0.033 |
| GWW_61_On1 | On.Ar | Wet | 0.293 | 0.046 | 0.269 | 0.058 | 1.329 | 0.108 | | 0.306 | 0.041 | 0.916 | 0.040 |
| GWW_61_On1 | In.Ar | Wet | 0.282 | 0.071 | 0.250 | 0.060 | 1.316 | 0.265 | | 0.313 | 0.056 | 0.874 | 0.049 |
| GWW_76_On1 | On.Ar | Wet | 0.468 | 0.100 | 0.339 | 0.080 | 1.428 | 0.126 | | 0.275 | 0.045 | 0.985 | 0.047 |
| GWW_76_On1 | In.Ar | Wet | 0.336 | 0.076 | 0.299 | 0.121 | 1.338 | 0.207 | | 0.235 | 0.044 | 1.006 | 0.047 |
| GWW_76_On2 | On.Ar | Wet | 0.334 | 0.083 | 0.261 | 0.075 | 1.481 | 0.486 | | 0.338 | 0.047 | 0.933 | 0.057 |
| GWW_76_On2 | In.Ar | Wet | 0.302 | 0.054 | 0.251 | 0.091 | 1.455 | 0.243 | | 0.330 | 0.033 | 0.946 | 0.041 |
| 13.0.2_1_On3 | On.Ar | Fossil | 0.019 | 0.010 | 0.080 | 0.033 | 0.491 | 0.110 | | 0.144 | 0.173 | 4.447 | 12.516 |
| 13.0.2_1_On3 | In.Ar | Fossil | 0.016 | 0.009 | 0.047 | 0.022 | 0.458 | 0.154 | | 0.165 | 0.087 | 3.999 | 19.851 |
| 13.0.2_1_On4 | On.Ar | Fossil | 0.012 | 0.008 | 0.037 | 0.020 | 0.353 | 0.130 | | 2.869 | 2.408 | 6.427 | 8.141 |
| 13.0.2_1_On4 | In.Ar | Fossil | 0.018 | 0.011 | 0.049 | 0.033 | 0.464 | 0.179 | | 0.723 | 0.819 | 5.645 | 53.172 |
| 32_6_On3 | On.Ar | Fossil | 0.021 | 0.014 | 0.052 | 0.043 | 0.600 | 0.320 | | 0.070 | 0.018 | 2.081 | 1.787 |
| 32_6_On3 | In.Ar | Fossil | 0.023 | 0.009 | 0.046 | 0.023 | 0.674 | 0.213 | | 0.175 | 0.143 | 2.175 | 2.995 |
| 32_6_On4 | On.Ar | Fossil | 0.017 | 0.012 | 0.038 | 0.020 | 0.887 | 0.542 | | 0.294 | 0.179 | 1.107 | 0.681 |
| 32_6_On4 | In.Ar | Fossil | 0.038 | 0.056 | 0.057 | 0.041 | 1.244 | 0.436 | | 0.188 | 0.103 | 1.229 | 0.990 |
| 4.0.1_1_On1.2 | On.Ar | Fossil | 0.045 | 0.018 | 0.062 | 0.037 | 1.134 | 0.723 | | 0.248 | 0.093 | 0.966 | 0.297 |
| 4.0.1_1_On1.2 | In.Ar | Fossil | 0.075 | 0.022 | 0.083 | 0.029 | 1.224 | 2.088 | | 0.308 | 0.080 | 0.945 | 0.091 |
| 4.0.1_1_On2 | On.Ar | Fossil | 0.028 | 0.012 | 0.059 | 0.037 | 0.700 | 0.188 | | 0.111 | 0.078 | 1.337 | 0.512 |
| 4.0.1_1_On2 | In.Ar | Fossil | 0.029 | 0.011 | 0.053 | 0.025 | 0.761 | 0.528 | | 0.171 | 0.163 | 1.379 | 0.635 |
| 4.1.5_1_On3 | On.Ar | Fossil | 0.022 | 0.011 | 0.074 | 0.053 | 0.531 | 0.151 | | 0.981 | 2.075 | 5.544 | 15.075 |
| 4.1.5_1_On3 | In.Ar | Fossil | 0.011 | 0.011 | 0.022 | 0.018 | 1.362 | 3.484 | | 2.002 | 4.961 | 6.216 | 7.043 |
| 4.1.5_1_On4 | On.Ar | Fossil | 0.013 | 0.010 | 0.059 | 0.043 | 0.414 | 0.157 | | - | - | 4.779 | 6.308 |
| 4.1.5_1_On4 | In.Ar | Fossil | 0.018 | 0.009 | 0.049 | 0.022 | 0.523 | 0.179 | | - | - | 8.802 | 12.898 |
| 5.0.32_5_On3 | On.Ar | Fossil | 0.017 | 0.011 | 0.047 | 0.030 | 0.432 | 0.212 | | - | - | 11.930 | 17.793 |
| 5.0.32_5_On3 | In.Ar | Fossil | 0.016 | 0.011 | 0.046 | 0.031 | 0.523 | 0.172 | | 0.270 | 0.178 | 7.655 | 40.157 |
| 7.13.2_1_On3 | On.Ar | Fossil | 0.022 | 0.011 | 0.041 | 0.027 | 0.773 | 0.389 | | 0.153 | 0.103 | 2.781 | 5.693 |
| 7.13.2_1_On3 | In.Ar | Fossil | 0.031 | 0.009 | 0.059 | 0.016 | 0.623 | 0.108 | | 0.098 | 0.027 | 2.562 | 2.563 |
| 7.13.2_1_On4 | On.Ar | Fossil | 0.034 | 0.011 | 0.058 | 0.020 | 0.621 | 0.155 | | 0.136 | 0.105 | 2.082 | 6.161 |
| 7.13.2_1_On4 | In.Ar | Fossil | 0.035 | 0.011 | 0.064 | 0.028 | 0.788 | 0.376 | | 0.149 | 0.097 | 1.214 | 0.307 |
| 7.8.4_1_On1 | On.Ar | Fossil | 0.009 | 0.006 | 0.048 | 0.019 | 0.244 | 0.128 | | - | - | 6.247 | 4.429 |
| 7.8.4_1_On1 | In.Ar | Fossil | 0.034 | 0.011 | 0.109 | 0.034 | 0.536 | 0.115 | | 0.310 | 0.146 | 9.119 | 31.738 |
| 7.8.4_1_On2 | On.Ar | Fossil | 0.021 | 0.011 | 0.046 | 0.031 | 0.647 | 0.137 | | 0.084 | 0.086 | 6.553 | 22.207 |
| 7.8.4_1_On2 | In.Ar | Fossil | 0.024 | 0.010 | 0.048 | 0.022 | 0.712 | 0.826 | | 0.158 | 0.143 | 4.034 | 40.293 |

LEBON, M., MÜLLER, K., BAHAIN, J.-J., FRÖHLICH, F., FALGUÈRES, C., BERTRAND, L., SANDT, C. & REICHE, I. 2011. Imaging fossil bone alterations at the microscale by SR-FTIR microspectroscopy. *Journal of Analytical Atomic Spectrometry,* 26.

REICHE, I., LEBON, M., CHADEFAUX, C., MÜLLER, K., LE HÔ, A.-S., GENSCH, M. & SCHADE, U. 2010. Microscale imaging of the preservation state of 5,000-year-old archaeological bones by synchrotron infrared microspectroscopy. *Analytical and Bioanalytical Chemistry,* 397**,** 2491-2499.

SCAGGION, C., DAL SASSO, G., NODARI, L., PAGANI, L., CARRARA, N., ZOTTI, A., BANZATO, T., USAI, D., PASQUALETTO, L., GADIOLI, G. & ARTIOLI, G. 2024. An FTIR-based model for the diagenetic alteration of archaeological bones. *Journal of Archaeological Science,* 161.
