## Supplementary 2 for "Underwater caves preserve biochemical composition and histological structure in modern and fossil mammalian bones"

**Supplementary 2: Birefringence data and polarised light microscopy images for: *Underwater caves preserve biochemical composition and histological structure in modern and fossil mammalian bones***

Walker, MM, Miszkiewicz, JJ, Rowe, JM, Matheson, CD, Vongsvivut, J, Sims, NA and Louys J

Transmitted and polarised light microscopy was used to assess histological integrity of bones collected from wet, dry, and fossil depositional environments at Green Waterhole and Gouldens Hole, South Australia. A region of interest (ROI) strip was captured on a BX63, under 10x magnification.

### 1 Birefringence data collection

Birefringence was scored based on the birefringence index (BI), where thin sections viewed under polarised light were classified as those comparable to fresh bone (1), reduced birefringence (0.5) and absent (0) [1-3]. Table 1 presents these data for each specimen analysed in each depositional environment (wet, dry, fossil). Not all samples analysed in the current study were assessed for birefringence due to sampling issues in the production of histological thin section slides. Although images are provided for samples 13.0.2_1 and 7.13.2_1, a full ROI strip could not be assessed due to sample quality.

Table 1 Birefringence data across wet, dry, and fossil bone specimens.

|  |  | **Burial Condition** | **Sample No.** | **Taxon** | **Element** | **Bone type** | **BI** |
| --- | --- | --- | --- | --- | --- | --- | --- |
|  |  | Wet | GWW61 | Ovicapra | Metacarpal | Plexiform | 1 |
|  |  | Wet | GWW4 | Ovicapra | Tibia | Plexiform | 1 |
|  |  | Wet | GWW76 | Ovicapra | Radius | Plexiform/Haversian | 0.5 |
|  |  | Wet | GH09 | Ovicapra | Metatarsal | Plexiform | 0 |
|  |  | Wet | GH10 | Ovicapra | Metatarsal | Plexiform | 0.5 |
|  |  | Wet | GH19 | Ovicapra | Metatarsal | Plexiform | 1 |
|  |  | Wet | GH04 | *Macropus* | Rib | Haversian | 0.5 |
|  |  | Wet | GH18 | *Macropus* | Tibia | Haversian/Radial | 1 |
|  |  | Dry | GWS60 | Ovicapra | Tibia | Plexiform | 0.5 |
|  |  | Dry | GWS02 | Ovicapra | Tibia | Plexiform | 0.5 |
|  |  | Dry | GWS19 | Ovicapra | Tibia | Plexiform | 0.5 |
|  |  | Dry | GWS47 | Ovicapra | Metatarsal | Plexiform | NA |
|  |  | Dry | GWS48 | Ovicapra | Metatarsal | Plexiform | 1 |
|  |  | Dry | GWS69 | Ovicapra | Metatarsal | Plexiform | 1 |
|  |  | Fossil | 5.0.32_5 | Sthenurine | Tibia | Haversian/Radial | 0.5 |
|  |  | Fossil | 13.0.2_1 | *Macropus* | Tibia | Haversian/Radial | NA |
|  |  | Fossil | 7.13.2_1 | Sthenurine | Tibia | Haversian/Radial | NA |
|  |  | Fossil | 7.8.4_1 | *Macropus* | Tibia | Haversian/Radial | 1 |
|  |  | Fossil | 4.1.5_1 | Sthenurine | Tibia | Haversian/Radial | 0.5 |
|  |  | Fossil | 32_11 | Sthenurine | Tibia | Haversian/Radial | 0.5 |
|  |  | Fossil | 32_6 | *Macropus* | Tibia | Haversian/Radial | 1 |
|  |  | Fossil | 12.0.1_7 | *Macropus* | Tibia | Haversian/Radial | 1 |
|  |  | Fossil | 4.0.1_1 | *Notamacropus* | Tibia | Haversian/Radial | NA |

Table 2 Birefringence summary table across depositional environments

|  | **Birefringence Index (BI)** | | |
| --- | --- | --- | --- |
| **Depositional environment** | **0** | **0.5** | **1** |
| Wet (n=8) | **1** | **3** | **4** |
| Dry (n=5) | **0** | **3** | **2** |
| Fossil (n=6) | **0** | **3** | **3** |

### 2 Light microscopy data

#### 2.1 Wet

##### GWW61

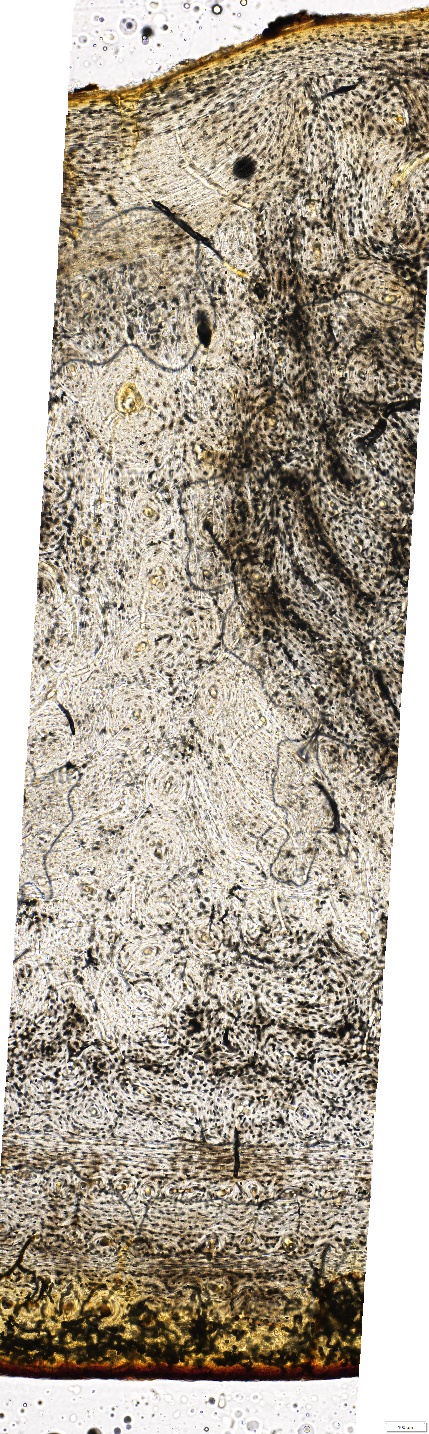

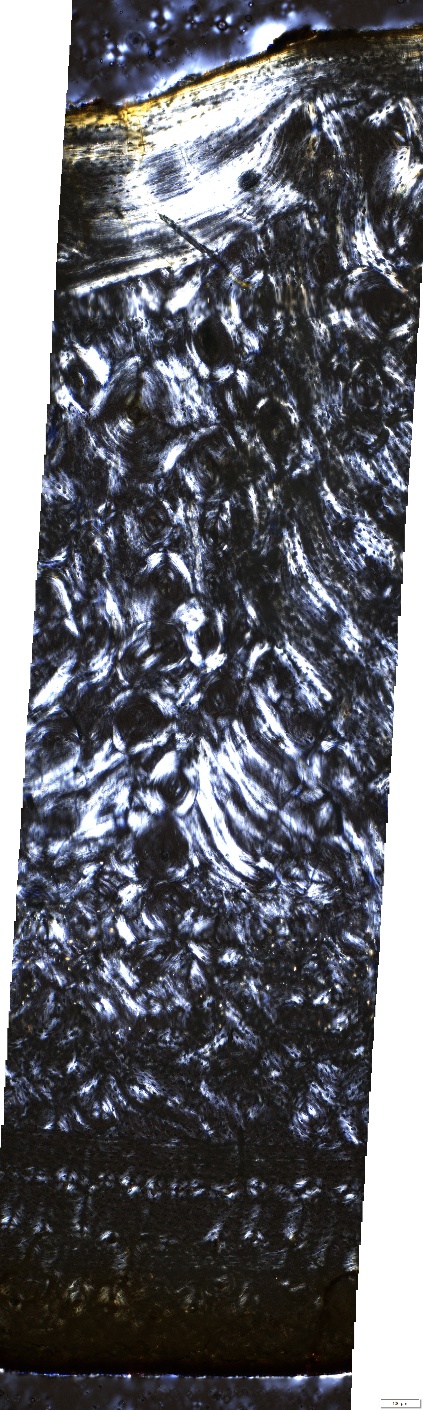

##### GWW4

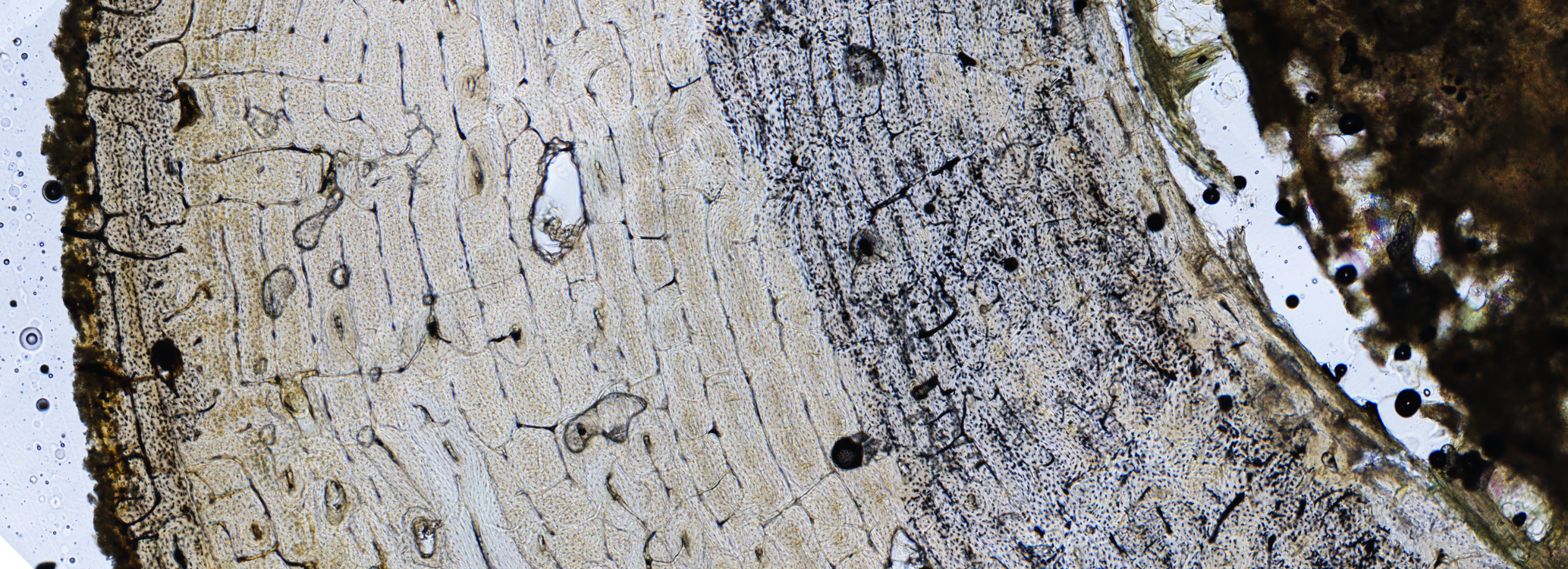

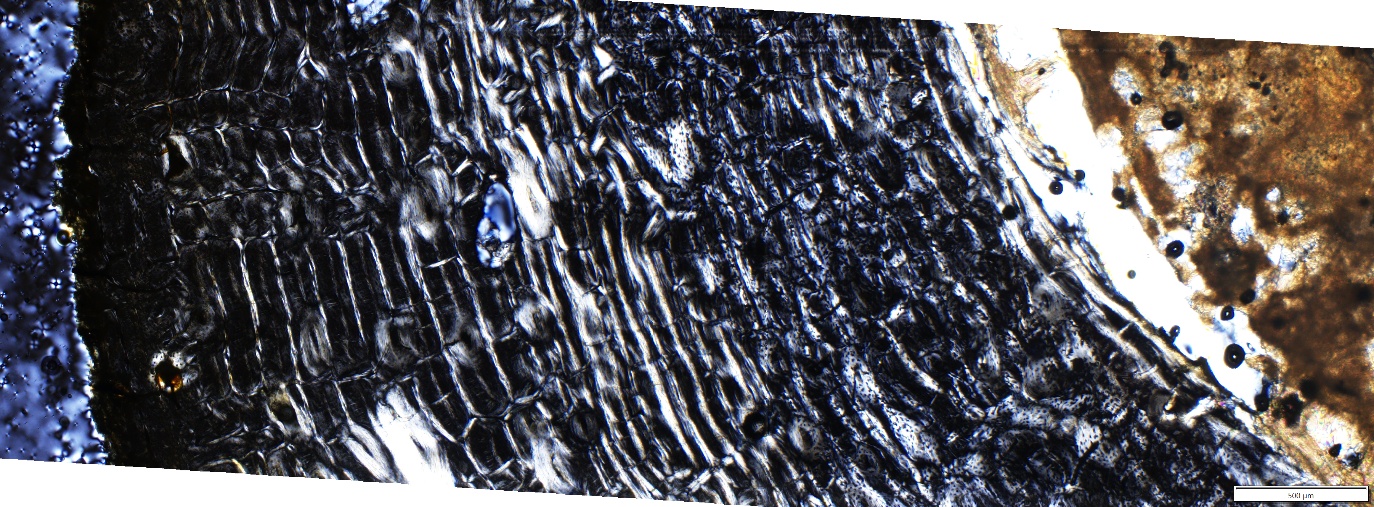

##### GWW76

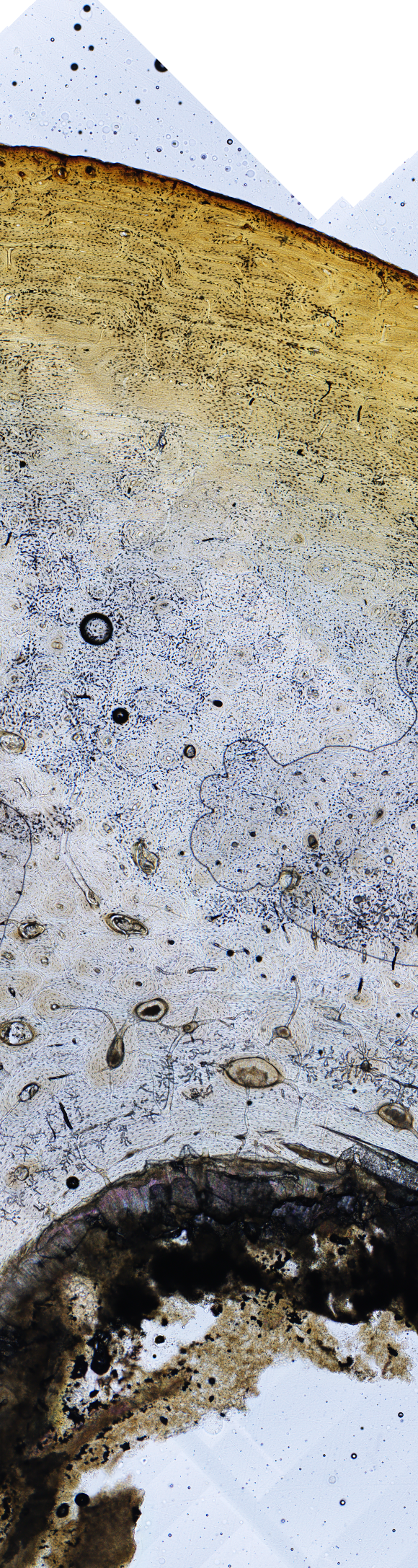

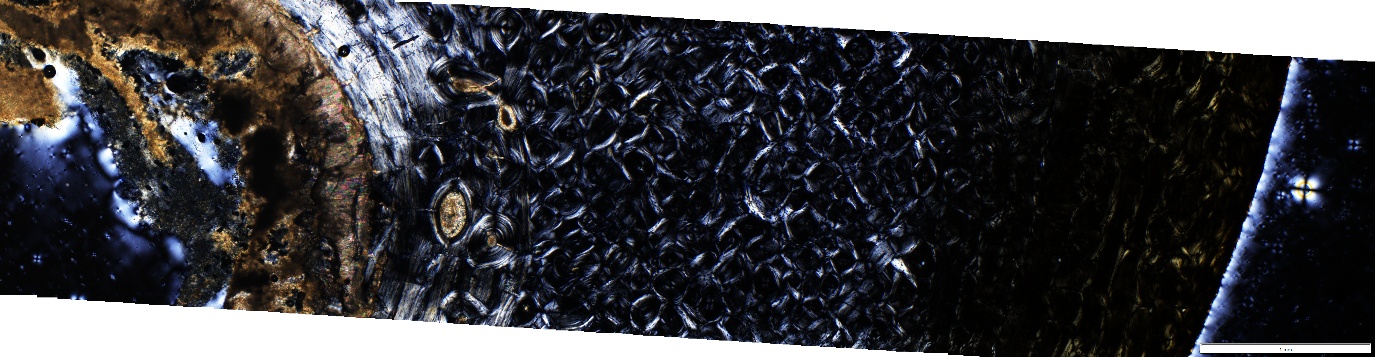

### GH04

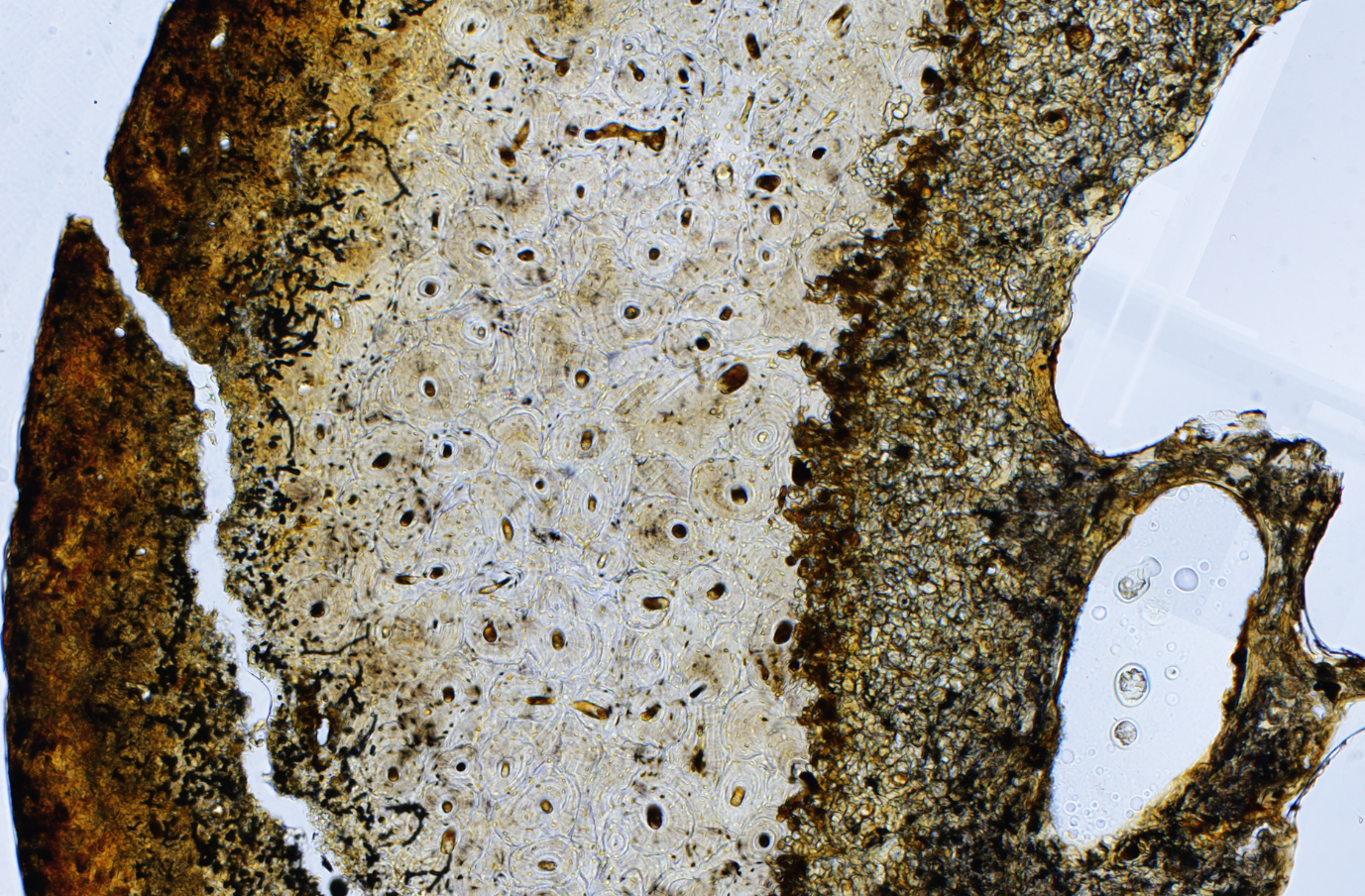

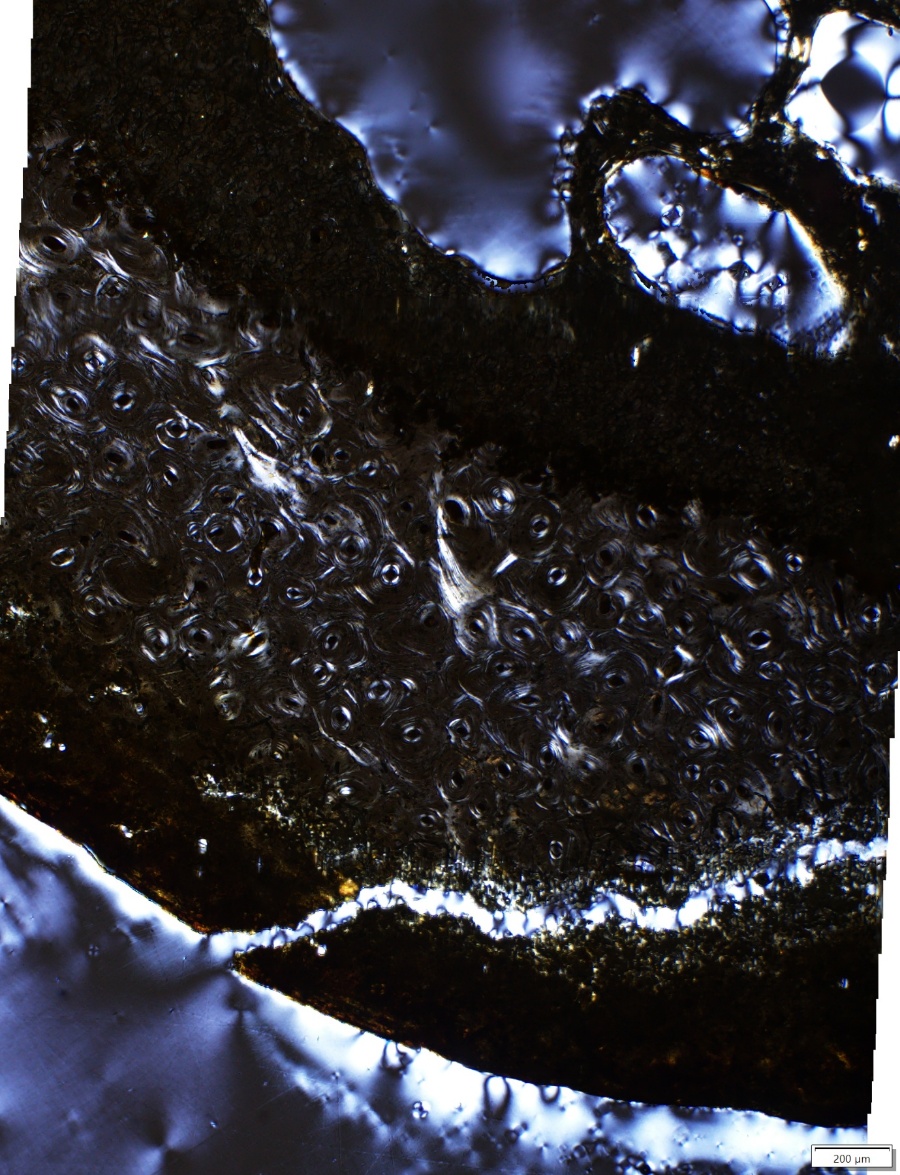

### GH09

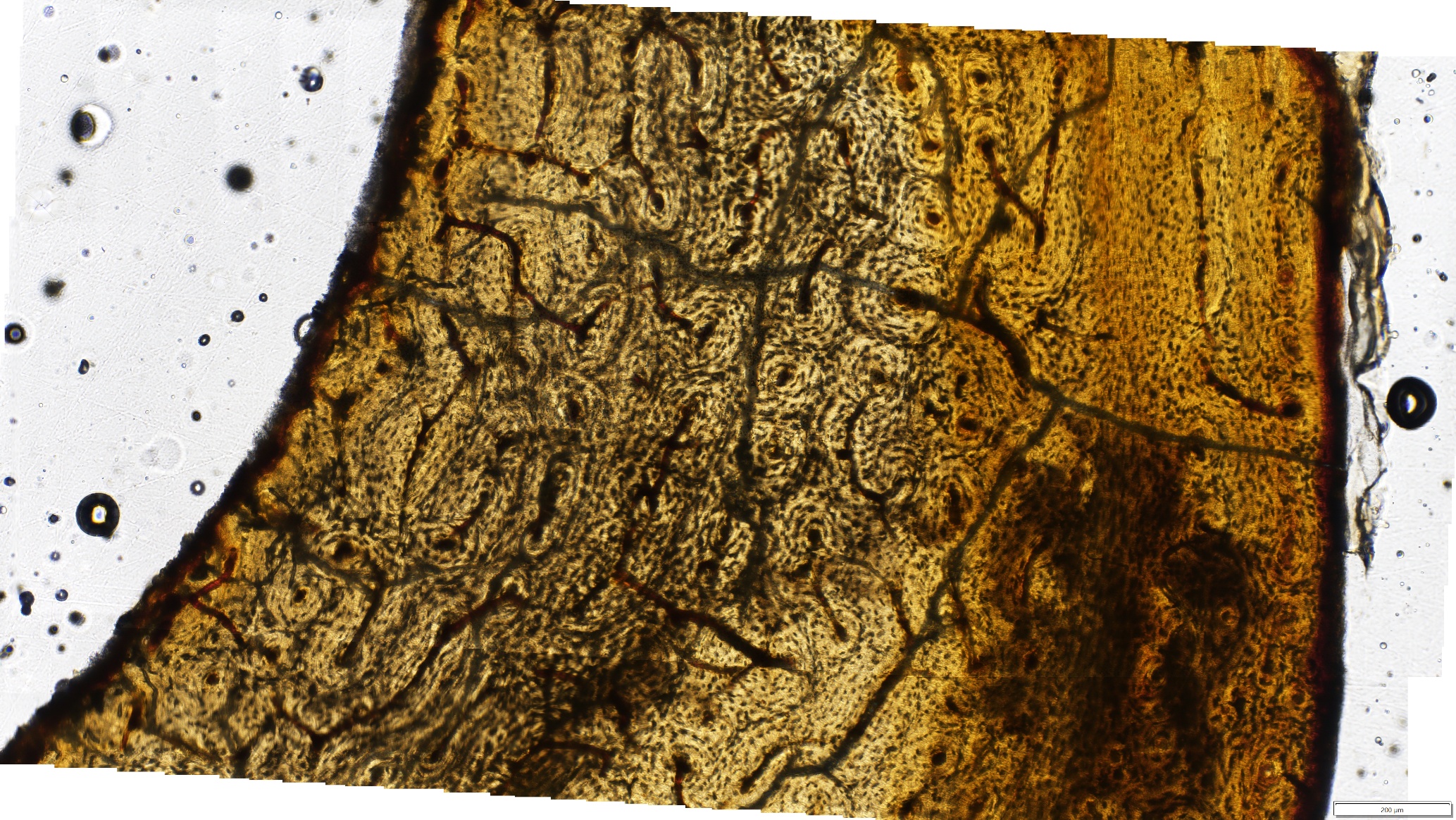

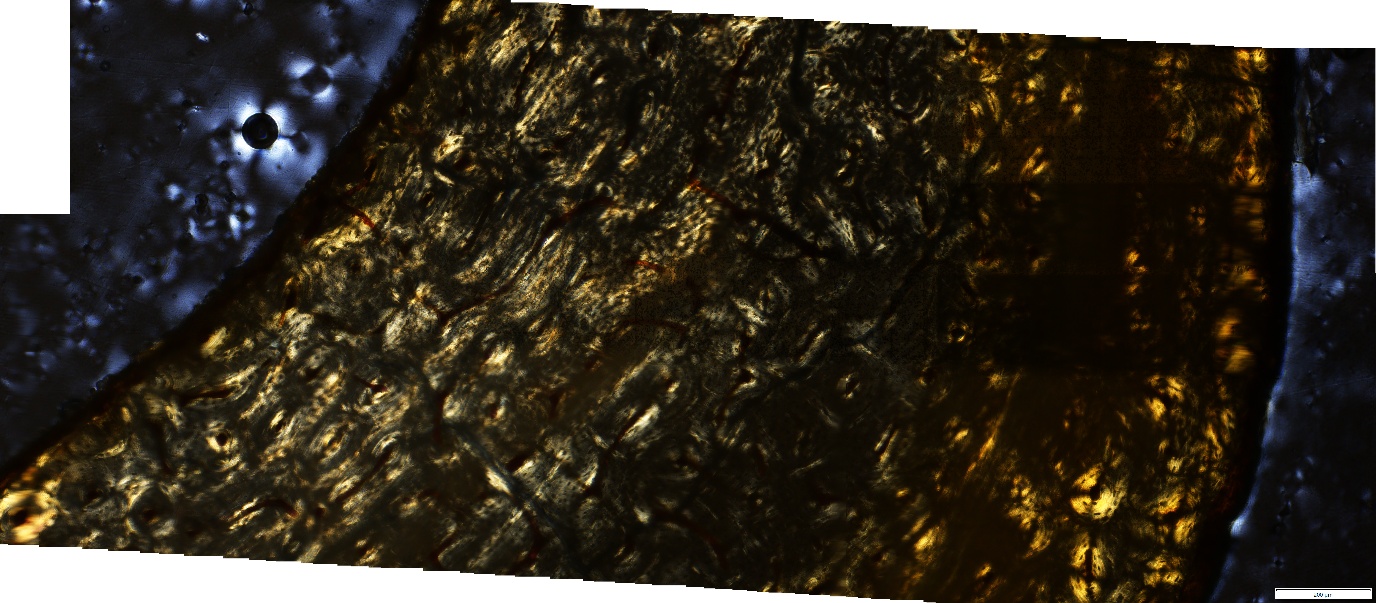

### GH10

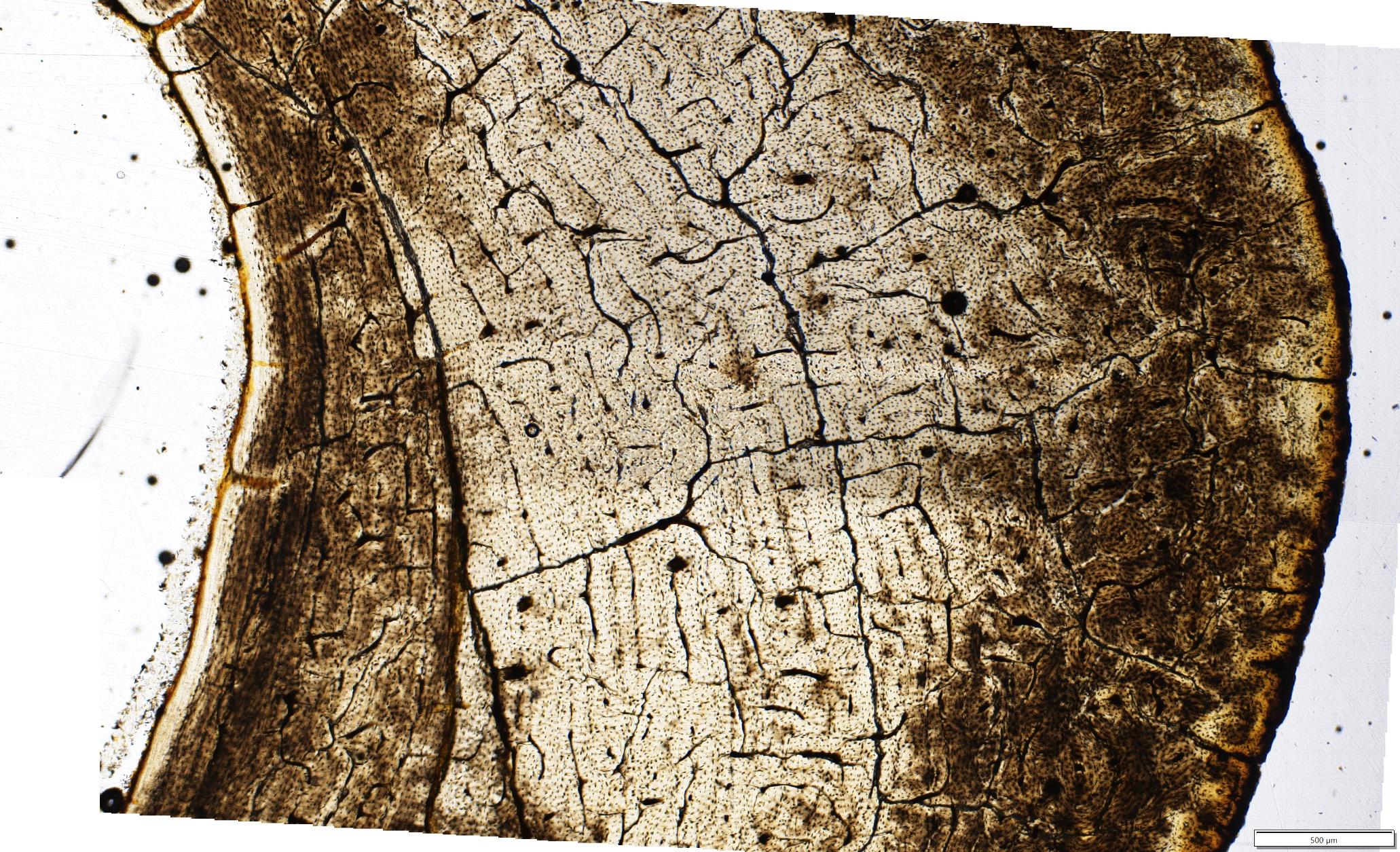

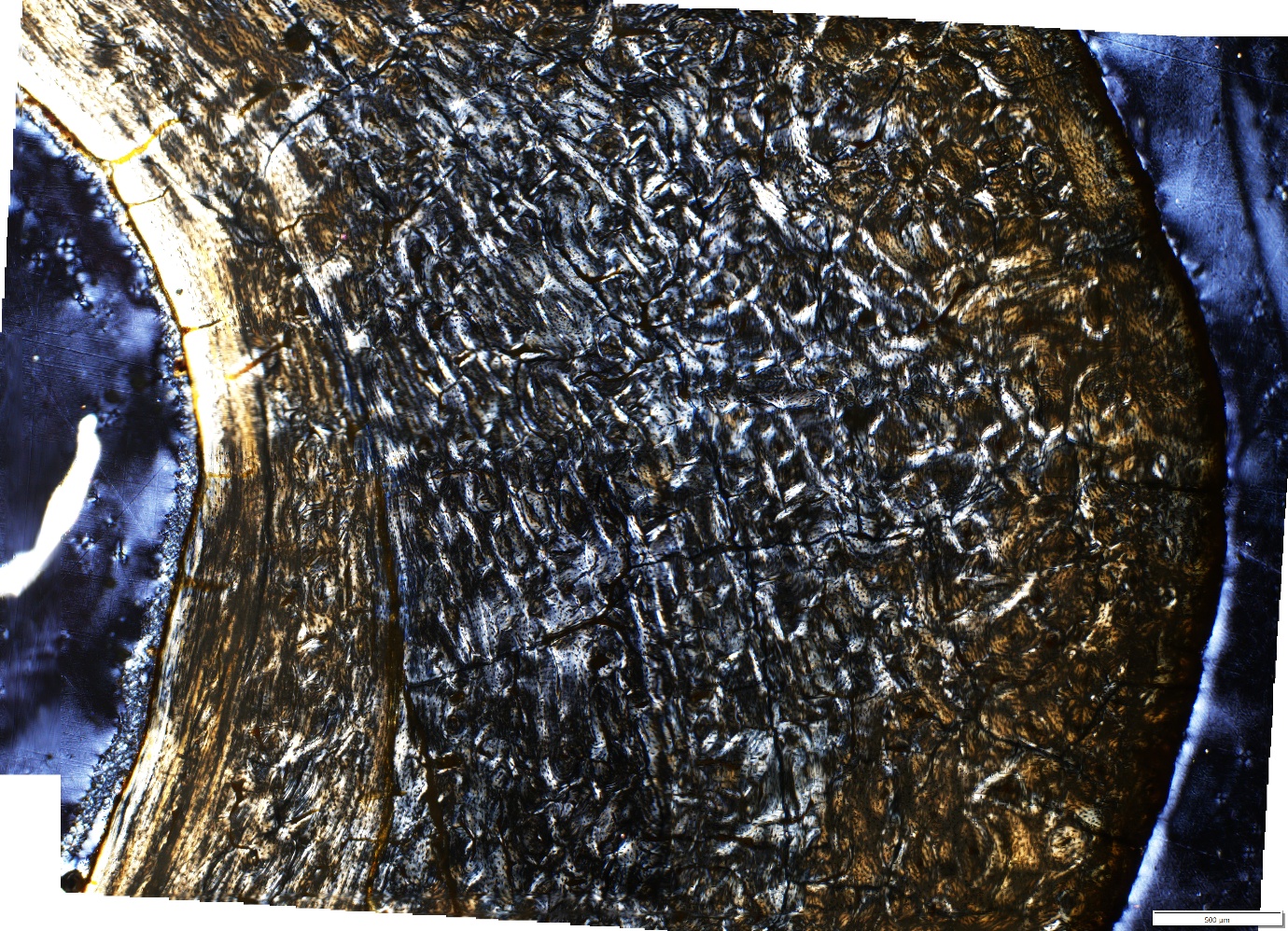

### GH18

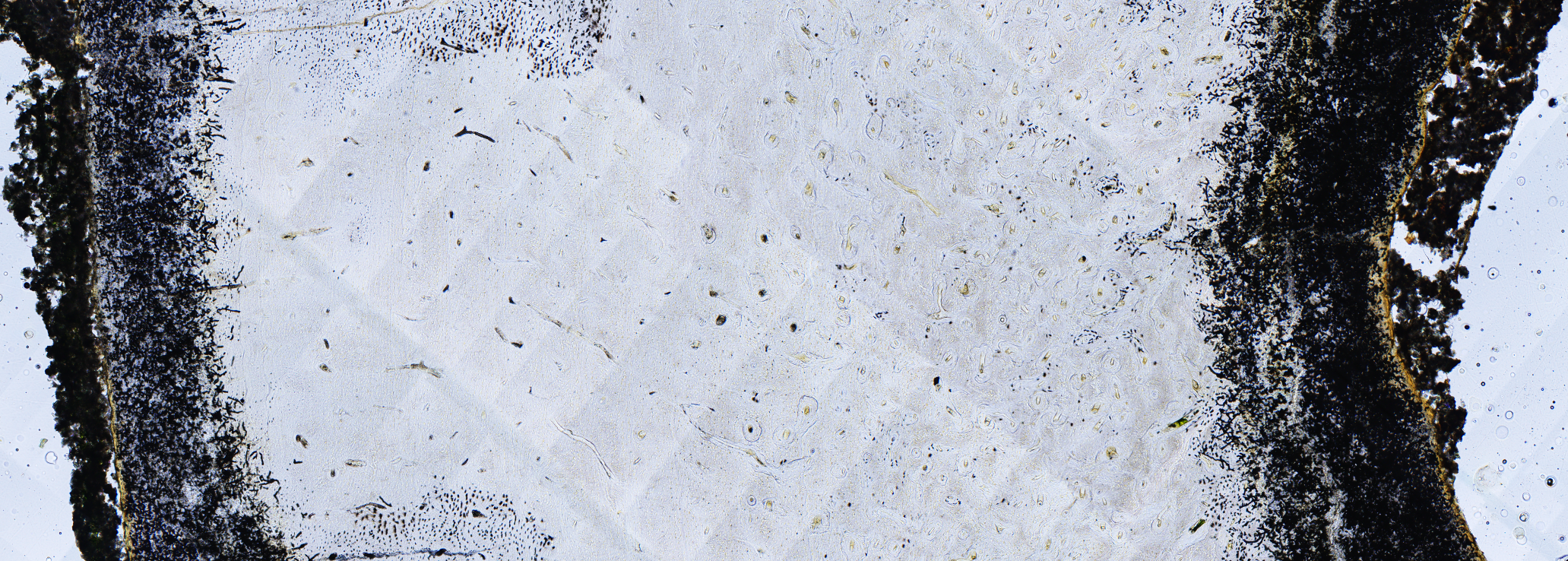

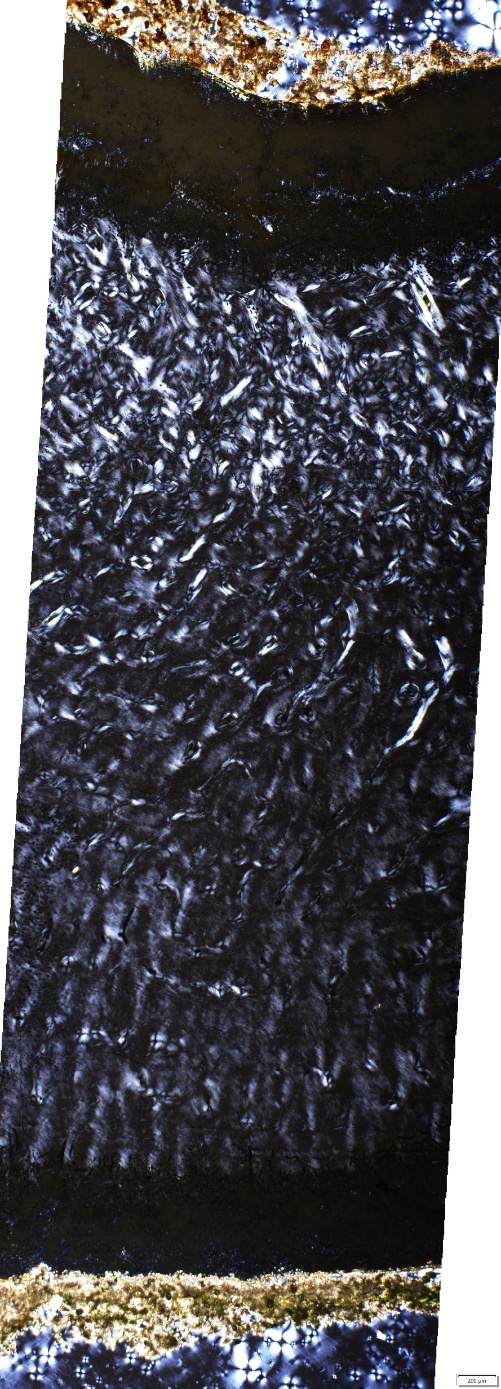

### GH19

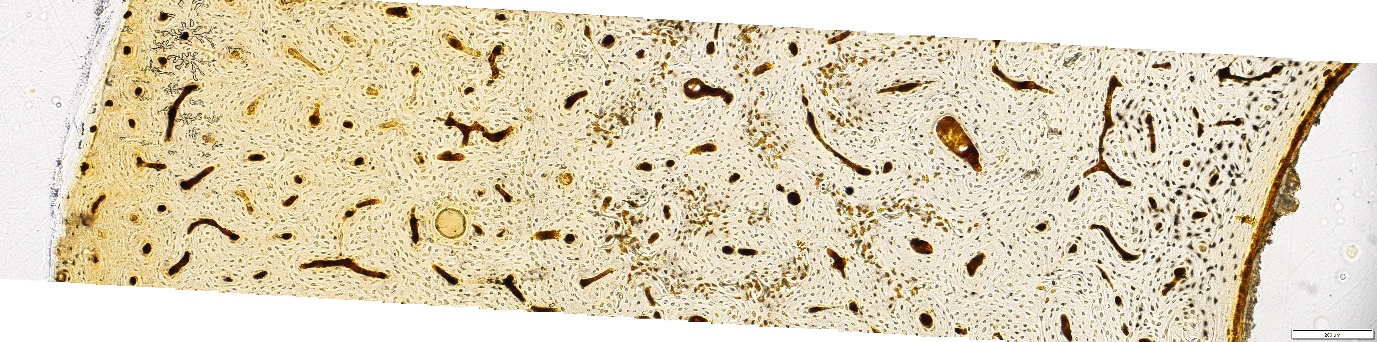

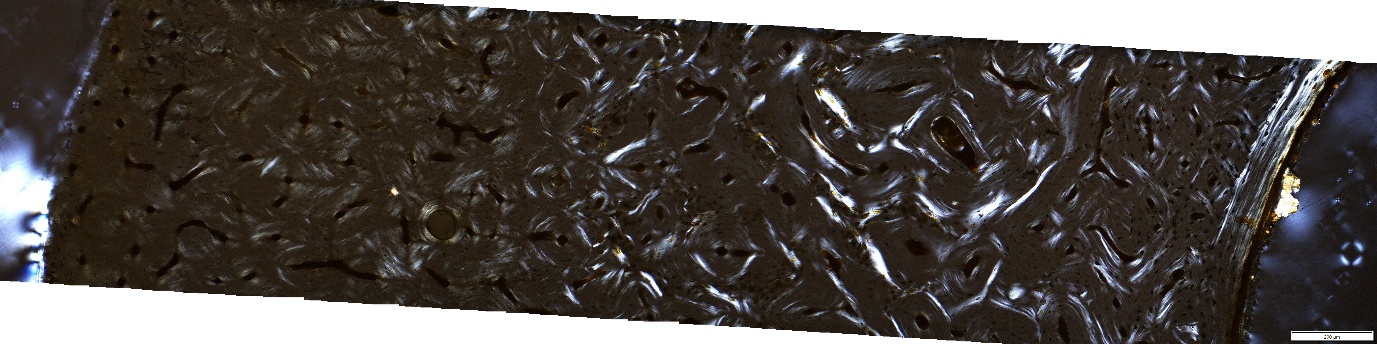

#### 2.2 Dry

##### GWS02

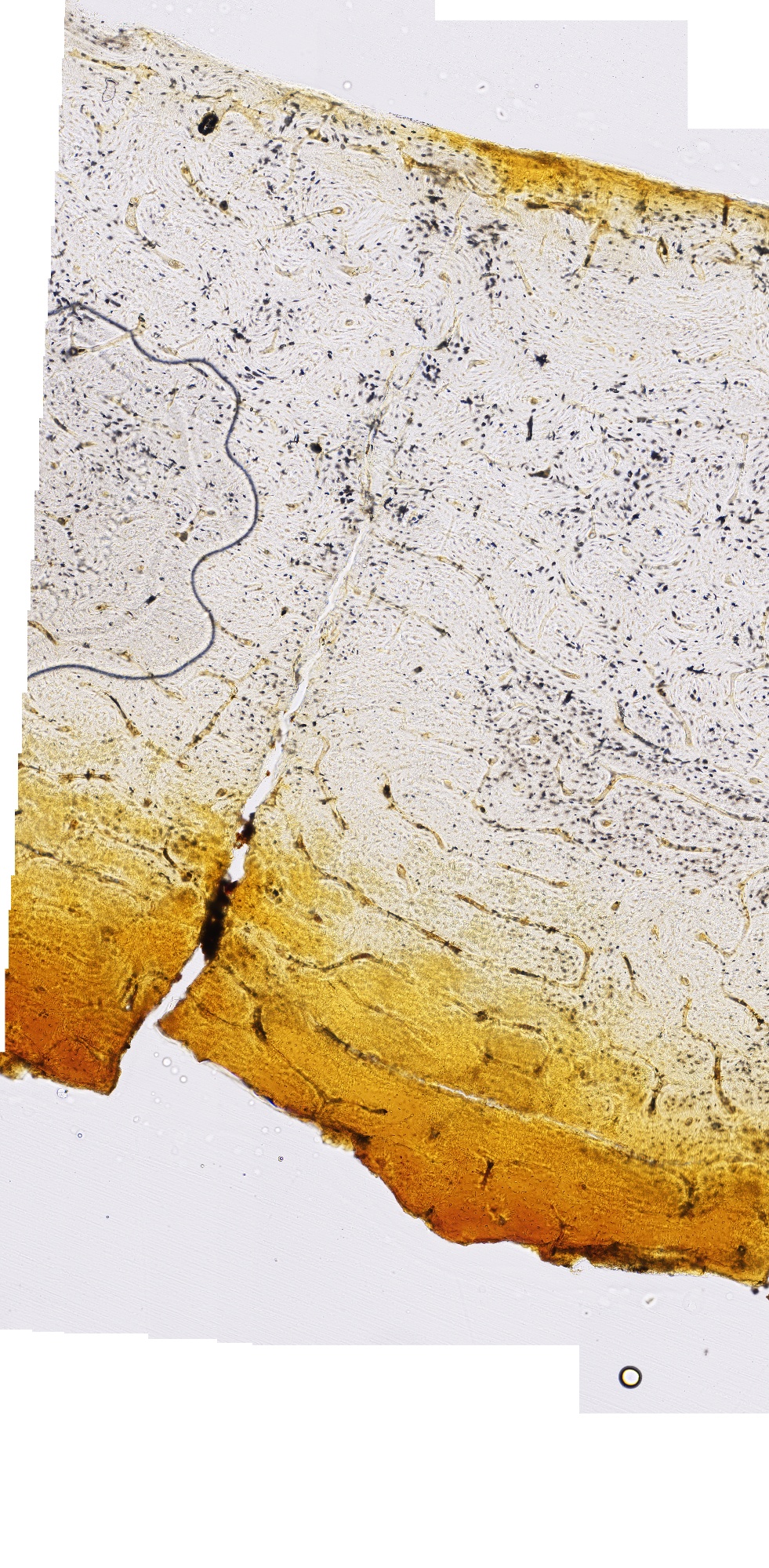

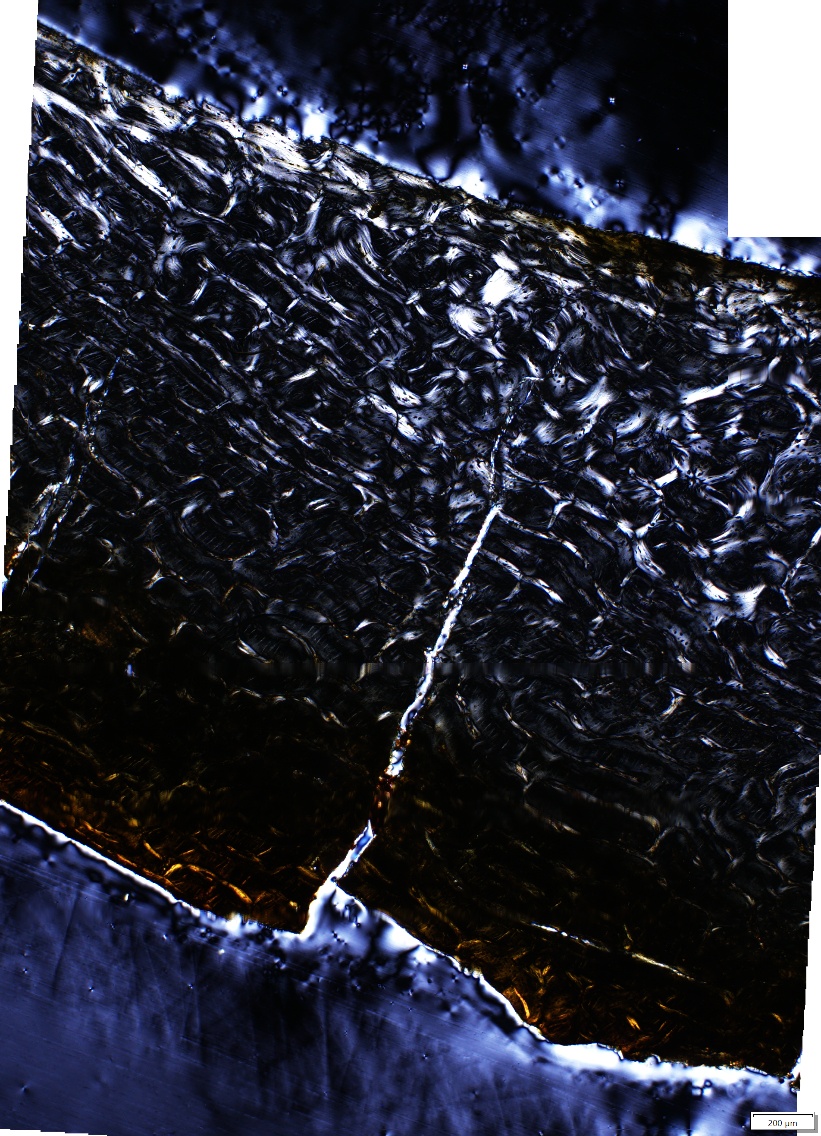

##### GWS19

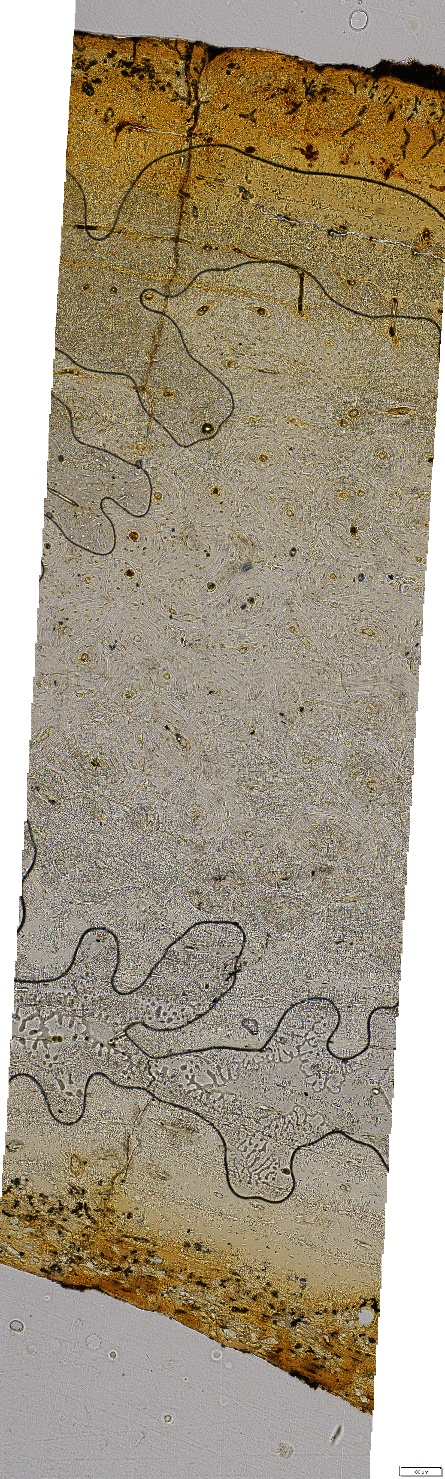

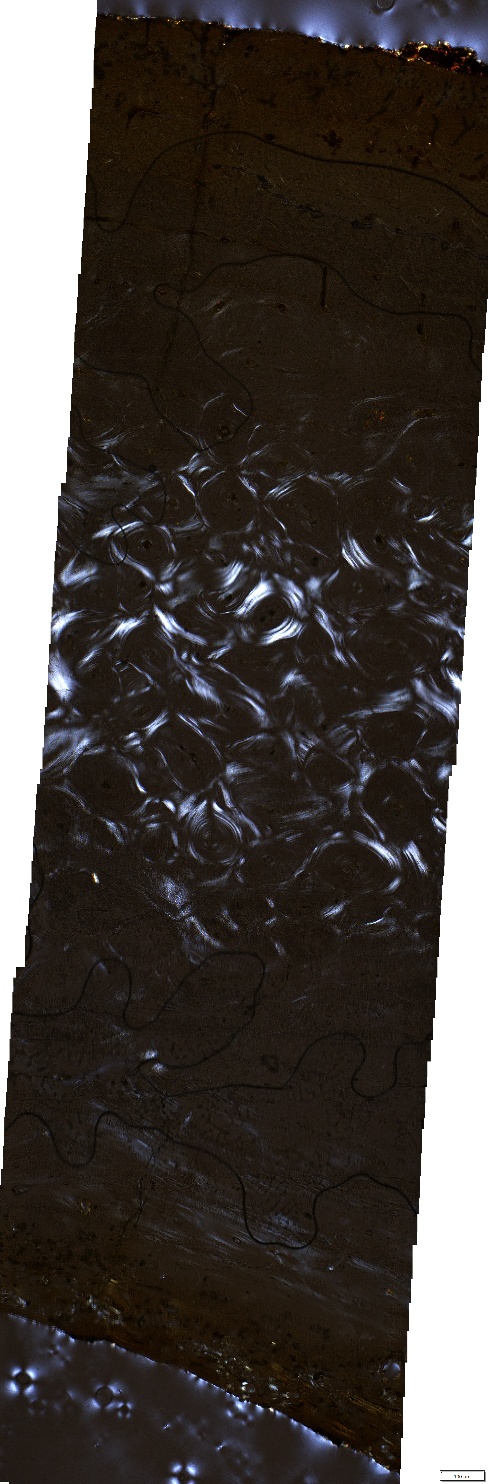

##### GWS48

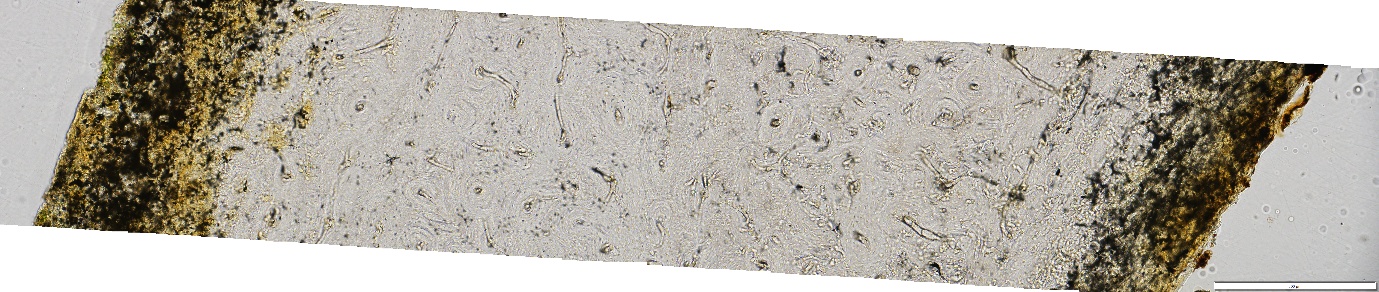

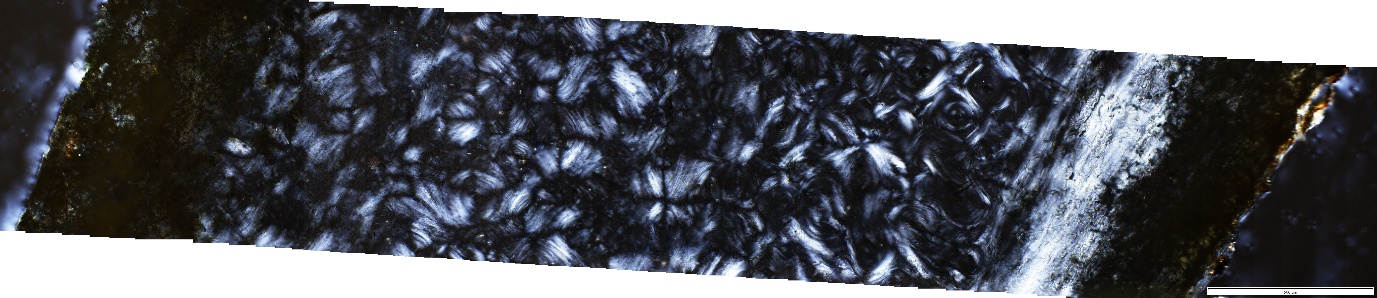

##### GWS69

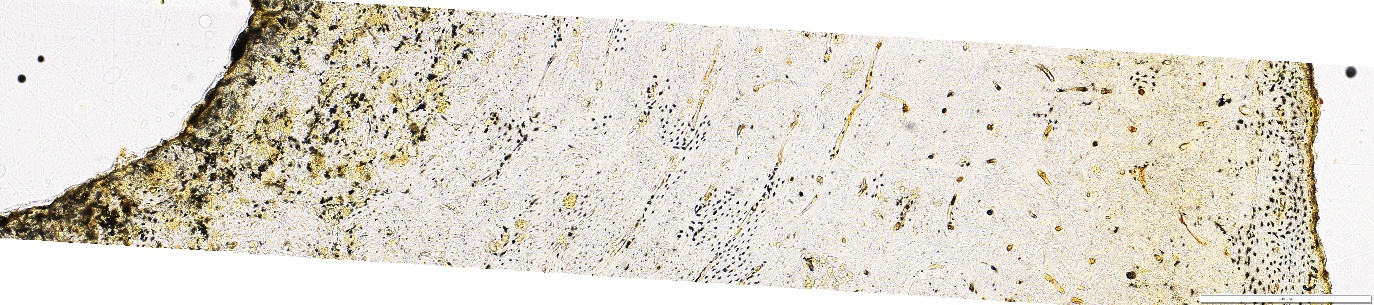

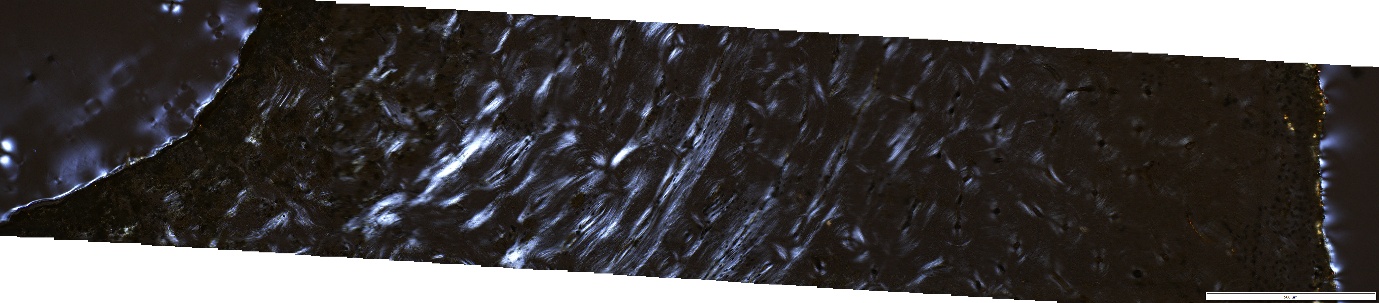

##### GWS60

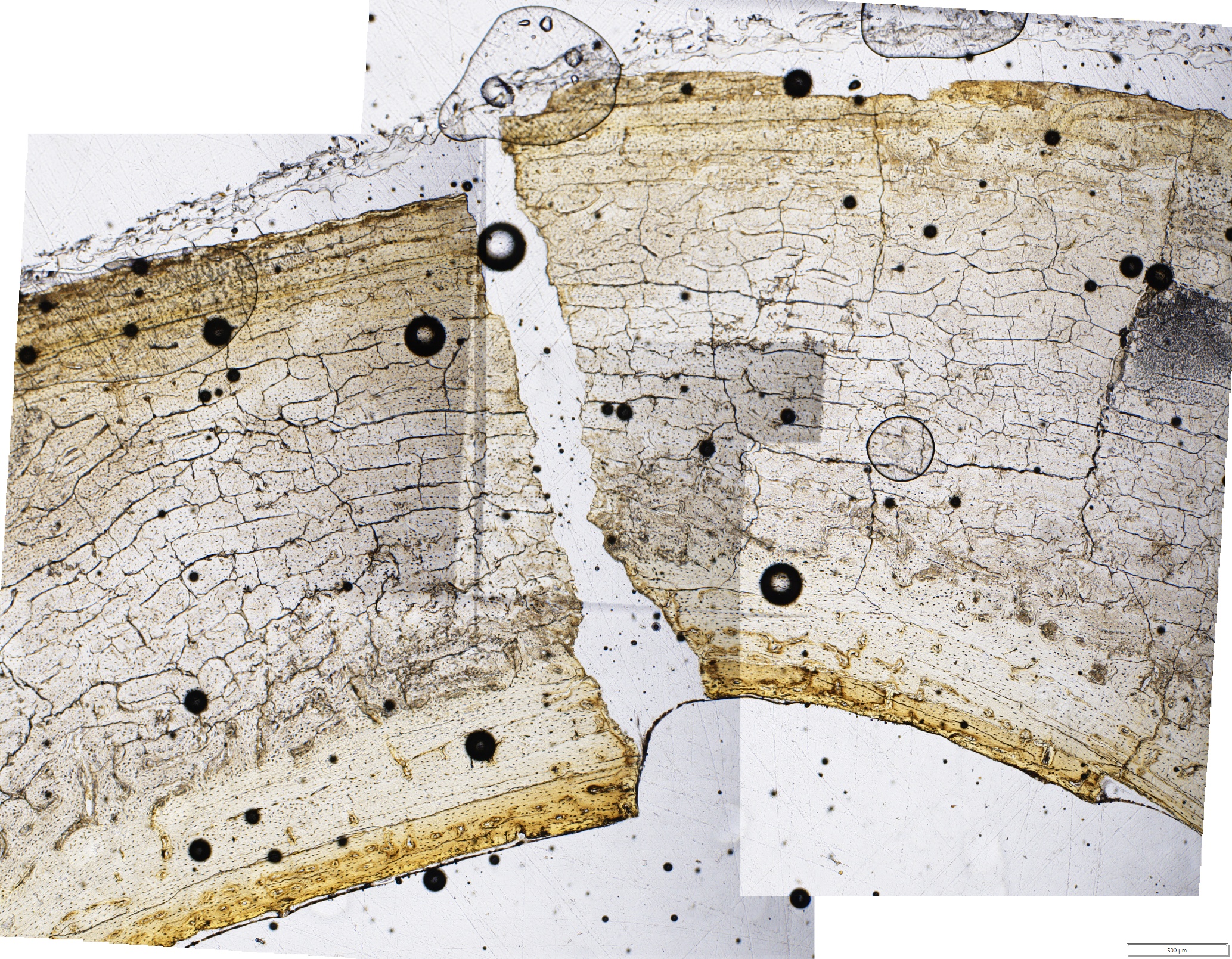

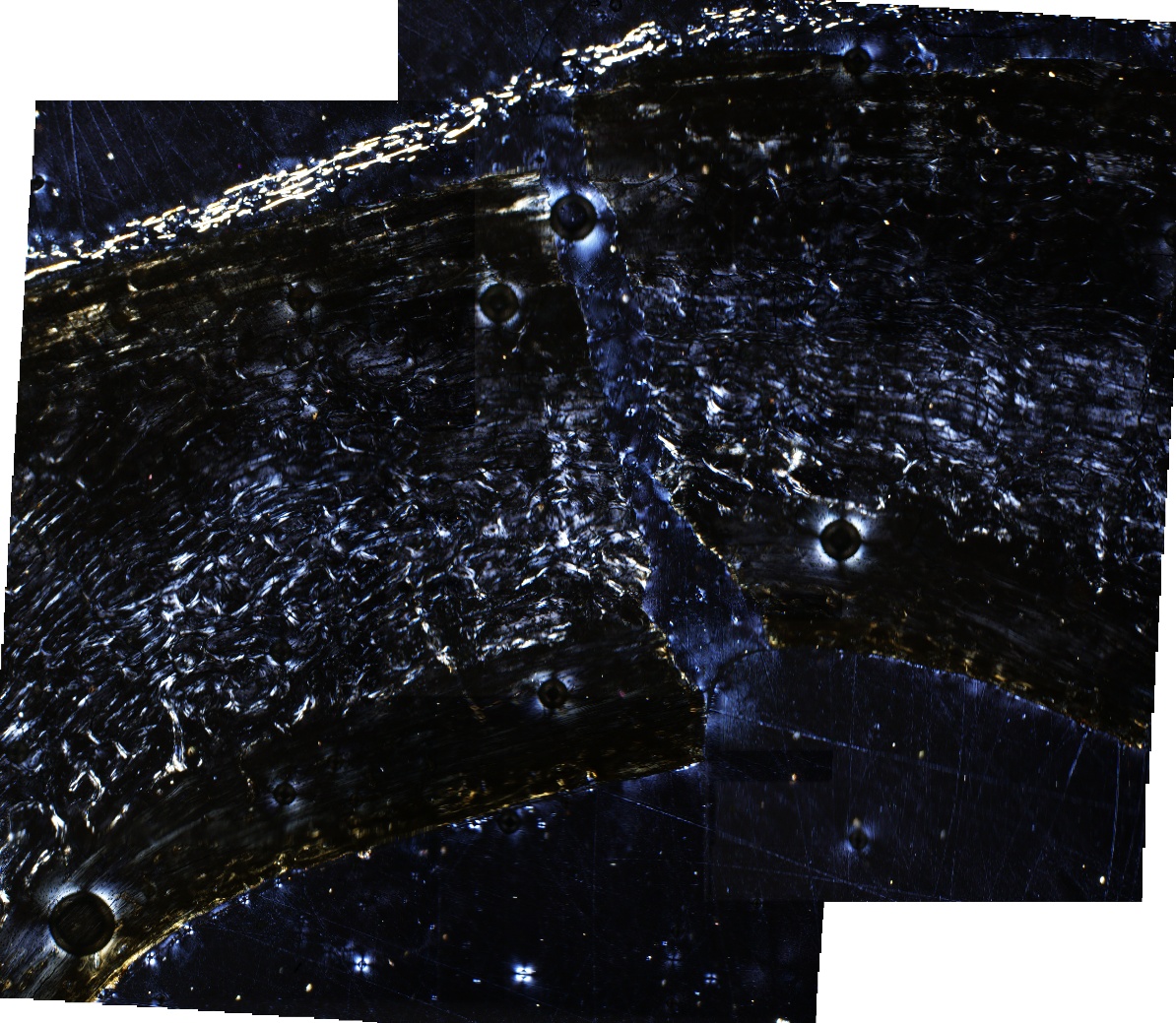

#### 2.3 Fossil

**5.0.32_5**

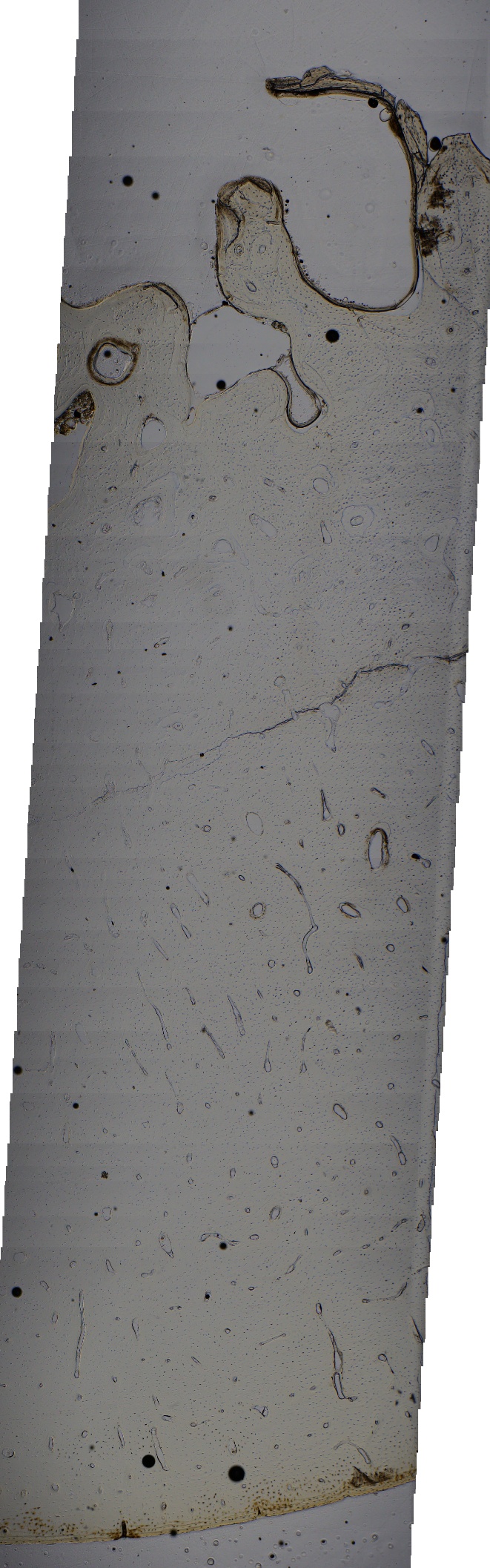

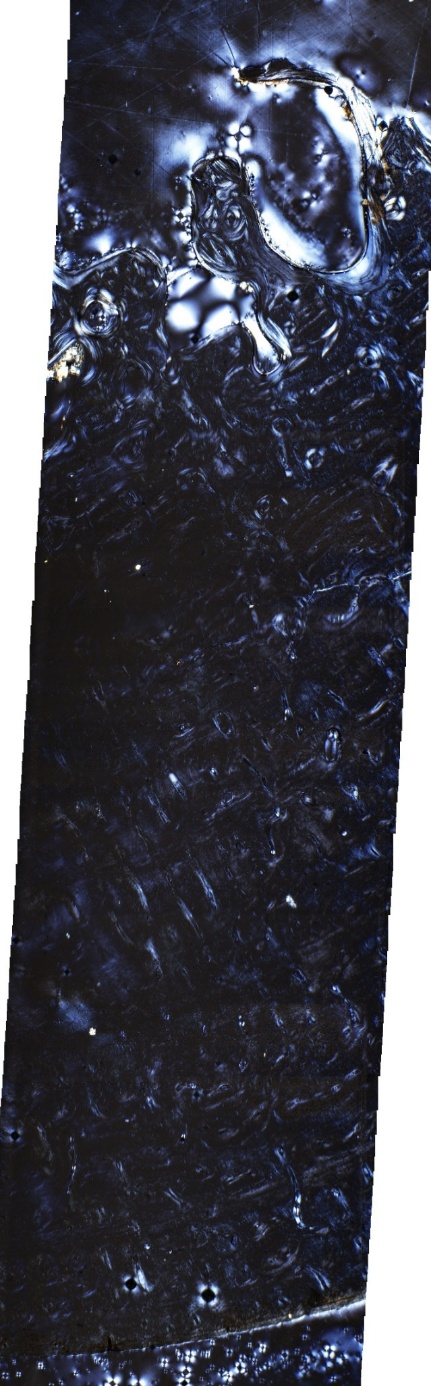

**13.0.2_1**

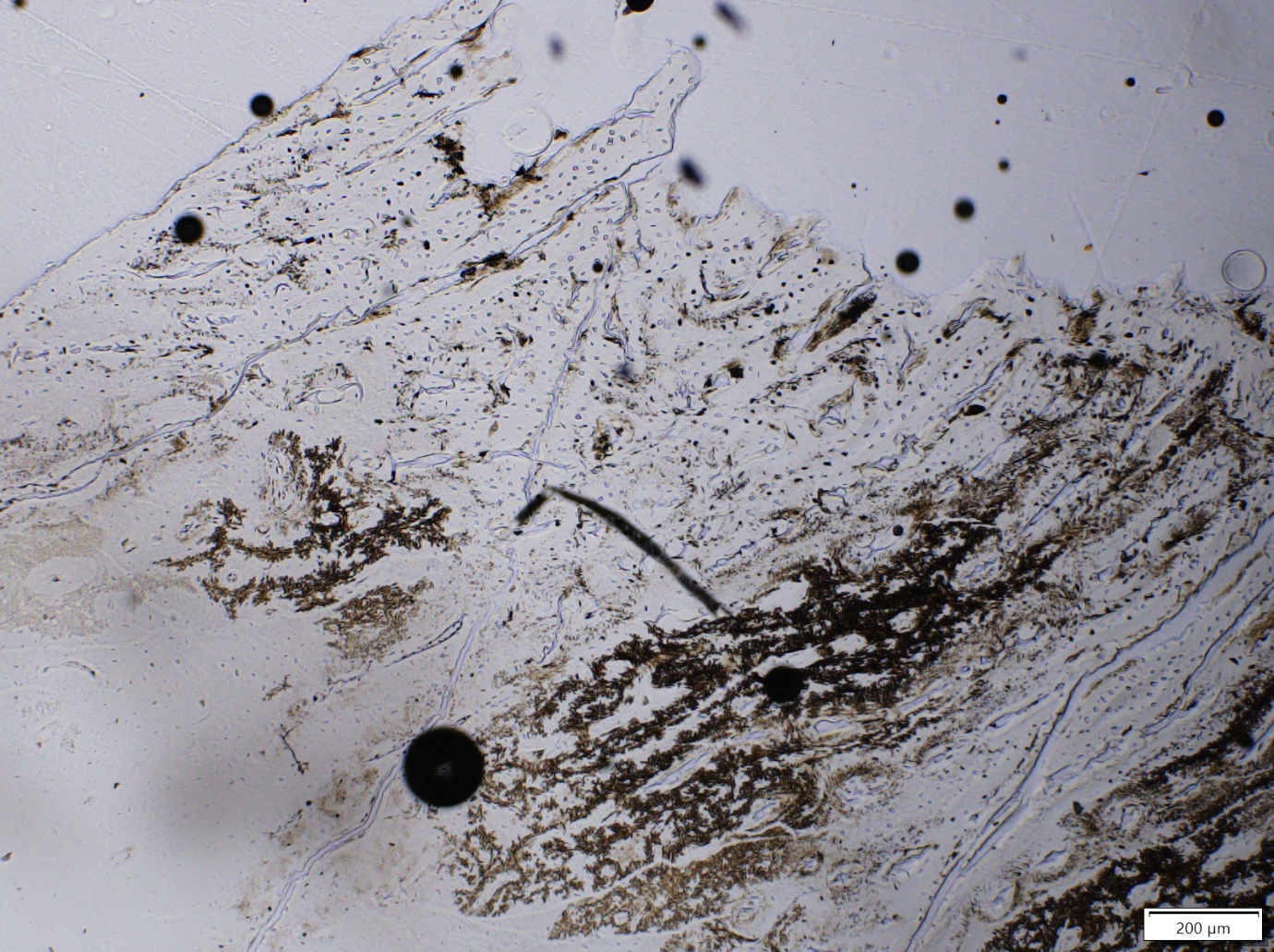

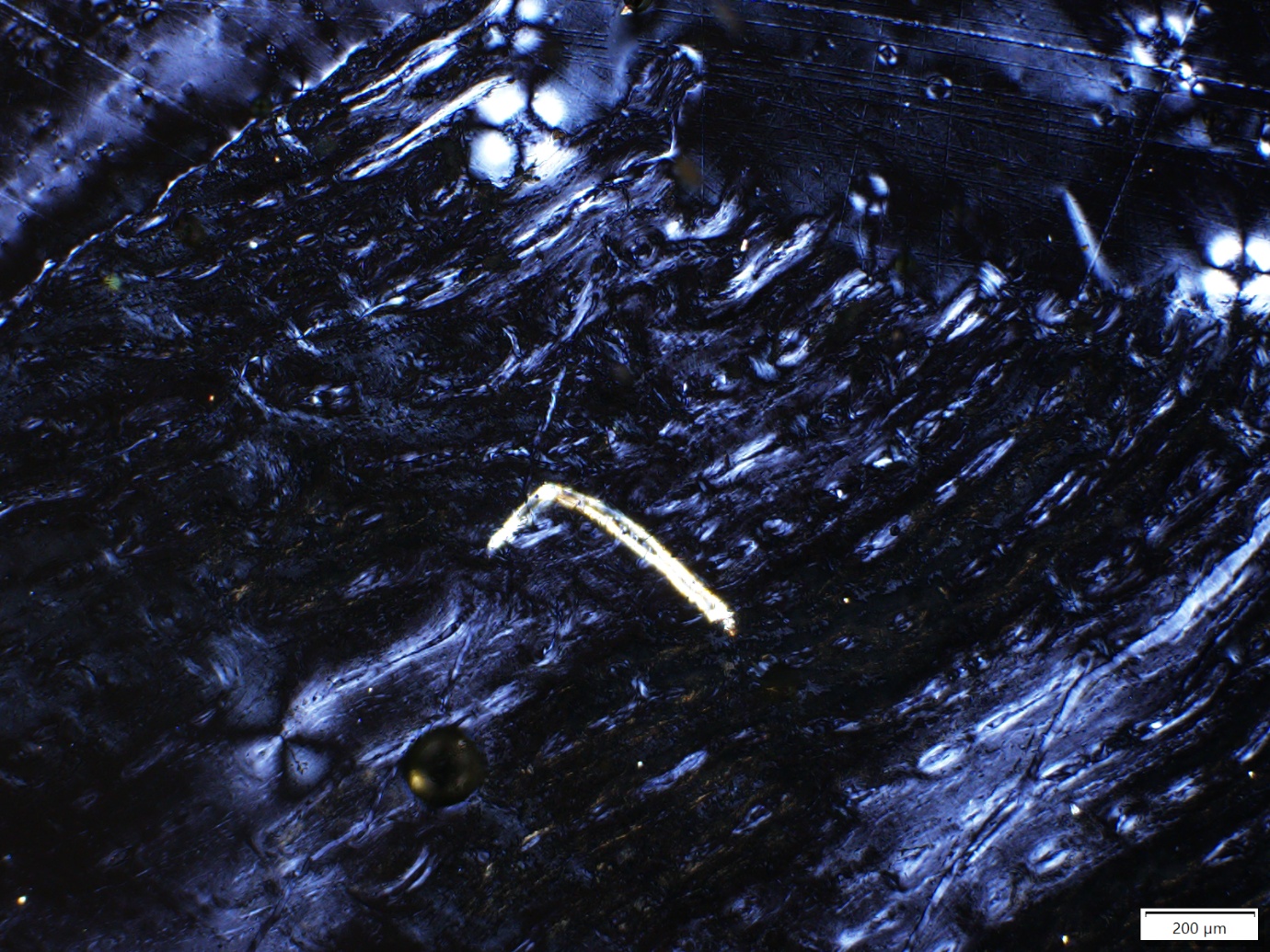

**7.13.2_1**

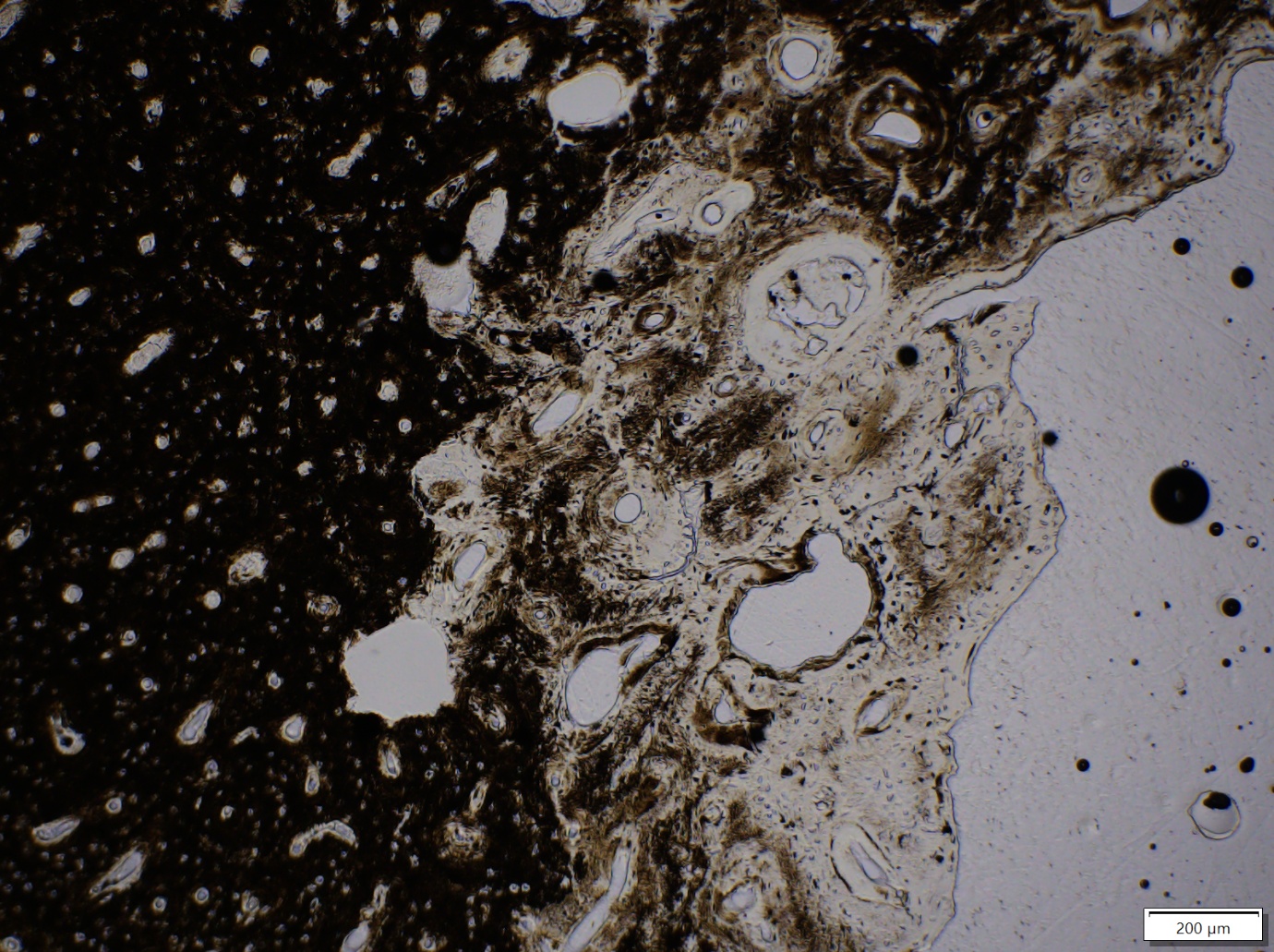

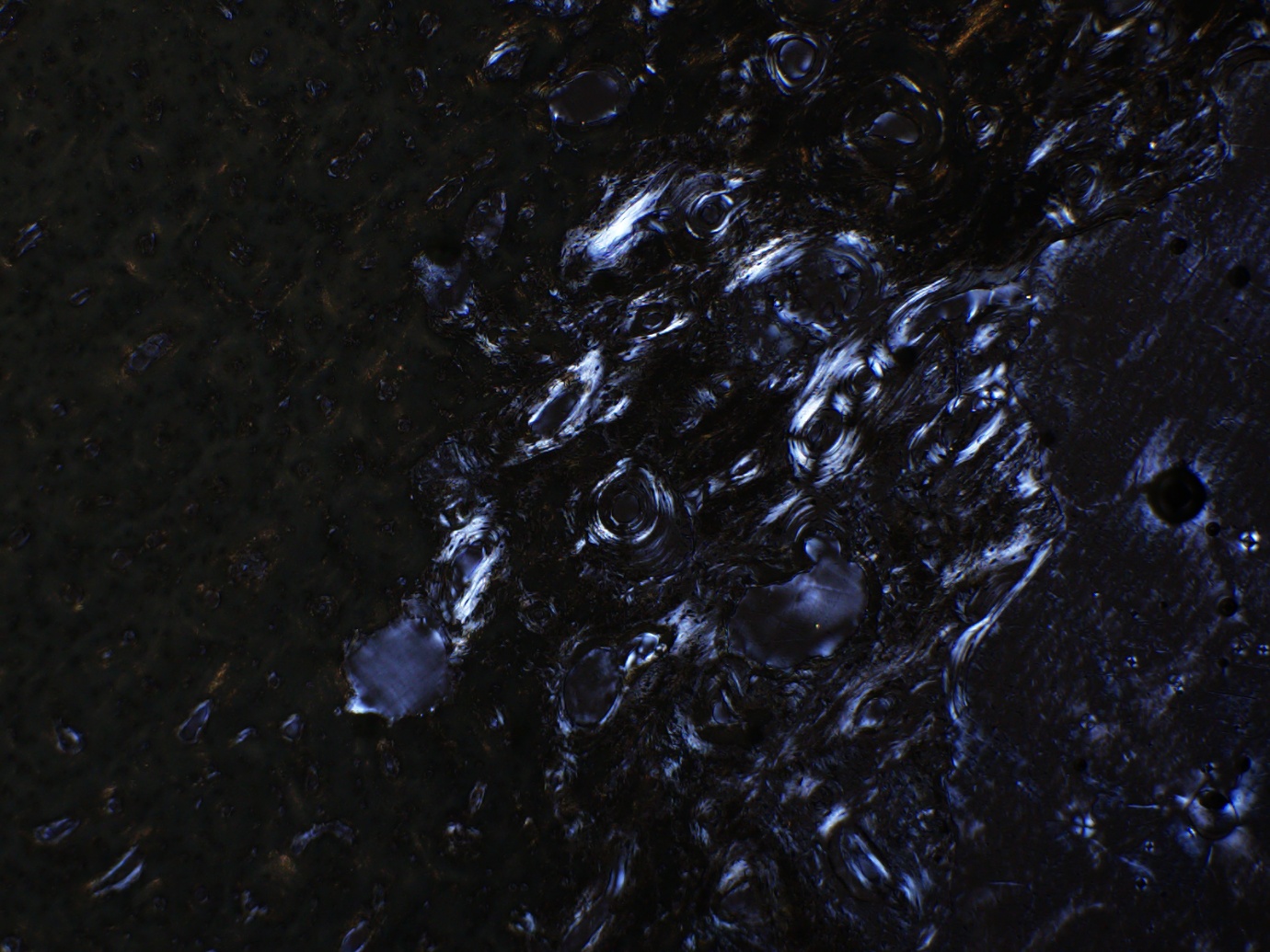

**7.8.4_1**

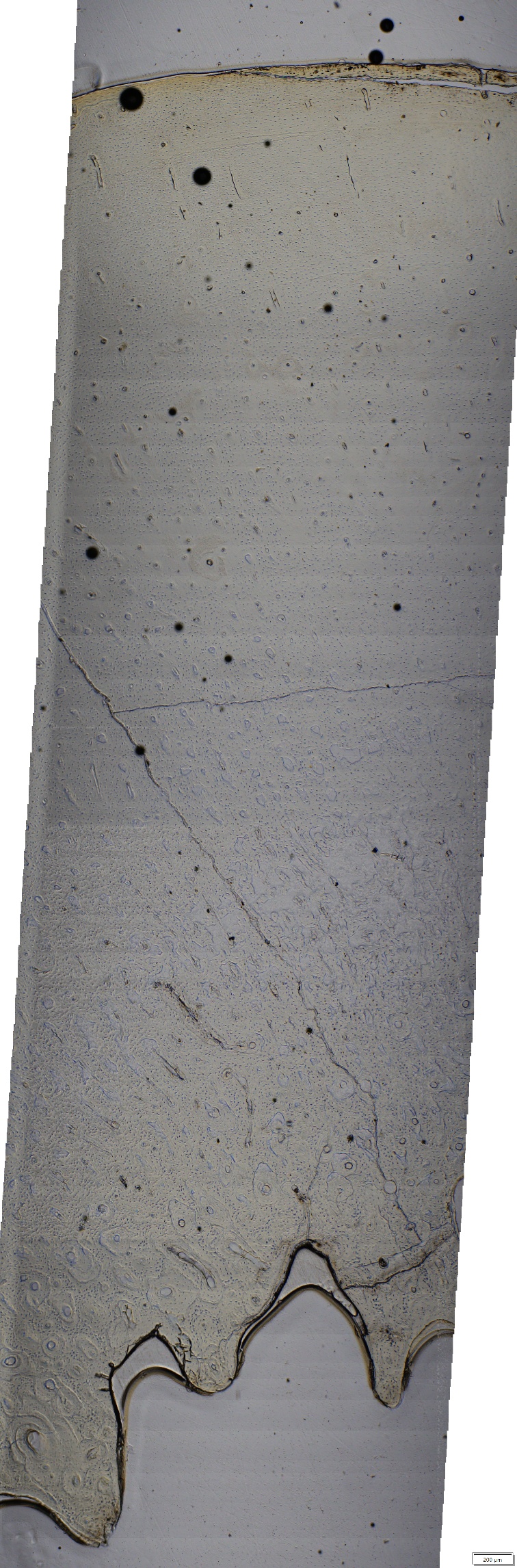

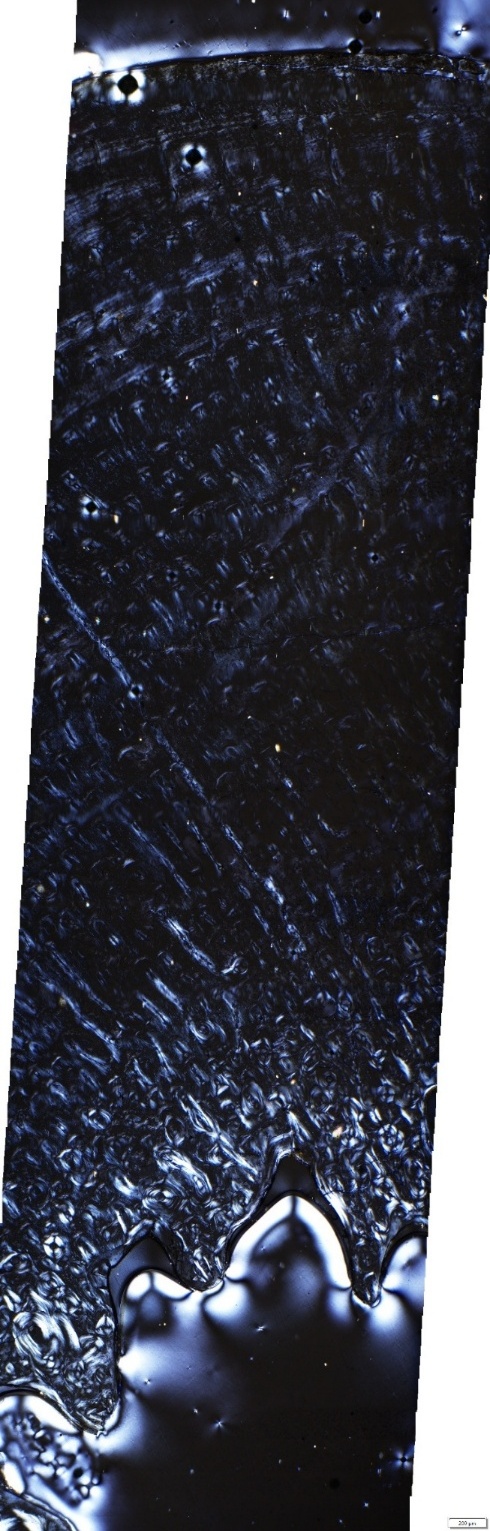

**4.1.5_1**

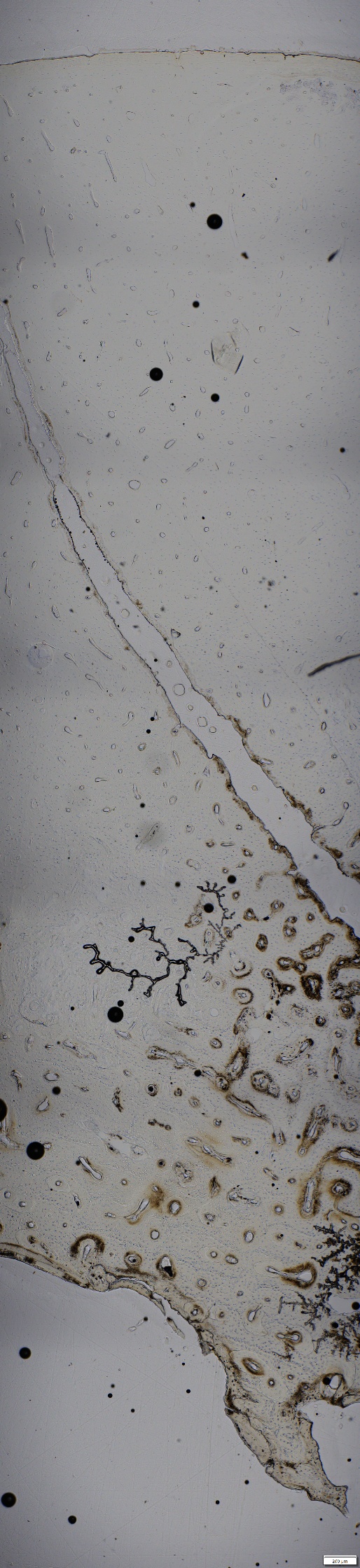

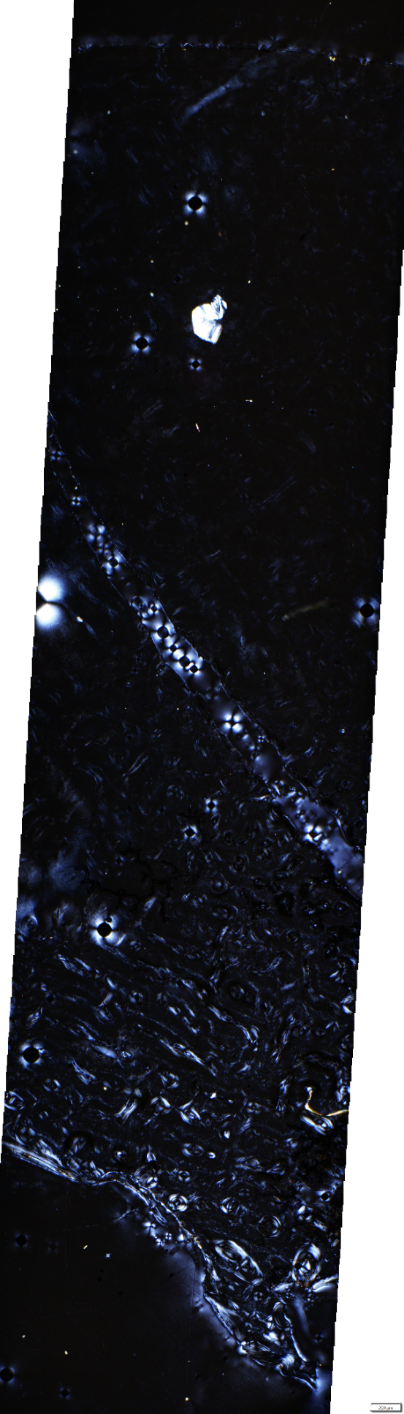

**32_11**

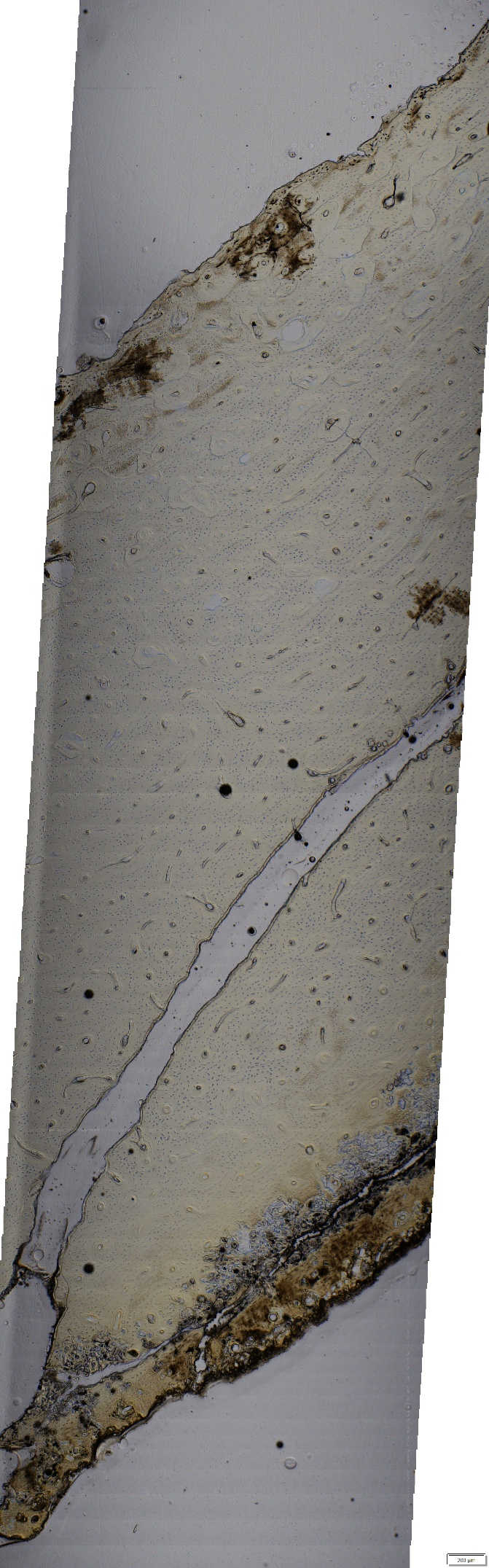

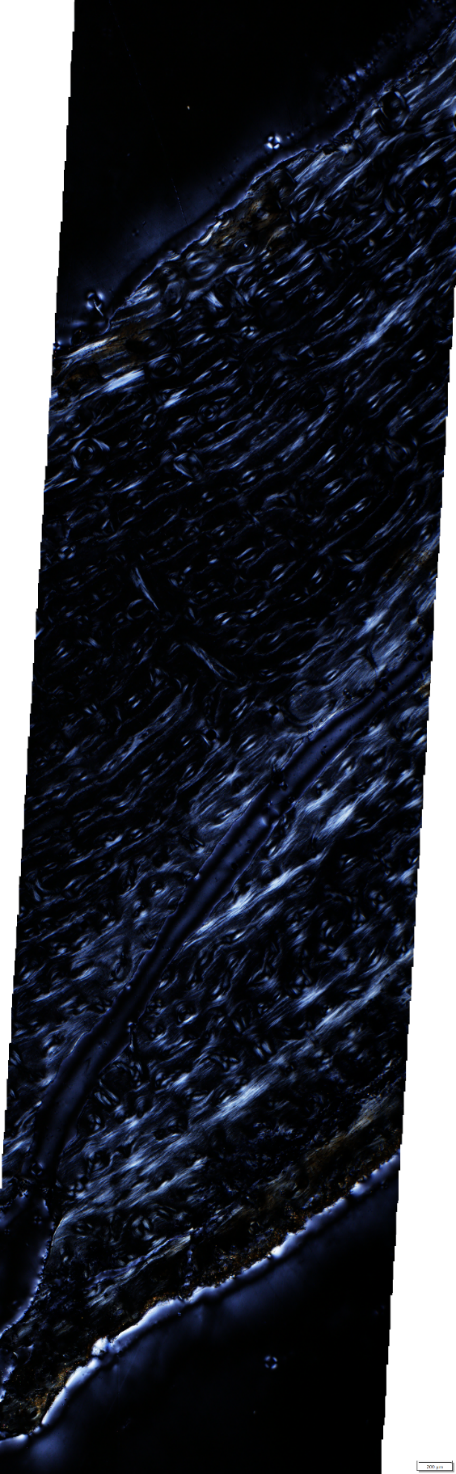

**32_6**

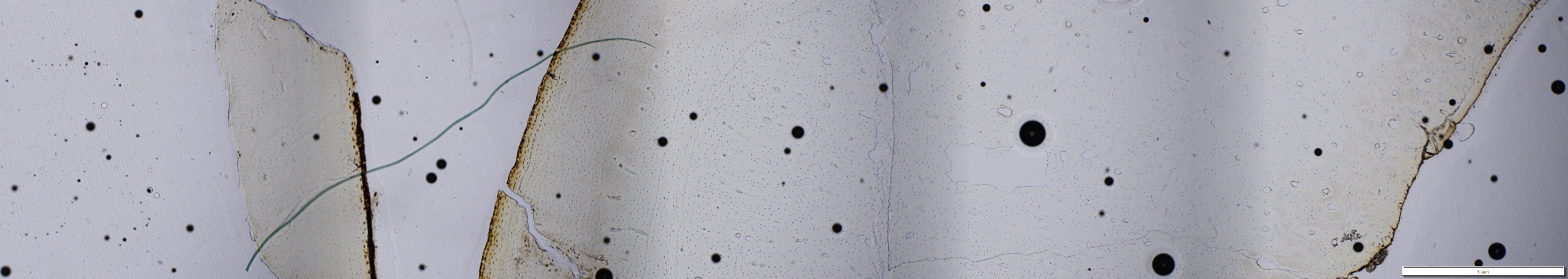

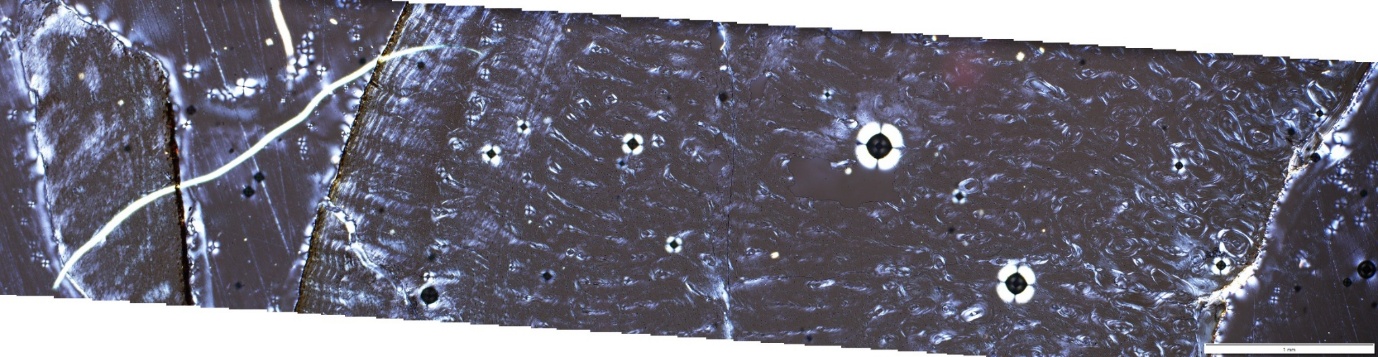

**12.0.1_7**

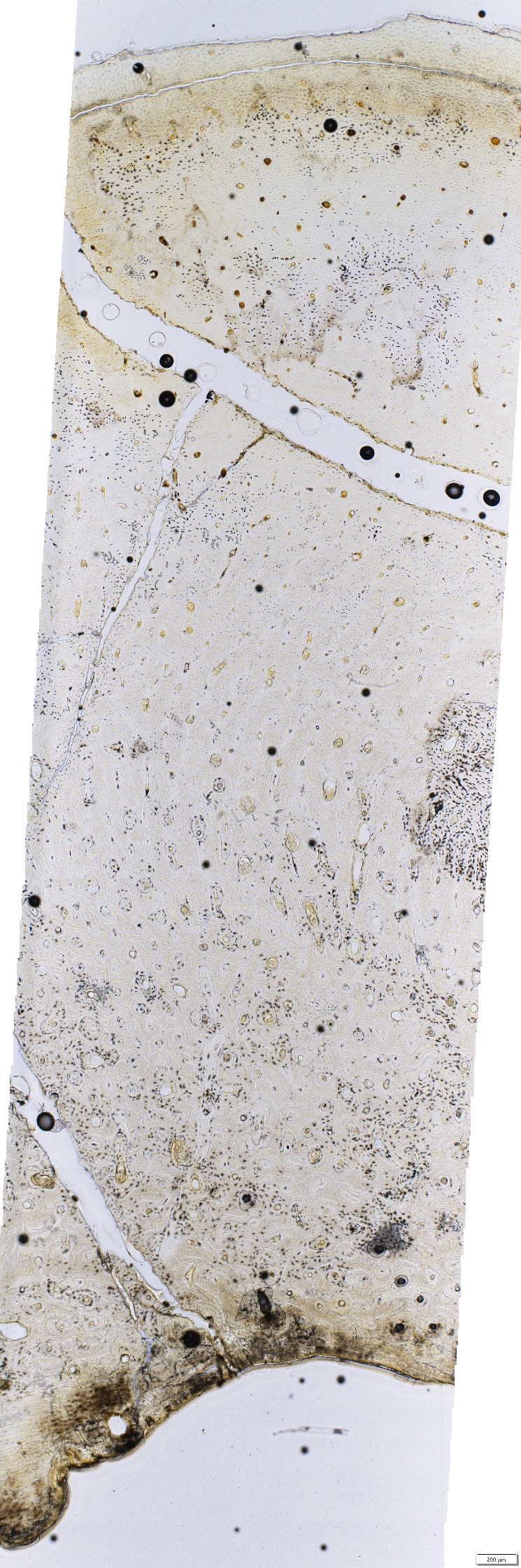

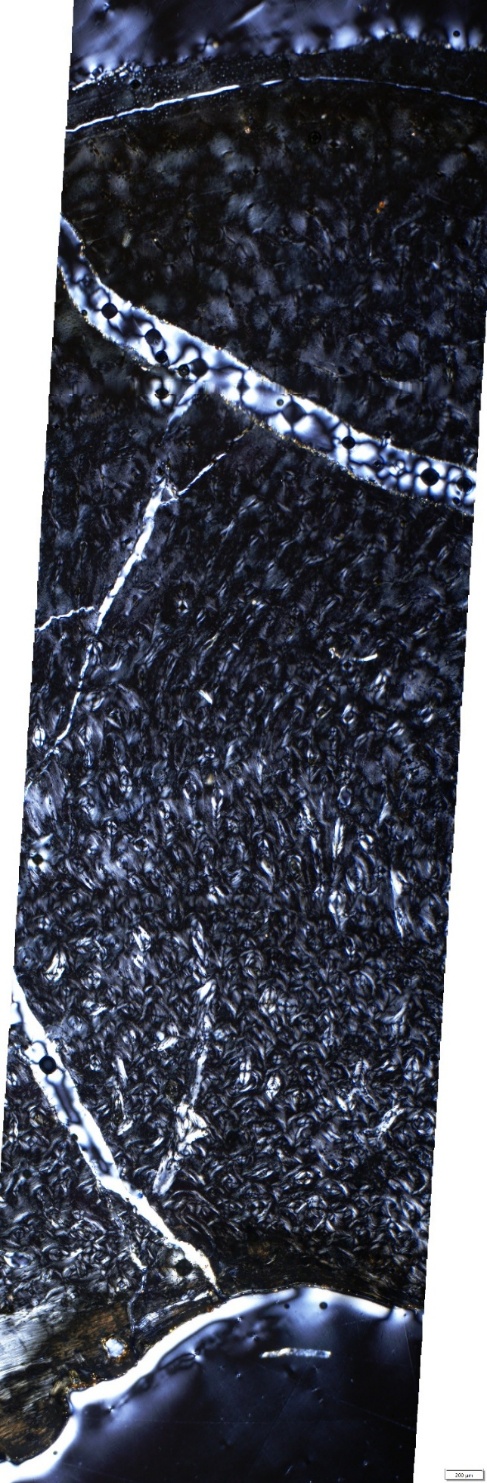
