## Supplementary 3 for "Underwater caves preserve biochemical composition and histological structure in modern and fossil mammalian bones"

**Supplementary 3: Descriptive and non-parametric statistical comparisons of r-FTIRM and s-FTIRM data between environmental conditions, bone regions, and tissue types for: *Underwater caves preserve biochemical composition and histological structure in modern and fossil mammalian bones***

Walker, MM, Miszkiewicz, JJ, Rowe, JM, Matheson, CD, Vongsvivut, J, Sims, NA and Louys J

Quantitative assessment of the reflectance Fourier-transform infrared microspectroscopy (r-FTIRM) and synchrotron-source (s-)FTIR in macro attenuated total reflectance Fourier-transform infrared mode are here reported in full. Data are broken down into the following categories, Section 1) r-FTIRM analysis, Section 2) s-FTIRM analysis, and Section 3) principal component analysis (PCA) of s-FTIRM data between depositional environments and tissue types. Significant results are determined using a 95% confidence interval.

Sections 1 (r-FTIRM) and 2 (s-FTIRM) report descriptive statistics and non-parametric assessments of individual diagenetic indices between depositional environments (historic wet, historic dry, and geologic fossil), and bone regions (sub-periosteal, mid-cortical, sub-endosteal). Analyses are limited to our research questions:

1. “*Are there any microstructural and biochemical differences in diagenesis between bones from wet and dry cave conditions?*
2. *Are there any microstructural and biochemical diagenetic differences between different regions of wet bone, specifically the secondary osteon and surrounding interstitial bone?*
3. *… explore a palaeontological assemblage … and theorise that early diagenesis will help inform the fossils’ original depositional environment (wet or dry).”*

The following tables report descriptive statistics for r-FTIRM and s-FTIRM data:

Table 1 **Descriptive statistics for r-FTIRM data** between each environmental condition and bone region.	3

Table 9 **Descriptive statistics of averaged region of interests (ROI)** between assemblage context, and interstitial (In.Ar) and osteonal (On.Ar) bone type sub-assemblages.	10

The following tables report statistical analyses that question:

- Are there bone region diagenetic differences between the median and distributions of all depositional environment groups?

Table 2 Statistical **comparison of diagenetic indices medians and distributions (Kruskal Wallis h test) from 1) historical dry, 2) historical wet, and 3) geologic fossil depositional environments**, from bone regions.	5

Table 10 Statistical analysis of independent distributions and medians of full **region of interests (ROI) compared between environmental conditions**.	12

- Are there bone region diagenetic differences between the median and distributions of wet and dry, wet and fossil, and dry and fossil depositional environment groups?

Table 3 **Comparison of wet and dry depositional environments** between different regions of the bone.	5

Table 4 **Comparison of wet and fossil depositional environments** between the bone regions.	6

Table 5 **Comparison of dry and fossil depositional environments** between bone regions:	7

- Are there diagenetic differences between the median and distributions of bone tissue regions within each depositional environment groups?

Table 6 Statistical comparison of diagenetic indices medians and distributions (Kruskal Wallis h test) from **1) sub-periosteal, 2) mid-cortical, and 3) sub-endosteal regions of the bone**, in different environments.	7

Table 7 Comparison of diagenetic indices **within wet bone specimens** (N=8) regions: lower (sub-endosteal), middle (mid-cortical), and upper (sub-periosteal).	8

Table 8 Comparison of diagenetic indices **within fossil bone specimens** (N=9) regions: lower (sub-endosteal), middle (mid-cortical), and upper (sub-periosteal).	8

Table 11 Comparison of **interstitial (In.Ar) and osteonal (On.Ar) bone differences in each environment**.	12

Section 3 reports PCA analyses that compare depositional environments and bone tissues based the diagenesis indices calculated from s-FTIRM spectral data. For each comparison, three PCAs were run based on a different suit of variables: 1) all mineral and organic diagenetic variables, 2) all mineral indices and AmI/P, and 3) only mineral indices. Each PCA is supported by model of fitness data, including: the total variance explained, component matrix, and PCA scatter plots. Model of fitness data are summarised in Section 3.1:

Table 12 Principal component analysis model fitness	14

Statistical significances of the principal components identified by the models are summarised in Section 3.1.1 and Section 3.1.2. Here, Mann Whitney U and median nonparametric tests compared principal components (PC) between the depositional environment or bone tissue type variables assessed by the model. The following tables report the model of fitness and statistical power of PCA analyses:

Table 13 Summary table of significance **comparing environments between whole region of interest (ROI)** based on principal components derived from diagenetic indices	14

Table 14 Summary Table of significance **between In and On bone type for each environment** based on Principal components derived from diagenetic indices	16

The following PCA tests were conducted (Section 3.2 and Section 3.3):

1. Comparison of ROI data for all environments (Section 3.2.1)
2. Comparison of ROI data for wet and dry environments (Section 3.2.2)
3. Comparison of In.Ar and On.Ar data for each depositional environment (fossil, dry and wet). (Section 3.3)

### 1 r-FTIRM: statistical comparison of group distribution and means

The following tables summarise descriptive statistics of each diagenesis index (Table 1), compare diagenesis indices between each depositional environment (Tables 2-5), and between bone regions within the same depositional environment (Tables 6-8). Statistical comparisons of medians and distributions are based on the mean average values from the sub-periosteal (upper), mid-cortical (middle) and sub-endosteal (lower) 600-700 microns of each strip of bones collected by r-FTIRM analysis.

#### 1.1 Descriptive statistics

Table 1 Descriptive statistics for r-FTIRM data between each environmental condition and bone region.

|  |  |  | **Sub-endosteal** | | | | | **Mid-cortical** | | | | | **Sub-periosteal** | | | | |
| --- | --- | --- | --- | --- | --- | --- | --- | --- | --- | --- | --- | --- | --- | --- | --- | --- | --- |
| **Index** | **Condition** | **N** | **x̄** | **x͂** | **σ** | **min** | **max** | **x̄** | **x͂** | **σ** | **min** | **max** | **x̄** | **x͂** | **σ** | **min** | **max** |
| CI1 | Dry | 6 | 1.291 | 1.289 | 0.030 | 1.248 | 1.342 | 1.232 | 1.218 | 0.042 | 1.179 | 1.299 | 1.331 | 1.324 | 0.060 | 1.263 | 1.425 |
|  | Wet | 8 | 1.295 | 1.270 | 0.102 | 1.181 | 1.501 | 1.257 | 1.254 | 0.042 | 1.185 | 1.306 | 1.343 | 1.325 | 0.069 | 1.270 | 1.463 |
|  | Fossil | 10 | 1.741 | 1.668 | 0.335 | 1.244 | 2.388 | 1.764 | 1.629 | 0.448 | 1.197 | 2.942 | 1.641 | 1.528 | 0.303 | 1.382 | 2.424 |
| CI2 | Dry | 6 | 1.175 | 1.170 | 0.041 | 1.129 | 1.246 | 1.175 | 1.175 | 0.018 | 1.151 | 1.210 | 1.147 | 1.151 | 0.027 | 1.095 | 1.176 |
|  | Wet | 8 | 1.162 | 1.156 | 0.033 | 1.117 | 1.208 | 1.167 | 1.176 | 0.021 | 1.136 | 1.192 | 1.156 | 1.152 | 0.035 | 1.110 | 1.231 |
|  | Fossil | 10 | 1.220 | 1.213 | 0.033 | 1.185 | 1.288 | 1.232 | 1.258 | 0.045 | 1.134 | 1.283 | 1.229 | 1.222 | 0.035 | 1.173 | 1.281 |
| C/P | Dry | 6 | 0.204 | 0.212 | 0.025 | 0.163 | 0.227 | 0.226 | 0.227 | 0.007 | 0.218 | 0.238 | 0.236 | 0.237 | 0.012 | 0.214 | 0.249 |
|  | Wet | 8 | 0.201 | 0.203 | 0.017 | 0.173 | 0.222 | 0.216 | 0.211 | 0.025 | 0.189 | 0.258 | 0.212 | 0.216 | 0.023 | 0.163 | 0.237 |
|  | Fossil | 10 | 0.187 | 0.194 | 0.019 | 0.160 | 0.217 | 0.186 | 0.190 | 0.015 | 0.155 | 0.200 | 0.204 | 0.209 | 0.020 | 0.176 | 0.243 |
| HPO_4_/P | Dry | 6 | 0.612 | 0.603 | 0.066 | 0.513 | 0.688 | 0.606 | 0.615 | 0.049 | 0.527 | 0.656 | 0.665 | 0.647 | 0.062 | 0.587 | 0.772 |
|  | Wet | 8 | 0.629 | 0.622 | 0.079 | 0.510 | 0.773 | 0.602 | 0.581 | 0.064 | 0.525 | 0.694 | 0.624 | 0.634 | 0.080 | 0.476 | 0.728 |
|  | Fossil | 10 | 0.571 | 0.576 | 0.047 | 0.498 | 0.648 | 0.521 | 0.486 | 0.064 | 0.452 | 0.671 | 0.522 | 0.524 | 0.055 | 0.421 | 0.604 |
| AmI/P | Dry | 6 | 0.230 | 0.272 | 0.131 | 0.049 | 0.367 | 0.374 | 0.370 | 0.053 | 0.292 | 0.454 | 0.228 | 0.204 | 0.079 | 0.166 | 0.384 |
|  | Wet | 8 | 0.345 | 0.370 | 0.104 | 0.152 | 0.481 | 0.403 | 0.395 | 0.062 | 0.308 | 0.497 | 0.317 | 0.321 | 0.067 | 0.233 | 0.429 |
|  | Fossil | 10 | 0.060 | 0.028 | 0.060 | 0.022 | 0.221 | 0.058 | 0.027 | 0.087 | 0.022 | 0.305 | 0.057 | 0.046 | 0.046 | 0.015 | 0.183 |
| AmII/P | Dry | 6 | 0.125 | 0.141 | 0.047 | 0.055 | 0.175 | 0.165 | 0.162 | 0.016 | 0.145 | 0.187 | 0.135 | 0.129 | 0.020 | 0.117 | 0.173 |
|  | Wet | 8 | 0.167 | 0.176 | 0.029 | 0.110 | 0.194 | 0.183 | 0.182 | 0.023 | 0.142 | 0.210 | 0.169 | 0.173 | 0.022 | 0.134 | 0.202 |
|  | Fossil | 10 | 0.087 | 0.062 | 0.065 | 0.044 | 0.258 | 0.058 | 0.046 | 0.032 | 0.038 | 0.147 | 0.486 | 0.050 | 0.255 | 0.033 | 1.279 |
| AmI/ AmII | Dry | 6 | 1.640 | 1.835 | 0.526 | 0.901 | 2.089 | 2.258 | 2.333 | 0.183 | 2.014 | 2.427 | 1.625 | 1.536 | 0.327 | 1.346 | 2.216 |
|  | Wet | 8 | 2.012 | 2.087 | 0.354 | 1.397 | 2.477 | 2.194 | 2.176 | 0.148 | 1.989 | 2.419 | 1.876 | 1.864 | 0.295 | 1.478 | 2.472 |
|  | Fossil | 10 | 0.742 | 0.571 | 0.406 | 0.356 | 1.729 | 0.747 | 0.596 | 0.477 | 0.451 | 2.081 | 0.916 | 0.932 | 0.300 | 0.446 | 1.591 |
| Collagen Integrity | Dry | 6 | 0.431 | 0.422 | 0.070 | 0.365 | 0.551 | 0.359 | 0.361 | 0.012 | 0.335 | 0.369 | 0.456 | 0.477 | 0.061 | 0.359 | 0.517 |
|  | Wet | 8 | 0.365 | 0.342 | 0.088 | 0.269 | 0.551 | 0.357 | 0.358 | 0.013 | 0.333 | 0.377 | 0.373 | 0.371 | 0.047 | 0.304 | 0.437 |
|  | Fossil | 10 | 0.539 | 0.550 | 0.069 | 0.443 | 0.657 | 0.601 | 0.615 | 0.093 | 0.364 | 0.708 | 0.554 | 0.542 | 0.072 | 0.458 | 0.669 |
| Random Coils (α) | Dry | 6 | 0.922 | 0.886 | 0.080 | 0.864 | 1.072 | 0.858 | 0.844 | 0.031 | 0.833 | 0.904 | 0.934 | 0.936 | 0.040 | 0.873 | 0.994 |
|  | Wet | 8 | 0.892 | 0.869 | 0.057 | 0.846 | 1.003 | 0.867 | 0.855 | 0.029 | 0.838 | 0.926 | 0.905 | 0.902 | 0.026 | 0.872 | 0.951 |
|  | Fossil | 10 | 1.118 | 1.140 | 0.086 | 0.928 | 1.224 | 1.110 | 1.129 | 0.087 | 0.873 | 1.191 | 1.090 | 1.069 | 0.084 | 0.944 | 1.279 |

#### 1.2 Non-parametric statistical comparison of bone regions between environments

Table 2 Statistical **comparison of diagenetic indices medians and distributions (Kruskal-Wallis h test) from 1) historical dry, 2) historical wet, and 3) geologic fossil depositional environments**, between different sections of the bone. N=23. Red data are not considered reliable due to the heavily degraded amide spectra. Significant values are bolded and annotated by an asterisk*.

|  |  | **Sub-endosteal** | | **Mid-cortical** | | **Sub-periosteal** | | |
| --- | --- | --- | --- | --- | --- | --- | --- | --- |
|  | **Diagenesis Index** | **Test statistic** | ***p-*value** | **Test statistic** | ***p-*value** | | **Test statistic** | ***p-*value** |
| C/P | *Distribution* | 3.470 | 0.176 | 13.166 | **0.001*** | | 7.580 | **0.023*** |
|  | *Median* | 1.626 | 0.443 | 15.486 | **<0.001*** | | 5..411 | 0.067 |
| HPO_4_/P | *Distribution* | 3.168 | 0.205 | 7.025 | **0.030*** | | 11.212 | **0.004*** |
|  | *Median* | 1.459 | 0.482 | 8.584 | **0.014*** | | 8.584 | **0.014*** |
| CI1 | *Distribution* | 10.471 | **0.005*** | 11.109 | **0.004*** | | 13.458 | **0.001*** |
|  | *Median* | 10.087 | **0.006*** | 10.087 | **0.006*** | | 10.087 | **0.006*** |
| CI2 | *Distribution* | 7.599 | **0.022*** | 8.444 | **0.015*** | | 12.935 | **0.002*** |
|  | *Median* | 5.411 | 0.067 | 10.087 | **0.006*** | | 16.154 | **<0.001*** |
| AmI/P | *Distribution* | 14.944 | <0.001 | 15.720 | <0.001 | | 17.212 | <0.001 |
|  | *Median* | 14.150 | <0.001 | 13.649 | 0.001 | | 16.989 | <0.001 |
| AmII/P | *Distribution* | 9.054 | 0.11 | 15.701 | <0.001 | | 18.039 | <0.001 |
|  | *Median* | 9.920 | 0.007 | 14.150 | <0.001 | | 16.989 | <0.001 |
| AmI/ AmII | *Distribution* | 14.213 | <0.001 | 14.099 | <0.001 | | 14.749 | <0.001 |
|  | *Median* | 9.920 | 0.007 | 8.083 | 0.018 | | 9.920 | 0.007 |
| Collagen Integrity | *Distribution* | 12.343 | 0.002 | 13.911 | <0.001 | | 14.687 | <0.001 |
|  | *Median* | 10.588 | 0.005 | 10.087 | 0.006 | | 13.426 | 0.001 |
| Random Coils (α) | *Distribution* | 14.097 | <0.001 | 14.295 | <0.001 | | 15.311 | <0.001 |
|  | *Median* | 10.087 | 0.006 | 10.087 | 0.006 | | 10.588 | 0.005 |

Table 3 **Comparison of wet and dry depositional environments** between different regions of the bone: wet (N=8), dry (N=6), unknown (N=9). Significant values are bolded and annotated by an asterisk*.

| **Wet Vs Dry (Historical)** | | **Sub-endosteal** | | **Mid-cortical** | | **Sub-periosteal** | | |
| --- | --- | --- | --- | --- | --- | --- | --- | --- |
|  | **Diagenesis Index** | **Test statistic** | ***p-*value** | **Test statistic** | ***p-*value** | | **Test statistic** | ***p-*value** |
| C/P | *Distribution* | 19.000 | 0.573 | 12.000 | 0.142 | | 6.000 | **0.020*** |
|  | *Median* | 1.167 | 0.592 | 4.667 | 0.103 | | 4.667 | 0.103 |
| HPO_4_/P | *Distribution* | 26.000 | 0.852 | 22.000 | 0.852 | | 19.000 | 0.573 |
|  | *Median* | 0.000 | 1.000 | 1.167 | 0.592 | | 0.000 | 1.000 |
| CI1 | *Distribution* | 20.000 | 0.662 | 35.000 | 0.181 | | 27.000 | 0.755 |
|  | *Median* | 0.000 | 1.000 | 1.167 | 0.592 | | 0.000 | 1.000 |
| CI2 | *Distribution* | 28.000 | 0.662 | 27.000 | 0.755 | | 23.000 | 0.950 |
|  | *Median* | 1.167 | 0.592 | 0.000 | 1.000 | | 0.000 | 1.000 |
| AmI/P | *Distribution* | 38.000 | 0.081 | 32.000 | 0.345 | | 41.000 | 0.029 |
|  | *Median* | 1.167 | 0.592 | 1.167 | 0.592 | | 4.667 | 0.103 |
| AmII/P | *Distribution* | 39.000 | 0.059 | 36.000 | 0.142 | | 43.000 | 0.013 |
|  | *Median* | 4.667 | 0.103 | 1.167 | 0.592 | | 4.667 | 0.103 |
| AmI/AmII | *Distribution* | 36.000 | 0.142 | 17.000 | 0.414 | | 37.000 | 0.108 |
|  | *Median* | 1.167 | 0.592 | 1.167 | 0.592 | | 4.667 | 0.103 |
| Collagen Integrity | *Distribution* | 8.000 | **0.043*** | 19.000 | 0.573 | | 8.000 | 0.043 |
|  | *Median* | 4.667 | 0.103 | 1.167 | 0.592 | | 1.167 | 0.592 |
| Random Coils (α) | *Distribution* | 14.000 | 0.228 | 33.000 | 0.282 | | 12.000 | 0.142 |
|  | *Median* | 1.167 | 0.592 | 1.167 | 0.592 | | 1.167 | 0.592 |

Table 4 **Comparison of wet and fossil depositional environments** between the bone regions: wet (N=8), fossil (N=9). Red data are not considered reliable due to the heavily degraded amide spectra. Significant values are bolded and annotated by an asterisk*.

| **Wet V Fossil** | | **Sub-endosteal** | | **Mid-cortical** | | **Sub-periosteal** | | |
| --- | --- | --- | --- | --- | --- | --- | --- | --- |
|  | **Diagenesis Index** | **Test statistic** | ***p-*value** | **Test statistic** | ***p-*value** | | **Test statistic** | ***p-*value** |
| C/P | *Distribution* | 53.000 | 0.114 | 63.000 | **0.008*** | | 46.000 | 0.370 |
|  | *Median* | 1.446 | 0.347 | 4.735 | 0.057 | | 0.052 | 1.000 |
| HPO_4_/P | *Distribution* | 53.000 | 0.114 | 60.000 | **0.021*** | | 62.000 | **0.011*** |
|  | *Median* | 1.446 | 0.347 | 4.735 | 0.057 | | 4.735 | 0.57 |
| CI1 | *Distribution* | 6.000 | **0.002*** | 7.000 | **0.004*** | | 4.000 | **0.001*** |
|  | *Median* | 7.244 | **0.015*** | 13.432 | **0.000*** | | 7.244 | **0.015*** |
| CI2 | *Distribution* | 9.000 | **0.008*** | 10.000 | **0.011*** | | 5.000 | **0.002*** |
|  | *Median* | 2.951 | 0.153 | 7.244 | **0.015*** | | 7.244 | **0.015*** |
| AmI/P | *Distribution* | 71.000 | 0.000 | 72.000 | 0.000 | | 72.000 | 0.000 |
|  | *Median* | 9.920 | 0.003 | 17.000 | 0.000 | | 17.000 | 0.000 |
| AmII/P | *Distribution* | 63.000 | 0.008 | 71.000 | 0.000 | | 72.000 | 0.000 |
|  | *Median* | 9.920 | 0.003 | 9.920 | 0.003 | | 17.000 | 0.000 |
| AmI/ AmII | *Distribution* | 71.000 | 0.000 | 70.000 | 0.000 | | 71.000 | 0.000 |
|  | *Median* | 9.920 | 0.003 | 9.920 | 0.003 | | 9.920 | 0.003 |
| Collagen Integrity | *Distribution* | 5.000 | 0.002 | 2.000 | 0.000 | | 0.000 | 0.000 |
|  | *Median* | 7.244 | 0.015 | 13.432 | 0.000 | | 13.432 | 0.000 |
| Random Coils (α) | *Distribution* | 2.000 | 0.000 | 2.000 | 0.000 | | 1.000 | 0.000 |
|  | *Median* | 13.432 | 0.000 | 13.432 | 0.000 | | 13.432 | 0.000 |

Table 5 **Comparison of dry and fossil depositional environments** between bone regions: dry (N=6), fossil (N=9). Red data are not considered reliable due to the heavily degraded amide spectra. Significant values are bolded and annotated by an asterisk*.

| **Dry Vs Fossil** | | **Sub-endosteal** | | **Mid-cortical** | | **Sub-periosteal** | | |
| --- | --- | --- | --- | --- | --- | --- | --- | --- |
|  | **Diagenesis Index** | **Test statistic** | ***p-*value** | **Test statistic** | ***p-*value** | | **Test statistic** | ***p-*value** |
| C/P | *Distribution* | 15.000 | 0.181 | 0.000 | **0.000*** | | 7.000 | **0.018*** |
|  | *Median* | 1. 607 | 0.315 | 11.429 | **0.001*** | | 5.402 | **0.041*** |
| HPO_4_/P | *Distribution* | 16.000 | 0.224 | 9.000 | **0.036*** | | 1.000 | **0.001*** |
|  | *Median* | 1.607 | 0.315 | 5.402 | **0.041*** | | 11.429 | **0.001*** |
| CI1 | *Distribution* | 48.000 | **0.012*** | 49.000 | **0.008*** | | 53.000 | **0.001*** |
|  | *Median* | 8.750 | **0.007*** | 8.750 | **0.007*** | | 8.750 | **0.007*** |
| CI2 | *Distribution* | 43.000 | 0.066 | 47.000 | **0.018*** | | 53.000 | **0.001*** |
|  | *Median* | 0.714 | 0.608 | 8.750 | **0.007*** | | 8.750 | **0.007*** |
| AmI/P | *Distribution* | 4.000 | 0.005 | 1.000 | 0.001 | | 1.000 | 0.001 |
|  | *Median* | 5.402 | 0.041 | 11.429 | 0.001 | | 11.429 | 0.001 |
| AmII/P | *Distribution* | 12.000 | 0.088 | 1.000 | 0.001 | | 0.000 | 0.000 |
|  | *Median* | 1.607 | 0.315 | 11.429 | 0.001 | | 11.429 | 0.001 |
| AmI/ AmII | *Distribution* | 5.000 | 0.008 | 2.000 | 0.002 | | 4.000 | 0.005 |
|  | *Median* | 5.402 | 0.041 | 11.429 | 0.001 | | 11.429 | 0.001 |
| Collagen Integrity | *Distribution* | 47.000 | 0.018 | 52.000 | 0.002 | | 46.000 | 0.026 |
|  | *Median* | 3.616 | 0.119 | 8.750 | 0.007 | | 3.616 | 0.119 |
| Random Coils (α) | *Distribution* | 51.000 | 0.003 | 52.000 | 0.002 | | 52.000 | 0.002 |
|  | *Median* | 8.750 | 0.007 | 8.750 | 0.007 | | 8.750 | 0.007 |

#### 1.3 r-FTIRM: comparison of bone regions within each environment

Table 6 Statistical comparison of diagenetic indices medians and distributions (Kruskal Wallis h test) from 1) sub-periosteal, 2) mid-cortical, and 3) sub-endosteal regions of the bone, in different environments. Red data are not considered reliable due to the heavily degraded amide spectra. Significant values are bolded and annotated by an asterisk*.

|  |  | **Dry (Historical)** | | | **Wet (Historical)** | | | **Fossil (Geological)** | |
| --- | --- | --- | --- | --- | --- | --- | --- | --- | --- |
| **Diagenesis Index** | | **Test statistic** | ***p*** | **Test statistic** | | ***p*** | **Test statistic** | | ***p*** |
| C/P | *Distribution* | 8.667 | **0.013*** | 1.545 | | 0.462 | 4.025 | | 0.134 |
|  | *Median* | 9.333 | **0.009*** | 4.000 | | 0.135 | 2.077 | | 0.354 |
| HPO_4_/P | *Distribution* | 2.257 | 0.323 | 0.740 | | 0.691 | 4.998 | | 0.082 |
|  | *Median* | 4.000 | 0.135 | 1.000 | | 0.607 | 2.077 | | 0.354 |
| Crystallinity 1 | *Distribution* | 7.684 | **0.021*** | 5.145 | | 0.076 | 1.168 | | 0.588 |
|  | *Median* | 5.333 | 0.069 | 4.000 | | 0.135 | 0.297 | | 0.862 |
| Crystallinity 2 | *Distribution* | 2.889 | 0.236 | 1.085 | | 0.581 | 1.083 | | 0.582 |
|  | *Median* | 1.333 | 0.513 | 1.000 | | 0.607 | 2.077 | | 0.354 |
| AmI/P | *Distribution* | 6.924 | **0.031*** | 4.515 | | 0.105 | 2.501 | | 0.286 |
|  | *Median* | 5.333 | 0.069 | 4.000 | | 0.135 | 5.637 | | 0.060 |
| AmII/P | *Distribution* | 5.626 | 0.060 | 1.295 | | 0.523 | 2.000 | | 0.386 |
|  | *Median* | 5.333 | 0.069 | 1.000 | | 0.607 | 0.297 | | 0.862 |
| AmI/AmII | *Distribution* | 7.450 | 0.024 | 5.285 | | 0.071 | 3.644 | | 0.162 |
|  | *Median* | 5.333 | 0.069 | 7.000 | | **0.030*** | 4.747 | | 0.093 |
| Collagen integrity | *Distribution* | 7.614 | **0.022*** | 1.055 | | 0.590 | 3.820 | | 0.148 |
|  | *Median* | 9.333 | **0.009*** | 1.000 | | 0.607 | 9.198 | | **0.010*** |
| Random Coils (α) | *Distribution* | 6.678 | **0.035*** | 5.505 | | 0.064 | 2.384 | | 0.304 |
|  | *Median* | 4.000 | 0.135 | 7.000 | | **0.030*** | 3.857 | | 0.145 |

Table 7 Comparison of diagenetic indices **within wet bone specimens** (N=8) regions: lower (sub-endosteal), middle (mid-cortical), and upper (sub-periosteal). Significant values are bolded and annotated by an asterisk*.

|  |  | **Lower V Upper** | | | **Middle V Lower** | | | **Middle V Upper** | |
| --- | --- | --- | --- | --- | --- | --- | --- | --- | --- |
| **Diagenesis Index** | | **Test statistic** | ***p*** | **Test statistic** | | ***p*** | **Test statistic** | | ***p*** |
| C/P | *Distribution* | 43.000 | 0.279 | 41.000 | | 0.382 | 34.000 | | 0.878 |
|  | *Median* | 4.000 | 0.132 | 1.000 | | 0.619 | 0.000 | | 1.000 |
| HPO_4_/P | *Distribution* | 32.000 | 1.000 | 26.000 | | 0.574 | 40.000 | | 0.442 |
|  | *Median* | 0.000 | 1.000 | 1.000 | | 0.619 | 1.000 | | 0.619 |
| Crystallinity 1 | *Distribution* | 45.000 | 0.195 | 26.000 | | 0.574 | 54.000 | | **0.021*** |
|  | *Median* | 1.000 | 0.619 | 0.000 | | 1.000 | 4.000 | | 0.132 |
| Crystallinity 2 | *Distribution* | 36.000 | 0.721 | 31.000 | | 0.959 | 44.000 | | 0.234 |
|  | *Median* | 0.000 | 1.000 | 1.000 | | 0.619 | 1.000 | | 0.619 |
| AmI/P | *Distribution* | 24.000 | 0.442 | 42.000 | | 0.328 | 11.000 | | 0.028 |
|  | *Median* | 1.000 | 0.619 | 1.000 | | 0.619 | 4.000 | | 0.132 |
| AmII/P | *Distribution* | 29.000 | 0.798 | 40.000 | | 0.442 | 22.000 | | 0.328 |
|  | *Median* | 1.000 | 0.619 | 0.000 | | 1.000 | 1.000 | | 0.619 |
| AmI/AmII | *Distribution* | 24.000 | 0.442 | 41.000 | | 0.328 | 8.000 | | **0.010*** |
|  | *Median* | 1.000 | 0.619 | 1.000 | | 0.619 | 9.000 | | **0.010*** |
| Collagen integrity | *Distribution* | 39.000 | 0.505 | 39.000 | | 0.505 | 40.000 | | 0.442 |
|  | *Median* | 1.000 | 0.619 | 1.000 | | 0.619 | 1.000 | | 0.619 |
| Random Coils (α) | *Distribution* | 46.000 | 0.161 | 22.000 | | 0.328 | 53.000 | | 0.028 |
|  | *Median* | 4.000 | 0.132 | 1.000 | | 0.619 | 4.000 | | 0.132 |

Table 8 Comparison of diagenetic indices **within fossil bone specimens** (N=9) regions: lower (sub-endosteal), middle (mid-cortical), and upper (sub-periosteal). Red data are not considered reliable due to the heavily degraded amide spectra. Significant values are bolded and annotated by an asterisk*.

|  |  | **Lower V Upper** | | | **Middle V Lower** | | | **Middle V Upper** | |
| --- | --- | --- | --- | --- | --- | --- | --- | --- | --- |
| **Diagenesis Index** | | **Test statistic** | ***p-*value** | **Test statistic** | | ***p-*value** | **Test statistic** | | ***p-*value** |
| C/P | *Distribution* | 60.000 | 0.094 | 40.000 | | 1.000 | 60.000 | | 0.094 |
|  | *Median* | 2.000 | 0.347 | 0.222 | | 1.000 | 0.222 | | 1.000 |
| HPO_4_/P | *Distribution* | 20.000 | 0.077 | 18.000 | | 0.050 | 45.000 | | 0.730 |
|  | *Median* | 2.000 | 0.347 | 2.000 | | 0.347 | 0.222 | | 1.000 |
| Crystallinity 1 | *Distribution* | 30.000 | 0.387 | 40.000 | | 1.000 | 30.000 | | 0.387 |
|  | *Median* | 0.222 | 1.000 | 0.222 | | 1.000 | 0.222 | | 1.000 |
| Crystallinity 2 | *Distribution* | 47.000 | 0.605 | 51.000 | | 0.387 | 33.000 | | 0.546 |
|  | *Median* | 0.222 | 1.000 | 2.000 | | 0.347 | 0.222 | | 1.000 |
| AmI/P | *Distribution* | 44.000 | 0.796 | 34.000 | | 0.605 | 62.000 | | 0.063 |
|  | *Median* | 0.222 | 1.000 | 0.222 | | 1.000 | 5.556 | | 0.057 |
| AmII/P | *Distribution* | 29.000 | 0.340 | 25.000 | | 0.190 | 43.000 | | 0.863 |
|  | *Median* | 0.222 | 1.000 | 0.222 | | 1.000 | 0.222 | | 1.000 |
| AmI/AmII | *Distribution* | 56.000 | 0.190 | 46.000 | | 0.666 | 62.000 | | 0.063 |
|  | *Median* | 2.000 | 0.347 | 0.222 | | 1.000 | 5.556 | | 0.057 |
| Collagen integrity | *Distribution* | 45.000 | 0.730 | 60.000 | | 0.063 | 25.000 | | 0.190 |
|  | *Median* | 0.222 | 1.000 | 5.556 | | 0.057 | 2.000 | | 0.347 |
| Random Coils (α) | *Distribution* | 28.000 | 0.297 | 37.000 | | 0.796 | 23.000 | | 0.136 |
|  | *Median* | 2.000 | 0.347 | 0.222 | | 1.000 | 5.556 | | 0.057 |

### 2 s-FTIRM statistical analyses

The following data compare the mean differences in s-FTIRM data collected from a 120 x 120-micron region of interest (ROI), composed of interstitial bone (In.Ar) and osteonal bone (On.Ar). The impact of burial environmental conditions is compared using mean ROI data (Table 10), and tissue types in each burial environment compare differences between In.Ar and On.Ar (Table 11).

Principal component analyses (PCA) were performed to further distinguish similarities and differences between environments, and tissue types in Section 3.

#### 2.1 Descriptive statistics

Table 9 Descriptive statistics of averaged region of interests (ROI) between assemblage context, and interstitial (In.Ar) and osteonal (On.Ar) bone type sub-assemblages. Red text highlights data not considered reliable due to severe degradation.

|  |  |  | **ROI** | | | | **In.Ar** | | | | | | **On.Ar** | | | | |
| --- | --- | --- | --- | --- | --- | --- | --- | --- | --- | --- | --- | --- | --- | --- | --- | --- | --- |
| **Index** | **Condition** | **N** | **x̄** | **x͂** | **σ** | **Range** | **N** | **x̄** | **x͂** | **σ** | **Range** | **N** | | **x̄** | **x͂** | **σ** | **Range** |
| C/P | Dry | 11 | 0.26 | 0.27 | 0.03 | 0.22-0.30 | 11 | 0.26 | 0.27 | 0.04 | 0.17 – 0.32 | 11 | | 0.27 | 0.27 | 0.03 | 022 – 0.34 |
|  | Wet | 14 | 0.25 | 0.25 | 0.03 | 0.18 -0.31 | 14 | 0.23 | 0.24 | 0.04 | 0.14 – 0.28 | 14 | | 0.27 | 0.26 | 0.04 | 0.21 – 0.37 |
|  | Fossil | 14 | 0.22 | 0.22 | 0.03 | 0.18-0.29 | 13 | 0.22 | 0.21 | 0.03 | 0.18 – 0.28 | 13 | | 0.19 | 0.20 | 0.04 | 0.11 – 0.27 |
| HPO4/P | Dry | 11 | 0.23 | 0.23 | 0.04 | 0.14-0.29 | 11 | 0.24 | 0.27 | 0.06 | 0.10 – 0.31 | 11 | | 0.23 | 0.21 | 0.06 | 0.16 – 0.32 |
|  | Wet | 14 | 0.23 | 0.23 | 0.03 | 0.17-0.27 | 14 | 0.20 | 0.20 | 0.07 | 0.11 – 0.38 | 14 | | 0.26 | 0.26 | 0.04 | 0.18 – 0.32 |
|  | Fossil | 14 | 0.18 | 0.18 | 0.04 | 0.07-0.21 | 13 | 0.18 | 0.18 | 0.06 | 0.06 – 0.32 | 13 | | 0.17 | 0.15 | 0.07 | 0.07 – 0.33 |
| CI1 | Dry | 11 | 0.94 | 0.94 | 0.02 | 0.89-0.97 | 11 | 0.95 | 0.95 | 0.04 | 0.86 – 1.00 | 11 | | 0.93 | 0.93 | 0.03 | 0.90 – 0.98 |
|  | Wet | 14 | 0.95 | 0.95 | 0.02 | 0.91-1.00 | 14 | 0.94 | 0.94 | 0.03 | 0.86 – 1.02 | 14 | | 0.97 | 0.97 | 0.03 | 0.91 – 1.03 |
|  | Fossil | 14 | 0.99 | 0.99 | 0.02 | 0.95-1.01 | 13 | 0.99 | 0.99 | 0.03 | 0.93 – 1.03 | 13 | | 0.99 | 0.99 | 0.04 | 0.91 – 1.04 |
| CI2 | Dry | 11 | 1.37 | 1.35 | 0.07 | 1.31-1.56 | 11 | 1.36 | 1.32 | 0.13 | 1.26 – 1.69 | 11 | | 1.39 | 1.38 | 0.13 | 1.28 – 1.75 |
|  | Wet | 14 | 1.38 | 1.37 | 0.06 | 1.31-1.53 | 14 | 1.41 | 1.40 | 0.11 | 1.24 – 1.64 | 14 | | 1.34 | 1.32 | 0.05 | 1.26 – 1.42 |
|  | Fossil | 14 | 1.55 | 1.51 | 0.15 | 1.45-1.97 | 13 | 1.51 | 1.50 | 0.12 | 1.36 – 1.86 | 13 | | 1.55 | 1.54 | 0.15 | 1.34 – 1.97 |
| AmI/P | Dry | 11 | 0.28 | 0.28 | 0.05 | 0.21-0.36 | 11 | 0.27 | 0.27 | 0.05 | 0.19 – 0.33 | 11 | | 0.30 | 0.29 | 0.05 | 0.24 – 0.4 |
|  | Wet | 14 | 0.29 | 0.29 | 0.06 | 0.17-0.41 | 14 | 0.25 | 0.27 | 0.07 | 0.09 – 0.34 | 14 | | 0.31 | 0.30 | 0.06 | 0.24 – 0.47 |
|  | Fossil | 14 | 0.03 | 0.02 | 0.01 | 0.02-0.06 | 13 | 0.03 | 0.02 | 0.02 | 0.01 – 0.08 | 13 | | 0.03 | 0.02 | 0.01 | 0.01 – 0.05 |
| AmII/P | Dry | 11 | 0.23 | 0.22 | 0.03 | 0.19-0.28 | 11 | 0.21 | 0.21 | 0.03 | 0.17 – 0.25 | 11 | | 0.24 | 0.23 | 0.04 | 0.20 – 0.30 |
|  | Wet | 14 | 0.23 | 0.24 | 0.04 | 0.14-0.31 | 14 | 0.22 | 0.24 | 0.06 | 0.08 – 0.30 | 14 | | 0.25 | 0.25 | 0.04 | 0.20 – 0.34 |
|  | Fossil | 14 | 0.06 | 0.05 | 0.01 | 0.05-0.09 | 13 | 0.06 | 0.05 | 0.02 | 0.02 – 0.11 | 13 | | 0.05 | 0.05 | 0.01 | 0.04 – 0.08 |
| AmI/ AmII | Dry | 11 | 1.48 | 1.48 | 0.05 | 1.14-1.59 | 11 | 1.50 | 1.47 | 0.13 | 1.36 – 1.86 | 11 | | 1.47 | 1.47 | 0.06 | 1.38 – 1.56 |
|  | Wet | 14 | 1.43 | 1.42 | 0.12 | 1.29-1.71 | 14 | 1.41 | 1.41 | 0.08 | 1.27 – 1.54 | 14 | | 1.40 | 1.41 | 0.24 | 0.24 – 1.13 |
|  | Fossil | 14 | 0.69 | 0.70 | 0.27 | 0.32-1.24 | 13 | 0.76 | 0.67 | 0.31 | 0.46 – 1.36 | 13 | | 0.60 | 0.60 | 0.12 | 1.24 – 1.61 |
| Collagen Integrity | Dry | 11 | 0.28 | 0.28 | 0.03 | 0.21-0.31 | 11 | 0.28 | 0.29 | 0.04 | 0.18 – 0.33 | 11 | | 0.28 | 0.28 | 0.03 | 0.23 – 0.31 |
|  | Wet | 14 | 0.28 | 0.29 | 0.04 | 0.18-0.33 | 14 | 0.28 | 0.29 | 0.05 | 0.17 – 0.33 | 14 | | 0.28 | 0.30 | 0.04 | 0.20 – 0.34 |
|  | Fossil | 13 | 0.37 | 0.20 | 0.46 | 0.13 – 1.80 | 12 | 0.39 | 0.18 | 0.53 | 0.10 – 2.00 | 10 | | 0.51 | 0.15 | 0.87 | 0.07 – 2.87 |
| Random Coils (α) | Dry | 11 | 0.95 | 0.95 | 0.03 | 0.90-1.01 | 11 | 0.94 | 0.94 | 0.06 | 0.82 – 1.04 | 11 | | 0.95 | 0.94 | 0.03 | 0.92 – 1.01 |
|  | Wet | 14 | 0.95 | 0.95 | 0.03 | 0.90-1.03 | 14 | 0.95 | 0.95 | 0.04 | 0.87 – 1.05 | 14 | | 0.96 | 0.96 | 0.03 | 0.90 – 1.00 |
|  | Fossil | 14 | 4.04 | 3.32 | 2.75 | 0.95-8.25 | 13 | 4.22 | 4.00 | 2.98 | 0.95 – 9.12 | 13 | | 4.33 | 4.45 | 3.10 | 0.97 – 11.93 |

#### 2.2 Statistical comparisons of group distributions and medians

Table 10 Statistical analysis of independent distributions and medians of full region of interests (ROI) compared between environmental conditions. Red data are not considered reliable due to the heavily degraded amide spectra. Significant values are bolded and annotated by an asterisk*.

|  |  | **ROI Between Environs** | | **Dry Vs Wet** | | **Dry Vs Fossil** | | **Wet Vs Fossil** | |
| --- | --- | --- | --- | --- | --- | --- | --- | --- | --- |
| **Diagenesis index** | | **Test statistic** | ***p-*value** | **Test statistic** | ***p-*value*** | **Test statistic** | ***p-*value*** | **Test statistic** | ***p-*value*** |
| C/P | *Distribution* | 11.393 | **0.003*** | 54.000 | 0.222 | 23.000 | **.002*** | 154.000 | **.009*** |
|  | *Median* | 10.540 | **0.005*** | 4.812 | **0.047*** | 4.812 | **.047*** | 5.143 | .057 |
| HPO_4_/P | *Distribution* | 16.888 | **<0.001*** | 72.000 | .809 | 12.000 | **.000*** | 173.000 | **.000*** |
|  | *Median* | 21.014 | **< 0.001*** | .051 | 1.000 | 14.490 | **.000*** | 9.143a | **.007*** |
| CI1 | *Distribution* | 19.377 | **<0.001*** | 99.000 | 0.244 | 149.000 | **.000*** | 24.000 | **.000*** |
|  | *Median* | 17.297 | **< 0.001*** | 0.051 | 1.000 | 18.132 | **.000*** | 14.286 | **.000*** |
| CI2 | *Distribution* | 19.530 | **<0.001*** | 92.00 | 0.434 | 142.000 | **.000*** | 13.000 | **.000*** |
|  | *Median* | 23.015 | **< 0.001*** | 1.066 | 0.428 | 1.464 | **.000*** | 14.286 | **.000*** |
| AmI/P | *Distribution* | 26.375 | **<0.001*** | 87.000 | 0.609 | .000 | .000 | 196.000 | .000 |
|  | *Median* | 20.832 | **<0.001*** | 3.381 | 0.111 | 21.280 | .000 | 28.000 | .000 |
| AmII/P | *Distribution* | 26.570 | **<0.001*** | 93.000 | .403 | .000 | .000 | 196.000 | .000 |
|  | *Median* | 21.950 | **< 0.001*** | 1.066 | .428 | 21.280 | .000 | 28.000 | .000 |
| AmI/AmII | *Distribution* | 27.868 | **<0.001*** | 41.000 | .051 | .000 | .000 | 196.000 | .000 |
|  | *Median* | 25.277 | **< 0.001*** | 4.812 | **.047*** | 21.280 | .000 | 28.000 | .000 |
| Collagen integrity | *Distribution* | 5.644 | 0.059 | 92.000 | .434 | 37.000 | .047 | 132.000 | .048 |
|  | *Median* | 5.730 | 0.057 | .051 | 1.000 | 4.196 | .100 | 3.033 | .128 |
| Random Coils (α) | *Distribution* | 22.590 | **<0.001*** | 85.000 | .687 | 149.000 | .000 | 8.000 | .000 |
|  | *Median* | 17.115 | **< 0.001*** | .051 | 1.000 | 18.132 | .000 | 20.571 | .000 |
|  | All tests conducted with N=25 samples and 1 degree of freedom  *Significance values based on Fisher exact 2-sided test  Significance level is set at 0.05 | | | | | | | | |

Table 11 Comparison of interstitial (In.Ar) and osteonal (On.Ar) bone differences in each environment. Red data are not considered reliable due to the heavily degraded amide spectra. Significant values are bolded and annotated by an asterisk*.

| **In.Ar Vs On.Ar** | | **Dry (historical)** | | **Wet (historical)** | | **Fossil (Geological)** | |
| --- | --- | --- | --- | --- | --- | --- | --- |
| **Diagenesis index** | | **Test statistic** | **p-value*** | **Test statistic** | **p-value*** | **Test statistic** | **p-value*** |
| C/P | *Distribution* | 68.000 | 0.652 | 144.000 | **0.035*** | 53.000 | 0.113 |
|  | *Median* | 0.182 | 1.000 | 5.143 | 0.057 | 3.846 | 0.115 |
| HPO4/P | *Distribution* | 48.000 | 0.545 | 153.000 | **0.011*** | 72.000 | 0.545 |
|  | *Median* | 1.636 | 0.395 | 5.143 | 0.57 | 0.154 | 1.000 |
| Crystallinity 1 (IRSF) | *Distribution* | 49.000 | 0.478 | 150.000 | **0.016*** | 85.000 | 0.980 |
|  | *Median* | 0.182 | 1.000 | 9.143 | **0.007*** | 0.154 | 1.000 |
| Crystallinity 2 | *Distribution* | 75.000 | 0.365 | 48.000 | **0.021*** | 110.000 | 0.204 |
|  | *Median* | 1.636 | 0.395 | 5.143 | 0.057 | 1.385 | 0.434 |
| AmI/P | *Distribution* | 73.000 | 0.438 | 140.000 | 0.056 | 62.000 | 0.264 |
|  | *Median* | 0.182 | 1.000 | 0.291 | 0.706 | 1.385 | 0.434 |
| AmII/P | *Distribution* | 85.000 | 0.116 | 114.000 | 0.482 | 81.000 | 0.462 |
|  | *Median* | 1.636 | 0.395 | 0.000 | 1.000 | 0.154 | 1.000 |
| AmI/ AmII | *Distribution* | 56.000 | 0.797 | 97.000 | 0.982 | 58.000 | 0.982 |
|  | *Median* | 0.182 | 1.000 | 0.000 | 1.000 | 1.385 | 0.434 |
| Collagen Integrity | *Distribution* | 55.000 | 0.718 | 93.000 | 0.839 | 43.000 | 0.262 |
|  | *Median* | 0.182 | 1.000 | 0.000 | 1.000 | 0.173 | 0.670 |
| Random Coils (α) | *Distribution* | 73.000 | 0.438 | 111.000 | 0.571 | 86.000 | 0.960 |
|  | *Median* | 0.182 | 1.000 | 0.571 | 0.706 | 0.154 | 1.000 |
| * Significance level set at 0.05 based on Fisher exact 2-sided test  - All tests conducted with N=23 samples,1 degree of freedom  - Independent Mann Whitney-U test used to compare distributions  - Independent Medians test (k samples) used to compare medians  - Organic diagenetic indices for the palaeontological condition should be viewed with caution due to the considerable degradation of Amide I and Amide II. | | | | | | | |

### 3 PCA comparisons of bone type and environments for s-FTIRM.

Diagenesis indices were compared together in PCA analyses to distinguish whole tissue (ROI) differences between environments (Section 3.2.1 and Section 3.2.2), and between tissue types for each depositional environment (fossil, dry, and wet). Model of fitness data (Table 12) and statistical comparison of groups between principal components (Tables 13-14) are first provided, followed by individual PCA comparisons. Each PCA compares a different suit of variables: : 1) all mineral and diagenetic variables, 2) all mineral and AmI/P, and 3) only mineral indices. Organic diagenesis for fossil specimens is not included except for AmI/P.

#### 3.1 Summary data

Table 12 Principal component analysis model fitness. Dark grey cells reflect conditions that were not tested based on the loss of organics in the fossil assemblage.

|  | **PCA: All Indices** | | | | | | | **PCA: Mineral Indices and AmI/P** | | | | | | **PCA: Mineral Indices only** | | | | | |
| --- | --- | --- | --- | --- | --- | --- | --- | --- | --- | --- | --- | --- | --- | --- | --- | --- | --- | --- | --- |
| **Condition** | **KMO** | **Sphericity** | **No. Components >1** | **Eigenvalue PC1** | **Eigenvalue PC2** | **Variance** | **KMO** | | **Sphericity Sig.** | **No. Components >1** | **Eigenvalue PC1** | **Eigenvalue PC2** | **Variance** | **KMO** | **Sphericity Sig.** | **No. Components** | **Eigenvalue PC1** | **Eigenvalue PC2** | **Variance** |
| *All burial Conditions: ROI* |  |  |  |  |  |  | 0.558 | | <0.001 | 2 | 3.08 | 1.06 | 82.65 | 0.451 | <0.001 | 2 | 2.31 | 1.01 | 82.95 |
| *Wet v Dry: ROI* | 0.541 | <0.001 | 3 | 4.41 | 1.63 | 67.09 | 0.587 | | <0.001 | 2 | 3.10 | 1.08 | 83.61 | 0.431 | <0.001 | 2 | 2.46 | 1.07 | 88.21 |
| *Fossil: In.Ar v. On.Ar* |  |  |  |  |  |  | 0.560 | | <0.01 | 2 | 2.34 | 1.44 | 75.56 | 0.558 | <0.001 | 2 | 2.31 | 1.02 | 83.25 |
| *Wet: In.Ar v. On.Ar* | 0.509 | <0.001 | 4 | 3.54 | 2.30 | 64.94 | **0.543** | | **<0.001** | **2** | **2.95** | **1.30** | **85.02** | 0.472 | <0.001 | 2 | 2.64 | 1.01 | 91.09 |
| *Dry: In.Ar v. On.Ar* | 0.658 | <0.01 | 2 | 4.66 | 2.54 | 80.01 | 0.661 | | <0.001 | 2 | 3.58 | 1.00 | 91.61 | 0.702 | <0.001 | 1 | 3.00 | 0.77 | 94.17 |

#### 3.1.1 Mann Whitney U results PCA of bone type assemblage comparing PC between All, and Wet and Dry environments.

Table 13 Summary table of significance comparing environments for whole region of interest (ROI) based on principal components derived from diagenetic indices. Dark grey cells reflect conditions that were not tested based on the loss of organics in the fossil assemblage. Significant values are bolded and annotated by an asterisk*.

|  | | **PCA: All Indices** | | | | **PCA: Mineral Indices and AmI/P** | | | | **PCA: Mineral Indices only** | | | |
| --- | --- | --- | --- | --- | --- | --- | --- | --- | --- | --- | --- | --- | --- |
| **Condition test*** | | **PC1** |  | **PC2** |  | **PC1** | | **PC2** | | **PC1** |  | **PC2** |  |
|  |  | *P* value | Test score | *P* value | Test score | P value | Test score | P value | Test score | P value | Test score | P value | Test score |
| All burial Conditions | *Median* |  |  |  |  | **<0.001*** | 21.014 | 0.078 | 5.108 | **<0.001*** | 21.014 | **0.010*** | 9.292 |
|  | *Distribution*** |  |  |  |  | **<0.001*** | 24.827 | 0.031 | 6.918 | **<0.001*** | 22.761 | **0.018*** | **7.933** |
| *Wet v Dry**** | *Median* |  |  |  |  | 1.000^a^ | 0.337 | 0.906^a^ | 1.066 | 0.085^a^ | 4.812 | 0.906^a^ | 1.066 |
|  | *Distribution* |  |  |  |  | 1.000^a^ | 2.701 | 0.943^a^ | -4.623 | 1.000^a^ | 4.156 | 0.660^a^ | -5.636 |
| *Wet v Fossil**** | *Median* |  |  |  |  | **0.000*^a^** | 20.571 | 1.000^a^ | 0.571 | **0.000*^a^** | 20.571 | 0.392^a^ | 2.286 |
|  | *Distribution* |  |  |  |  | **0.000*^a^** | -17.643 | 0.282^a^ | 7.214 | **0.001*^a^** | -16.000 | 0.282^a^ | 7.214 |
| *Dry v Fossil**** | *Median* |  |  |  |  | **0.000*^a^** | 21.280 | **0.024*^a^** | 6.997 | **0.000*^a^** | 14.490 | **0.024*^a^** | 6.997 |
|  | *Distribution* |  |  |  |  | **0.000* ^a^** | 20.334 | **0.030*^a^** | -11.838 | **0.000*^a^** | 20.156 | **0.015*^a^** | -12.851 |
| Wet v Dry | *Median* | 1.000^b^ | 0.051 | 0.695^b^ | 0.337 | 0.695^b^ | 0.337 | 0.111^b^ | 3.381 | 0.695^b^ | 0.337 | 0.111^b^ | 3.381 |
|  | *Distribution***** | 0.784 | 0.075 | 0.622 | 0.243 | 0.913 | 0.012 | 0.071 | 3.264 | 0.870 | 0.027 | 0.080 | 3.069 |

* Only PC1 and PC2 were tested.

** Independent-samples Kruskal-Wallis test for distributions

*** Pair-wise test of Kruskal-Wallis test

**** Independent-samples Mann Whitney-U test for distributions

NB: Asymptotic significance, at 95% confidence level, used unless stated otherwise

^a^ Adjusted by the Bonferroni correction for multiple tests

^b^ Fisher exact significance

#### 3.1.2 Mann Whitney U results PCA of environments comparing In.Ar and On.Ar.

*Table 14 Summary Table of significance between In.Ar and On.Ar bone type for each environment based on Principal components derived from diagenetic indices. Dark grey cells reflect conditions that were not tested based on the loss of organics in the fossil assemblage. Significant values are bolded and annotated by an asterisk*.*

|  | | **PCA: All Indices** | | | | **PCA: Mineral Indices and AmI/P** | | | | **PCA: Mineral Indices only** | | | |
| --- | --- | --- | --- | --- | --- | --- | --- | --- | --- | --- | --- | --- | --- |
|  |  | **PC1** | | **PC2** | | **PC1** | | **PC2** | | **PC1** | | **PC2** | |
| **Condition** | **Test** | **P value** | **Test score** | **P value** | **Test score** | **P value** | **Test score** | **P value** | **Test score** | **P value** | **Test score** | **P value** | **Test score** |
| Dry | *Median* | 0.086 | 4.545 | 0.395 | 1.636 | 0.395 | 1.636 | 1.000 | 0.182 | 1.000 | 0.182 | 1.000 | 0.182 |
|  | *Distribution* | 0.478 | 49.00 | 0.151 | 83.00 | 0.699 | 54.00 | 0.151 | 83.00 | 0.478 | 49.00 | 0.365 | 75.00 |
| Wet | *Median* | 0.057 | 5.143 | 0.057 | 5.143 | 0.057 | 5.143 | 0.257 | 2.286 | 0.057 | 5.143 | 0.706 | 0.571 |
|  | *Distribution* | **0.016*** | 150.00 | **0.031*** | 145.00 | **0.004*** | 159.00 | 0.541 | 84.00 | **0.005*** | 158.00 | 0.946 | 100.00 |
| Fossil | *Median* |  |  |  |  | 1.000 | 0.154 | 0.115 | 3.846 | 1.000 | 0.154 | 0.434 | 1.385 |
|  | *Distribution* |  |  |  |  | 0.687 | 76.00 | 0.091 | 51.00 | 0.511 | 71.00 | 0.125 | 54.00 |

NB: Fisher exact test used to determine significance at a 95% confidence level.

#### 3.2 PCA comparisons of ROI data between environments.

#### 3.2.1 ROI comparison of all depositional environments

Note: fossil depositional environment is also referred to as “palaeo” or “palaeontological” in graphs.

##### Mineral indices and AmI/P

| **Total Variance Explained^a^** | | | | | | |
| --- | --- | --- | --- | --- | --- | --- |
| *Component* | ***Initial Eigenvalues*** | | | ***Extraction Sums of Squared Loadings*** | | |
|  | *Total* | *% of Variance* | *Cumulative %* | *Total* | *% of Variance* | *Cumulative %* |
| **1** | 3.078 | 61.557 | 61.557 | 3.078 | 61.557 | 61.557 |
| **2** | 1.055 | 21.096 | 82.653 | 1.055 | 21.096 | 82.653 |
| **3** | .622 | 12.433 | 95.086 |  |  |  |
| **4** | .198 | 3.957 | 99.043 |  |  |  |
| **5** | .048 | .957 | 100.000 |  |  |  |
| Extraction Method: Principal Component Analysis. | | | | | | |
| a. Dataset = ROI | | | | | | |

| **Component Matrix^a,b^** | | |
| --- | --- | --- |
|  | **Component** | |
|  | PC1 | PC2 |
| CI1 | -.489 | .804 |
| AM1/P | .915 | -.186 |
| C/P | .701 | -.195 |
| CI2 | -.876 | -.332 |
| HPO4/P | .862 | .475 |
| Extraction Method: Principal Component Analysis. | | |
| a. Dataset = ROI | | |
| b. 2 components extracted. | | |

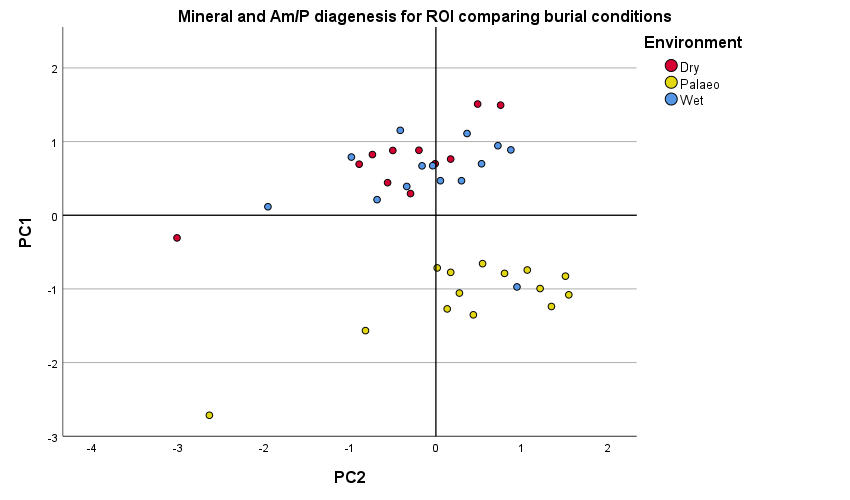

##### Mineral indices only

| **Total Variance Explained^a^** | | | | | | |
| --- | --- | --- | --- | --- | --- | --- |
| *Component* | **Initial Eigenvalues** | | | **Extraction Sums of Squared Loadings** | | |
|  | *Total* | *% of Variance* | *Cumulative %* | *Total* | *% of Variance* | *Cumulative %* |
| **1** | 2.309 | 57.726 | 57.726 | 2.309 | 57.726 | 57.726 |
| **2** | 1.009 | 25.224 | 82.950 | 1.009 | 25.224 | 82.950 |
| **3** | .620 | 15.497 | 98.447 |  |  |  |
| **4** | .062 | 1.553 | 100.000 |  |  |  |
| Extraction Method: Principal Component Analysis. | | | | | | |
| a. Dataset = ROI | | | | | | |

| **Component Matrix^a,b^** | | |
| --- | --- | --- |
|  | Component | |
|  | PC1 | PC2 |
| CI1 | -.396 | .854 |
| C/P | .694 | -.297 |
| CI2 | -.923 | -.220 |
| HPO4/P | .906 | .377 |
| Extraction Method: Principal Component Analysis. | | |
| a. Dataset = ROI | | |
| b. 2 components extracted. | | |

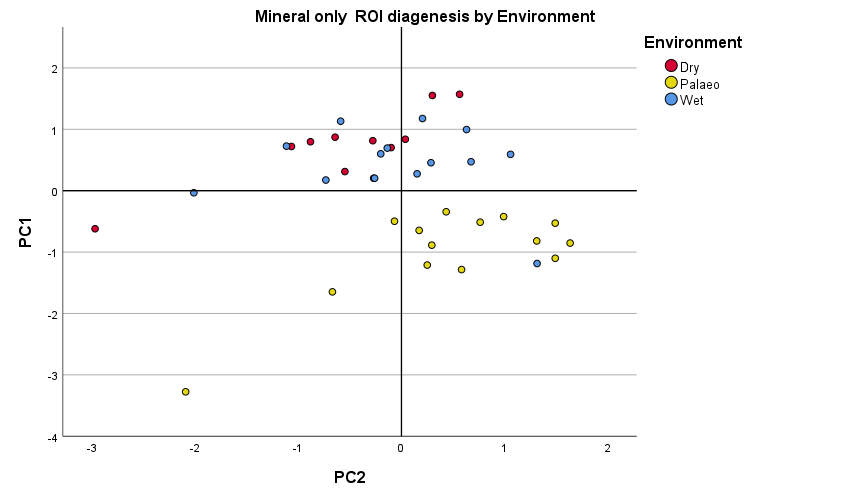

#### 3.2.2 ROI assemblages between wet and dry environments.

##### All indices

| **Total Variance Explained^a^** | | | | | | |
| --- | --- | --- | --- | --- | --- | --- |
| *Component* | **Initial Eigenvalues** | | | **Extraction Sums of Squared Loadings** | | |
|  | *Total* | *% of Variance* | *Cumulative %* | *Total* | *% of Variance* | *Cumulative %* |
| 1 | 4.406 | 48.958 | 48.958 | 4.406 | 48.958 | 48.958 |
| 2 | 1.632 | 18.129 | 67.086 | 1.632 | 18.129 | 67.086 |
| 3 | 1.301 | 14.455 | 81.541 | 1.301 | 14.455 | 81.541 |
| 4 | .808 | 8.977 | 90.518 |  |  |  |
| 5 | .436 | 4.849 | 95.367 |  |  |  |
| 6 | .245 | 2.726 | 98.093 |  |  |  |
| 7 | .122 | 1.357 | 99.450 |  |  |  |
| 8 | .040 | .446 | 99.896 |  |  |  |
| 9 | .009 | .104 | 100.000 |  |  |  |
| Extraction Method: Principal Component Analysis. | | | | | | |
| a. Dataset = ROI | | | | | | |

3 Eigenvalues >1. PC3 not included in scatter plot.

| **Component Matrix^a,b^** | | | |
| --- | --- | --- | --- |
|  | **Component** | | |
|  | PC1 | PC2 | PC3 |
| CI1 | .385 | .801 | .063 |
| Ami/P | .865 | .251 | .173 |
| C/P | .695 | -.350 | .461 |
| CI2 | -.864 | .039 | .326 |
| HPO4/P | .870 | .372 | -.094 |
| AmI/2 | .197 | .248 | -.782 |
| AM2/PO4 | .810 | .015 | .438 |
| Collagen cross links | .737 | -.357 | -.253 |
| Alpha helix | -.535 | .690 | .270 |
| Extraction Method: Principal Component Analysis. | | | |
| a. Dataset = ROI | | | |
| b. 3 components extracted. | | | |

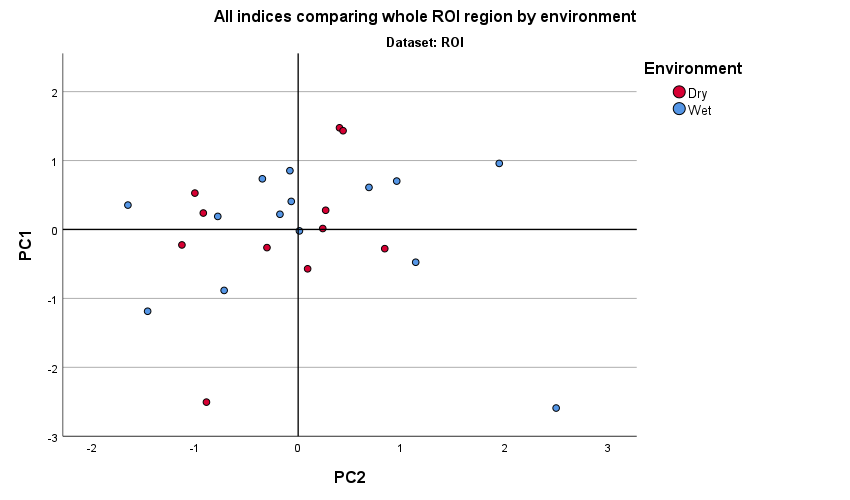

##### Mineral indices and AmI/P

| **Total Variance Explained^a^** | | | | | | |
| --- | --- | --- | --- | --- | --- | --- |
| *Component* | **Initial Eigenvalues** | | | **Extraction Sums of Squared Loadings** | | |
|  | *Total* | *% of Variance* | *Cumulative %* | *Total* | *% of Variance* | *Cumulative %* |
| **1** | 3.101 | 62.029 | 62.029 | 3.101 | 62.029 | 62.029 |
| **2** | 1.079 | 21.577 | 83.606 | 1.079 | 21.577 | 83.606 |
| **3** | .469 | 9.386 | 92.992 |  |  |  |
| **4** | .289 | 5.773 | 98.764 |  |  |  |
| **5** | .062 | 1.236 | 100.000 |  |  |  |
| Extraction Method: Principal Component Analysis. | | | | | | |
| a. Dataset = ROI | | | | | | |

| **Component Matrix^a,b^** | | |
| --- | --- | --- |
|  | Component | |
|  | PC1 | PC2 |
| CI1 | .594 | .752 |
| AmI/P | .853 | -.100 |
| C/P | .630 | -.683 |
| CI2 | -.839 | .105 |
| HPO4/P | .959 | .164 |
| Extraction Method: Principal Component Analysis. | | |
| a. Dataset = ROI | | |
| b. 2 components extracted. | | |

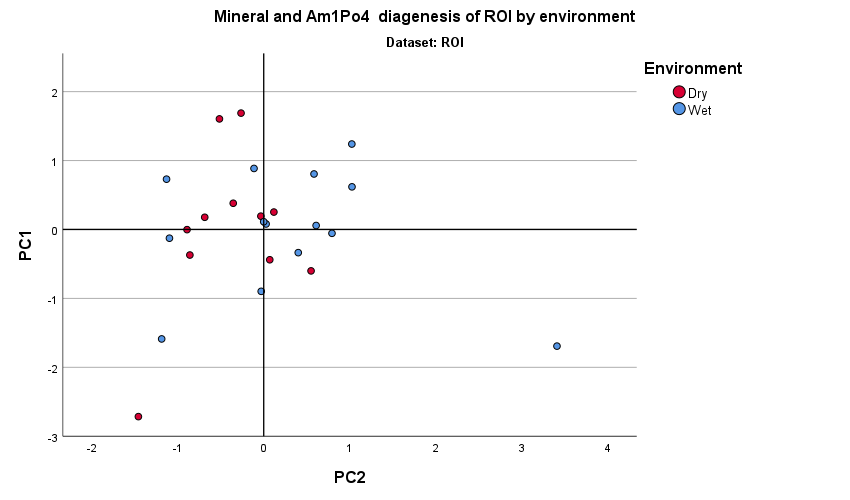

##### Mineral indices only

| **Total Variance Explained^a^** | | | | | | |
| --- | --- | --- | --- | --- | --- | --- |
| *Component* | **Initial Eigenvalues** | | | **Extraction Sums of Squared Loadings** | | |
|  | *Total* | *% of Variance* | *Cumulative %* | *Total* | *% of Variance* | *Cumulative %* |
| 1 | 2.459 | 61.480 | 61.480 | 2.459 | 61.480 | 61.480 |
| 2 | 1.069 | 26.725 | 88.205 | 1.069 | 26.725 | 88.205 |
| 3 | .406 | 10.147 | 98.352 |  |  |  |
| 4 | .066 | 1.648 | 100.000 |  |  |  |
| Extraction Method: Principal Component Analysis. | | | | | | |
| a. Dataset = ROI | | | | | | |

| **Component Matrix^a,b^** | | |
| --- | --- | --- |
|  | Component | |
|  | PC1 | PC2 |
| CI1 | .626 | .725 |
| C/P | .596 | -.709 |
| CI2 | -.874 | .165 |
| HPO4/P | .973 | .115 |
| Extraction Method: Principal Component Analysis. | | |
| a. Dataset = ROI | | |
| b. 2 components extracted. | | |

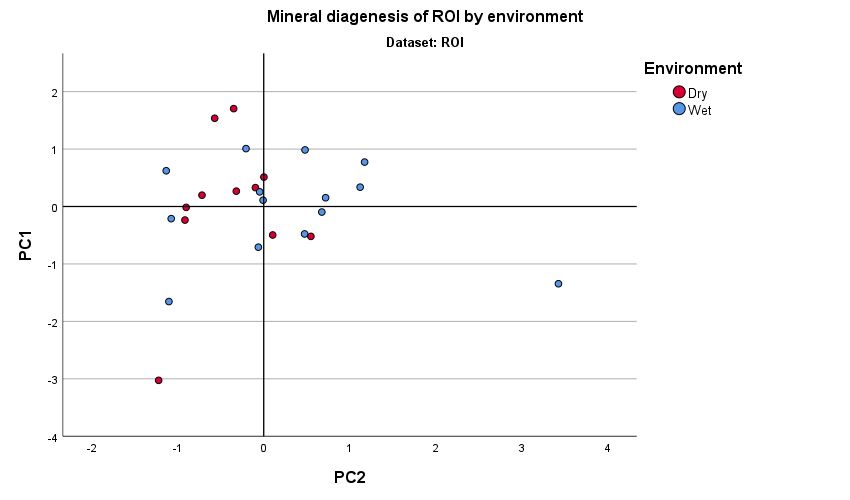

#### 3.3 PCA of environments comparing In.Ar and On.Ar

#### 3.3.1 Fossil depositional environment

##### Mineral indices and AmI/P

| **Total Variance Explained** | | | | | | |
| --- | --- | --- | --- | --- | --- | --- |
| *Component* | **Initial Eigenvalues** | | | **Extraction Sums of Squared Loadings** | | |
|  | *Total* | *% of Variance* | *Cumulative %* | *Total* | *% of Variance* | *Cumulative %* |
| 1 | 2.335 | 46.695 | 46.695 | 2.335 | 46.695 | 46.695 |
| 2 | 1.443 | 28.863 | 75.558 | 1.443 | 28.863 | 75.558 |
| 3 | .596 | 11.915 | 87.473 |  |  |  |
| 4 | .528 | 10.569 | 98.043 |  |  |  |
| 5 | .098 | 1.957 | 100.000 |  |  |  |
| Extraction Method: Principal Component Analysis. | | | | | | |

| **Component Matrix^a^** | | |
| --- | --- | --- |
|  | Component | |
|  | PC1 | PC2 |
| CI1 | .767 | -.151 |
| AmI/P | -.213 | .820 |
| C/P | .066 | .833 |
| CI2 | -.884 | -.229 |
| HPO4/P | .956 | .034 |
| Extraction Method: Principal Component Analysis. | | |
| a. 2 components extracted. | | |

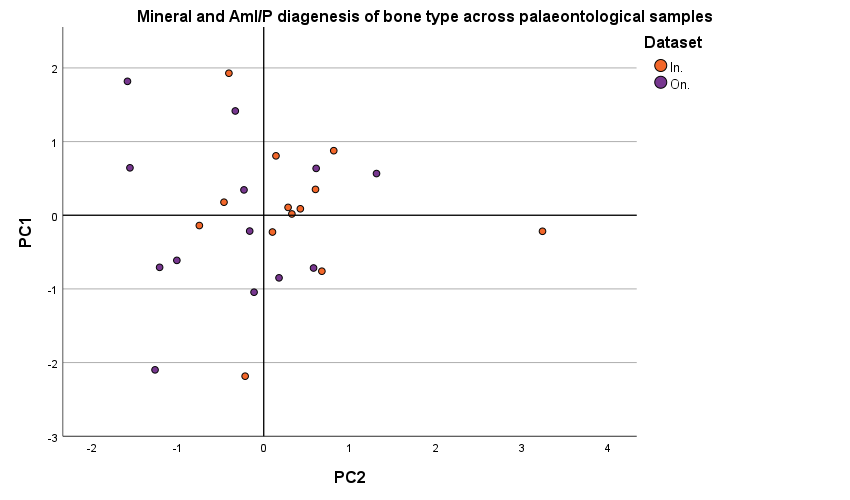

##### Mineral indices only

| **Total Variance Explained** | | | | | | |
| --- | --- | --- | --- | --- | --- | --- |
| *Component* | **Initial Eigenvalues** | | | **Extraction Sums of Squared Loadings** | | |
|  | *Total* | *% of Variance* | *Cumulative %* | *Total* | *% of Variance* | *Cumulative %* |
| 1 | 2.311 | 57.784 | 57.784 | 2.311 | 57.784 | 57.784 |
| 2 | 1.019 | 25.466 | 83.249 | 1.019 | 25.466 | 83.249 |
| 3 | .566 | 14.150 | 97.399 |  |  |  |
| 4 | .104 | 2.601 | 100.000 |  |  |  |
| **Extraction Method: Principal Component Analysis.** | | | | | | |

| **Component Matrix^a^** | | |
| --- | --- | --- |
|  | Component | |
|  | PC1 | PC2 |
| CI1 | .752 | -.234 |
| C/P | .135 | .975 |
| CI2 | -.903 | -.105 |
| HPO4/P | .955 | -.052 |
| Extraction Method: Principal Component Analysis. | | |
| a. 2 components extracted. | | |

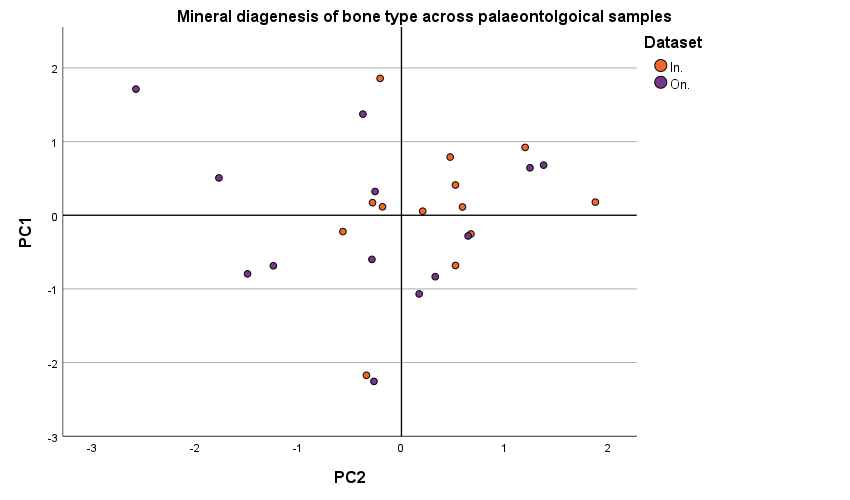

#### 3.3.2 Dry depositional environment

##### All indices

| **Total Variance Explained** | | | | | | |
| --- | --- | --- | --- | --- | --- | --- |
| *Component* | **Initial Eigenvalues** | | | **Extraction Sums of Squared Loadings** | | |
|  | *Total* | *% of Variance* | *Cumulative %* | *Total* | *% of Variance* | *Cumulative %* |
| 1 | 4.660 | 51.782 | 51.782 | 4.660 | 51.782 | 51.782 |
| 2 | 2.541 | 28.231 | 80.013 | 2.541 | 28.231 | 80.013 |
| 3 | .913 | 10.142 | 90.155 |  |  |  |
| 4 | .327 | 3.636 | 93.790 |  |  |  |
| 5 | .264 | 2.933 | 96.723 |  |  |  |
| 6 | .164 | 1.821 | 98.544 |  |  |  |
| 7 | .093 | 1.032 | 99.576 |  |  |  |
| 8 | .025 | .280 | 99.856 |  |  |  |
| 9 | .013 | .144 | 100.000 |  |  |  |
| Extraction Method: Principal Component Analysis. | | | | | | |

| **Component Matrix^a^** | | |
| --- | --- | --- |
|  | Component | |
|  | PC1 | PC2 |
| CI1 | .876 | -.010 |
| AmI/P | .832 | .405 |
| C/P | .686 | .660 |
| CI2 | -.786 | .328 |
| HPO4/P | .926 | -.193 |
| Ami/II2 | -.067 | -.841 |
| AmII/P | .612 | .598 |
| Collagen cross links | .734 | -.486 |
| Alpha Helix | -.582 | .704 |
| Extraction Method: Principal Component Analysis. | | |
| a. 2 components extracted. | | |

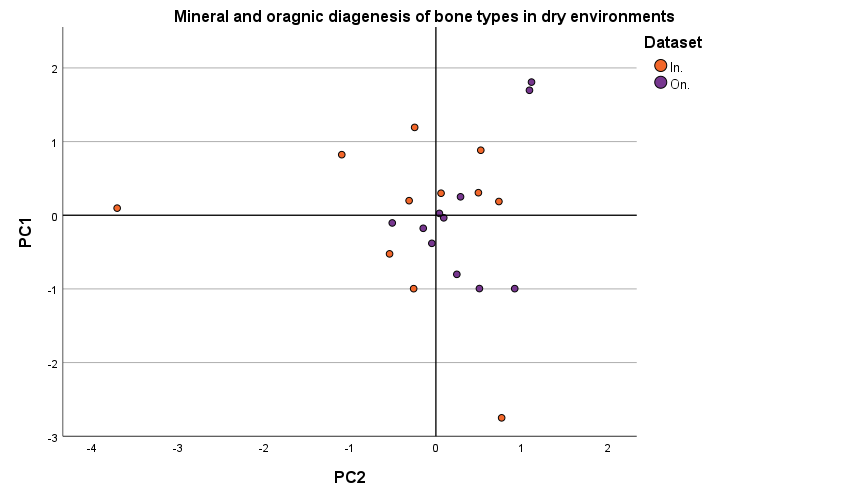

##### Mineral indices and AmI/P

| **Total Variance Explained** | | | | | | |
| --- | --- | --- | --- | --- | --- | --- |
| *Component* | **Initial Eigenvalues** | | | **Extraction Sums of Squared Loadings** | | |
|  | *Total* | *% of Variance* | *Cumulative %* | *Total* | *% of Variance* | *Cumulative %* |
| 1 | 3.579 | 71.573 | 71.573 | 3.579 | 71.573 | 71.573 |
| 2 | 1.002 | 20.034 | 91.607 | 1.002 | 20.034 | 91.607 |
| 3 | .231 | 4.630 | 96.237 |  |  |  |
| 4 | .151 | 3.015 | 99.252 |  |  |  |
| 5 | .037 | .748 | 100.000 |  |  |  |
| Extraction Method: Principal Component Analysis. | | | | | | |

| **Component Matrix^a^** | | |
| --- | --- | --- |
|  | Component | |
|  | PC1 | PC2 |
| CI1 | .935 | -.169 |
| AmI/P | .815 | .476 |
| C/P | .743 | .616 |
| CI2 | -.784 | .533 |
| HPO4/P | .935 | -.289 |
| Extraction Method: Principal Component Analysis. | | |
| a. 2 components extracted. | | |

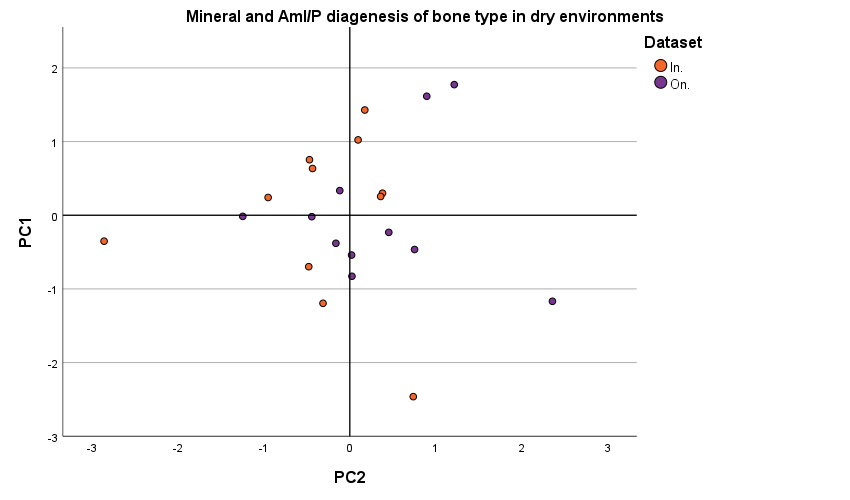

##### Mineral indices only

| **Total Variance Explained** | | | | | | |
| --- | --- | --- | --- | --- | --- | --- |
| *Component* | **Initial Eigenvalues** | | | **Extraction Sums of Squared Loadings** | | |
|  | *Total* | *% of Variance* | *Cumulative %* | *Total* | *% of Variance* | *Cumulative %* |
| 1 | 3.001 | 75.037 | 75.037 | 3.001 | 75.037 | 75.037 |
| 2 | .765 | 19.133 | 94.170 | .765 | 19.133 | 94.170 |
| 3 | .185 | 4.626 | 98.796 |  |  |  |
| 4 | .048 | 1.204 | 100.000 |  |  |  |
| Extraction Method: Principal Component Analysis. | | | | | | |

| **Component Matrix^a^** | | |
| --- | --- | --- |
|  | Component | |
|  | PC1 | PC2 |
| CI1 | .962 | .024 |
| C/P | .647 | .747 |
| CI2 | -.851 | .431 |
| HPO4/P | .967 | -.144 |
| Extraction Method: Principal Component Analysis. | | |
| a. 2 components extracted. | | |

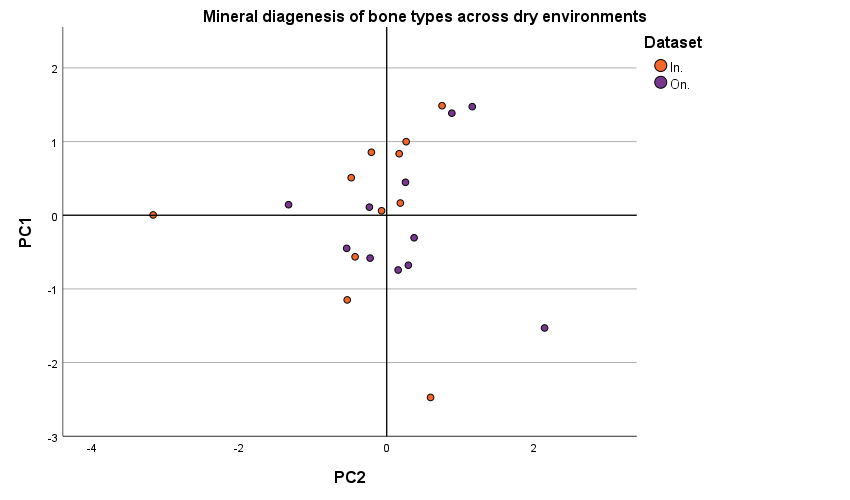

#### 3.3.3 Wet Environment

##### All indices

| **Total Variance Explained** | | | | | | | | | |
| --- | --- | --- | --- | --- | --- | --- | --- | --- | --- |
| *Component* | **Initial Eigenvalues** | | | | | | **Extraction Sums of Squared Loadings** | | |
|  | *Total* | | *% of Variance* | | *Cumulative %* | | *Total* | *% of Variance* | *Cumulative %* |
| 1 | 3.541 | | 39.346 | | 39.346 | | 3.541 | 39.346 | 39.346 |
| 2 | 2.303 | | 25.592 | | 64.937 | | 2.303 | 25.592 | 64.937 |
| 3 | 1.452 | | 16.138 | | 81.075 | |  |  |  |
| 4 | 1.133 | | 12.593 | | 93.668 | |  |  |  |
| 5 | .273 | | 3.031 | | 96.698 | |  |  |  |
| 6 | .163 | | 1.815 | | 98.513 | |  |  |  |
| 7 | .101 | | 1.126 | | 99.640 | |  |  |  |
| 8 | .024 | | .262 | | 99.902 | |  |  |  |
| 9 | .009 | | .098 | | 100.000 | |  |  |  |
| Extraction Method: Principal Component Analysis. | | | | | | | | | |
| **Component Matrix^a^** | | | | | |  |  |  |  |
|  | | Component | | | |  |  |  |  |
|  |  | PC1 | | PC2 | |  |  |  |  |
| CI1 | | .320 | | .848 | |  |  |  |  |
| AmI/P | | .825 | | -.138 | |  |  |  |  |
| C/P | | .772 | | -.346 | |  |  |  |  |
| CI2 | | -.822 | | -.461 | |  |  |  |  |
| HPO4/P | | .635 | | .742 | |  |  |  |  |
| Ami/II2 | | .096 | | .351 | |  |  |  |  |
| AmII/P | | .697 | | -.528 | |  |  |  |  |
| Collagen cross links | | .584 | | -.419 | |  |  |  |  |
| Alpha Helix | | -.495 | | .326 | |  |  |  |  |
| Extraction Method: Principal Component Analysis. | | | | | |  |  |  |  |
| a. 2 components extracted. | | | | | |  |  |  |  |

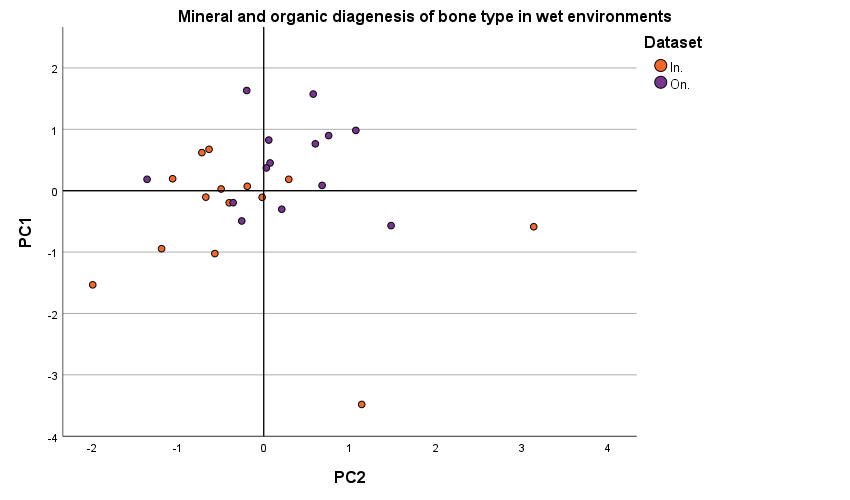

##### Mineral indices and AmI/P

| **Total Variance Explained** | | | | | | |
| --- | --- | --- | --- | --- | --- | --- |
| *Component* | **Initial Eigenvalues** | | | **Extraction Sums of Squared Loadings** | | |
|  | *Total* | *% of Variance* | *Cumulative %* | *Total* | *% of Variance* | *Cumulative %* |
| 1 | 2.947 | 58.933 | 58.933 | 2.947 | 58.933 | 58.933 |
| 2 | 1.304 | 26.090 | 85.022 | 1.304 | 26.090 | 85.022 |
| 3 | .423 | 8.463 | 93.485 |  |  |  |
| 4 | .294 | 5.885 | 99.370 |  |  |  |
| 5 | .031 | .630 | 100.000 |  |  |  |
| Extraction Method: Principal Component Analysis. | | | | | | |

| **Component Matrix^a^** | | |
| --- | --- | --- |
|  | Component | |
|  | PC1 | PC2 |
| CI1 | .731 | -.536 |
| AmI/P | .648 | .591 |
| C/P | .562 | .727 |
| CI2 | -.918 | .072 |
| HPO4/P | .912 | -.365 |
| Extraction Method: Principal Component Analysis. | | |
| a. 2 components extracted. | | |

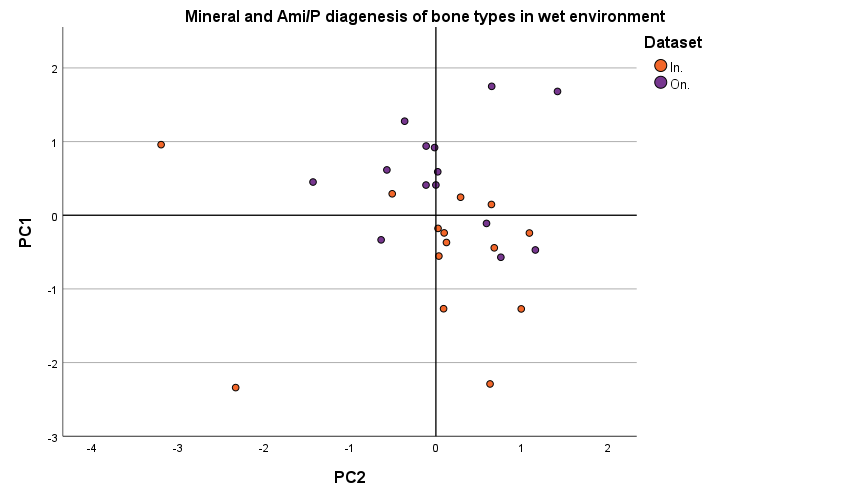

##### Mineral indices only

| **Total Variance Explained** | | | | | | |
| --- | --- | --- | --- | --- | --- | --- |
| *Component* | **Initial Eigenvalues** | | | **Extraction Sums of Squared Loadings** | | |
|  | *Total* | *% of Variance* | *Cumulative %* | *Total* | *% of Variance* | *Cumulative %* |
| 1 | 2.639 | 65.963 | 65.963 | 2.639 | 65.963 | 65.963 |
| 2 | 1.005 | 25.123 | 91.085 | 1.005 | 25.123 | 91.085 |
| 3 | .325 | 8.126 | 99.211 |  |  |  |
| 4 | .032 | .789 | 100.000 |  |  |  |
| Extraction Method: Principal Component Analysis. | | | | | | |

| **Component Matrix^a^** | | |
| --- | --- | --- |
|  | Component | |
|  | PC1 | PC2 |
| CI1 | .808 | -.437 |
| C/P | .431 | .872 |
| CI2 | -.928 | -.154 |
| HPO4/P | .969 | -.171 |
| Extraction Method: Principal Component Analysis. | | |
| a. 2 components extracted. | | |

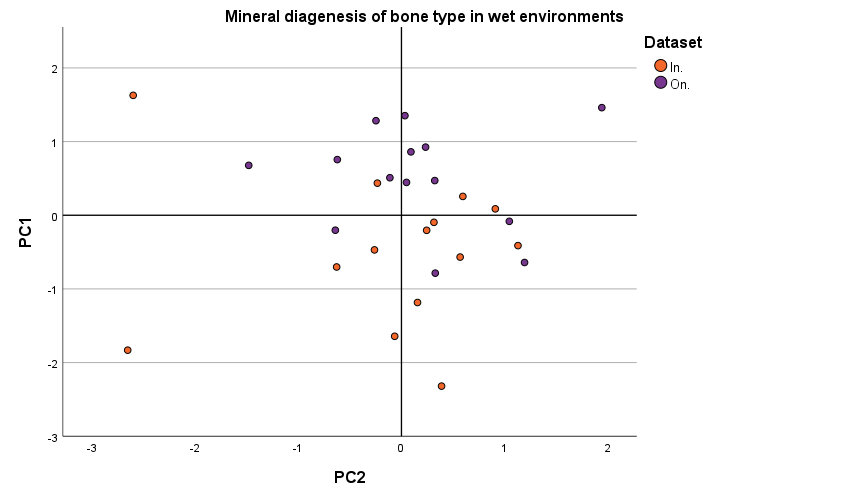
